# Serum triglycerides in Alzheimer’s disease: Relation to neuroimaging and CSF biomarkers

**DOI:** 10.1101/441394

**Authors:** Megan M. Bernath, Sudeepa Bhattacharyya, Kwangsik Nho, Dinesh Kumar Barupal, Oliver Fiehn, Rebecca Baillie, SL Risacher, Matthias Arnold, Tanner Jacobson, John Q. Trojanowski, Leslie M. Shaw, Michael W. Weiner, P. Murali Doraiswamy, Rima Kaddurah-Daouk, Andrew J. Saykin, for the Alzheimer’s Disease Neuroimaging Initiative, Alzheimer’s Disease Metabolomics Consortium

## Abstract

**Objective:** To investigate the association of triglyceride (TG) principal component scores with Alzheimer’s disease (AD) and the “A/T/N/V” (Amyloid, Tau, Neurodegeneration, and Cerebrovascular disease) biomarkers for AD.

**Methods:** Serum levels of 84 TG species were measured using untargeted lipid profiling of 689 participants from the Alzheimer’s Disease Neuroimaging Initiative (ADNI) cohort including 190 cognitively normal older adults (CN) and 339 mild cognitive impairment (MCI) and 160 AD. Principal component analysis with factor rotation was used for dimension reduction of TG species. Differences in principal components between diagnostic groups and associations between principal components and AD biomarkers (including CSF, MRI and [18F]FDG-PET) were assessed using a multivariate generalized linear model (GLM) approach. In both cases, the Bonferroni method of adjustment was employed to correct for multiple comparisons.

**Results:** The 84 TGs yielded 9 principal components, two of which consisting of long-chain, polyunsaturated fatty acid-containing TGs (PUTGs), were significantly associated with MCI and AD. Lower levels of PUTGs were observed in MCI and AD compared to CN. PUTG principal component scores were also significantly associated with hippocampal volume and entorhinal cortical thickness. In participants carrying *APOE* ε4 allele, these principal components were significantly associated with CSF amyloid-β_1-42_ values and entorhinal cortical thickness.

**Conclusions:** This study shows PUTG component scores significantly associated with diagnostic group and AD biomarkers, a finding that was more pronounced in *APOE* ε4 carriers. Replication in independent larger studies and longitudinal follow-up are warranted.

## Introduction

Triglycerides (TGs) may represent a risk factor for Alzheimer’s disease (AD), yet this relationship is not well understood. TGs are lipids that are comprised of three fatty acids (FA). Blood total TG levels are measured in routine clinical check-ups. Depending on the FAs that contribute to the TG, many combinations of carbon chains and double bonds result, leading to more than 6,000 species encompassed by the term “triglyceride”. Conflicting reports exist in the literature regarding TG homeostasis in AD. No relationship between AD and total TGs has been reported,^1^ while others suggest that elevated TGs early in life represented a risk factor for increased amyloidosis 20 years later.^2,3^ Finally, others found that individuals with probable AD had significantly decreased serum TG levels.^4,5^

Apolipoprotein E (*APOE*) ε4 carrier status may serve as a risk factor for altered TG levels in AD. TGs are transported by lipoproteins, due to their lipophilic nature. APOE regulates TG homeostasis by acting as a ligand for the TG-rich lipoproteins. The *APOE* ε4 allele is a major genetic risk factor for sporadic AD and is associated with a significant decrease in blood TG levels in AD.^5^ Here, we examined the association between TGs and AD biomarkers, with and without medication adjustment. We also investigated the effect of *APOE* ε4 by stratifying on *APOE* ε4 carrier status. This study shows that serum-based TG principal components differ as a function of diagnostic group and are associated with AD biomarkers.

## Methods

### Study sample

All individuals used in this study were participants from the Alzheimer’s Disease Neuroimaging Initiative Phase 1 (ADNI-1), which is a longitudinal study aimed to explore clinical, genetic, imaging, and biological biomarkers for early AD progression. ADNI-1 is a multicenter consortium across 59 sites in the U.S. and Canada, composed of approximately 200 cognitively normal (CN) older adults, 400 adults diagnosed with mild cognitive impairment (MCI) and 200 adults diagnosed with probable Alzheimer’s disease (AD), ranging in age from 47 to 91 years old.^6,7^ In partnership with ADNI, the Alzheimer’s Disease Metabolomics Consortium (ADMC) provided serum metabolic data for this group of participants. Demographic information, *APOE*, clinical information, neuroimaging and CSF biomarker data were downloaded from the ADNI data repository (http://adni.loni.usc.edu).

### Lipid analysis

Lipid analyses were performed as previously described.^8^ In summary, an untargeted lipidomics dataset was generated by the NIH West Coast Metabolomics Center (http://metabolomics.ucdavis.edu) using an ultra-high performance liquid chromatography-quadrupole time-of-flight (UHPLC-QTOF) mass spectrometer (Agilent, Santa Clara, California) for 807 baseline serum samples from ADNI-1 participants of the ADMC initiative. Lipid species were annotated by matching accurate mass, isotope abundance, retention times and MS/MS fragmentation spectra in LipidBlast^9^ in-silico mass spectral library and measured by signal intensities on the precursor mass level. After data processing, the quality of data was assessed and low-quality lipid species were removed. These quality control (QC) analyses resulted 349 annotated lipids, including 84 triglyceride species. The TG values obtained from the QC step were unadjusted and adjusted^10^ for the effect of medication use at baseline.

### Neuroimaging analysis

MRI scans were processed prior to download as previously described.^11,12^ These scans were further processed locally using FreeSurfer version 5.1.^13–15^ Regions of interest were extracted, including the bilateral hippocampal volumes, entorhinal cortical thickness, and total intracranial volume (ICV). [^18^F]fluorodeoxyglucose (FDG) PET scans were pre-processed prior to download.^13,16^ Standardized uptake value ratio (SUVR) images were created by intensity-normalization using a pons reference region. Mean SUVR values were extracted for each participant from an overall cortical ROI representing regions where CN>AD from the full ADNI-1 cohort. White matter hyperintensity volumes (WMHI) were assessed using previously described methods.^17,18^

### CSF biomarker analysis

The CSF biomarker data were downloaded from the ADNI data repository. As previously described,^19^ CSF measurements for amyloid β 1-42 peptide (Aβ_1-42_), total tau (t-tau), and tau phosphorylated at threonine 181 (p-tau_181P_) were obtained by the validated and highly automated Roche Elecsys electrochemiluminescence immunoassays.

### Cognitive Assessment

We used the modified Alzheimer’s Disease Assessment Scale-cognition sub-scale (ADAS-Cog 13)^20^ and memory (ADNI-MEM) as indices of general cognitive performance.^21,22^ ADAS-Cog includes eleven items, assessing memory, language, praxis, and orientation. ADAS-Cog 13 includes all items from ADAS-Cog, in addition to delayed recall and cancellation tasks. ADNI-MEM is a memory composite score calculated from the items in several memory tasks including the Rey Auditory Verbal Learning Test (RAVLT), ADAS-Cog, Logical Memory (passage recall), and the Mini-Mental Status Exam (MMSE). Alternate forms were accounted for where applicable.^21^

### A/T/N/V biomarkers

We used CSF Aβ_1-42_ levels as a biomarker of amyloid-β (“A”), CSF p-tau levels as a biomarker of tau (“T”), structural atrophy on MRI, FDG PET metabolism, and CSF t-tau levels as biomarkers of neurodegeneration (“N”), and white matter hyperintensity volume (WMHI) as a biomarker for microvascular disease burden (“V”), as described in the National Institute of Aging-Alzheimer’s Association (NIA-AA) Research Framework.^23^ Some ADNI participants had missing data for specific A/T/N/V biomarkers. The specific N available for each biomarker is presented in Table 1.

**Table 1.** Demographics of ADNI participants included in the analysis by diagnosis.

|  | N | CN (N=190) | N | MCI (N=339) | N | AD (N=160) |
| --- | --- | --- | --- | --- | --- | --- |
| Age | 190 | 75.29 (4.93) | 339 | 74.25 (7.54) | 160 | 74.43 (7.32) |
| Sex (Male/Female) | 190 | 94/96 | 339 | 218/121 | 160 | 79/81 |
| Education | 190 | 15.96 (2.97) | 339 | 15.63 (3.027) | 160 | 14.56 (3.093) |
| *APOE* ε4 status<br>(presence/absence) | 190 | 52/138 | 339 | 177/162 | 160 | 108/52 |
| BMI | 190 | 26.72 (4.55) | 339 | 26.15 (3.99) | 160 | 25.57 (4.00) |
| Total triglycerides | 190 | 140.27 (79.77) | 339 | 154.41 (142.53) | 160 | 156.83<br>(99.091) |
| Hippocampal volume | 189 | 3483.79<br>(445.30) | 339 | 3086.87<br>(528.88) | 159 | 2758.67<br>(502.71) |
| Entorhinal Thickness | 189 | 3.41 (0.29) | 339 | 3.11 (0.44) | 159 | 2.76 (0.45) |
| FDG Global Cortex | 77 | 1.44 (4.095) | 171 | 1.38 (0.15) | 77 | 1.25 (0.13) |
| WMHI | 186 | -0.68 (0.70) | 338 | -0.62 (0.73) | 159 | -0.44 (0.67) |
| CSF Aβ | 88 | 3.036 (0.23) | 168 | 2.88 (0.22) | 84 | 2.77 (0.20) |
| CSF t-tau | 86 | 2.37 (0.15) | 168 | 2.46 (0.17) | 83 | 2.52 (0.16) |
| CSF p-tau | 86 | 1.33 (0.16) | 168 | 1.44 (0.20) | 83 | 1.51 (0.19) |
| ADAS-13 | 190 | 9.16 (4.10) | 336 | 18.56 (6.20) | 156 | 28.72 (7.96) |
| ADNI-MEM | 190 | 1.014 (0.54) | 339 | -0.099 (0.60) | 160 | -0.84 (0.54) |

Data are reported as mean (standard deviation) unless otherwise indicated.
*Abbreviations:* CN: cognitively normal older adult controls; MCI: mild cognitive impairment;
AD: Alzheimer's disease; BMI: body mass index; WMHI: white matter hyperintensity volume;
FDG Global cortex: cortical glucose SUVR measured from [ $^{18}\text{F}$ ]FDG PET scans; CSF: cerebrospinal fluid; CSF Aβ : CSF amyloid $\beta$ 1-42 peptide (A $\beta_{1-42}$ ); CSF p-tau: CSF tau phosphorylated at threonine 181 (CSF p-tau<sub>181P</sub>); CSF t-tau: CSF total tau; ADAS-Cog 13: Alzheimer's Disease Assessment Scale-cognition sub-scale.

### Medication adjustment

Triglyceride data was corrected for 41 major medication classes used to treat psychiatric (including different categories of benzodiazepines, antipsychotics, and antidepressants) and cardiovascular conditions (including different categories of antihypertensives, cholesterol treatment, and antidiabetics), as well as dietary supplements (Co-Q10, fish oil, nicotinic acid, and acetyl L-carnitine). For complete medication list and procedures, please see previously reported methods.^10^

### Statistical analysis

Dimension reduction was performed on the original 84 triglycerides using Principal Component Analysis (PCA) as implemented in SPSS (v 24). Initial principal component (PC) extraction was followed by orthogonal rotation to yield principal component-based factor scores. Number of principal components extracted was pre-specified using the standard eigenvalues greater than 1 criterion. “Top contributors” of each rotated principal component were defined as those with a factor loading ≥ 0.8. All other contributors for each rotated principal component (with a factor loading < 0.8) were excluded from further consideration. A multivariate generalized linear model (GLM) was performed to assess between diagnosis group (CN, MCI, AD) differences across all principal components. A multivariate GLM was used to assess the association of A/T/N/V biomarkers with “PC3” and “PC5”. Potential covariates of interest were screened for possible inclusion in analyses of the PCA results including age, sex, body mass index (BMI), total triglycerides, and APOE ε4 status using linear regression. Years of education for cognitive performance, and years of education and intracranial volume (ICV) for MRI biomarkers were added as additional covariates. For example, statistical models for “PC5” and diagnosis (a), CSF Aβ_1-42_ (b), hippocampal volume (c), and APOE ε4 with entorhinal thickness interaction (d) respectively, appeared as follows:

a. Dependent variable (DV)= “PC5”; independent variable (IV)=Diagnosis (CN, MCI, AD); Covariates=age, sex, BMI, total triglycerides, APOE ε4 carrier status;
b. DV= “PC5”; IV= CSF Aβ_1-42_; Covariates= age, sex, BMI, total triglycerides, APOE ε4 carrier status;
c. DV= “PC5”; IV= Hippocampal volume; Covariates= age, sex, BMI, total triglycerides, APOE ε4 carrier status, intracranial volume (ICV), years of education
d. DV= “PC5”; IV= Entorhinal thickness*APOE ε4 carrier status; Covariates= age, sex, BMI, total triglycerides, intracranial volume (ICV), years of education, APOE ε4 carrier status, entorhinal thickness

Additional covariates assessed for cognitive performance (ADNI-MEM, ADAS-Cog 13) analyses included: years of education. We did not use total serum cholesterol levels as a covariate because total serum cholesterol levels had no effect on all 6 principal components. We did not use systolic and diastolic blood pressures as covariates because neither variable differed as a function of diagnosis or was associated with A/T/N/V biomarkers in our dataset. Significant associations were defined as p<0.05 after Bonferroni adjustment to correct for multiple testing.

### Whole brain surface-based analysis

As previously described, we performed a multivariate analysis of cortical thickness over the whole brain using SurfStat software (http://www.math.mcgill.ca/keith/surfstat/). We constructed a GLM using age, sex, *APOE* ε4 status, BMI, total triglycerides, years of education, and ICV as covariates. We corrected for multiple comparisons using the random field theory (RFT) correction method at p < 0.05 significance level. ^24^

### Data Availability Statement

All data used in the analyses reported here are available in the ADNI data repository (http://adni.loni.usc.edu).

### Standard Protocol Approvals, Registrations, and Patient Consents

Written informed consent was obtained at the time of enrollment for imaging and genetic sample collection and protocols of consent forms were approved by each participating sites’ Institutional Review Board (IRB).

## Results

In the analysis, we included 689 ADNI participants who had baseline data for the 84 TGs (190 cognitively normal older adults (CN), 339 mild cognitive impairment (MCI) and 160 AD) after quality control procedures including removing participants with non-fasting status (n=69). Demographic information is shown in Table 1.

### Principal component analysis (PCA) for dimension reduction of TGs

Dimension reduction with PCA resulted in 9 principal components with eigenvalues >1 (**Table e-1**). After selecting for the “top contributors” with a factor loading ≥ 0.8 in each component, 6 of 9 components remained for further analysis (**Table e-2**).

### Between group differences in TG principal components

Figure 1 illustrates the profile of group differences between CN and MCI and AD for principal components (“PC3” and “PC5”) after adjusting for multiple testing and covariates. We identified significant group differences in “PC5” between CN and AD (*p*-value= 4.32E-04, Cohen’s d= 0.386) and between CN and MCI (*p*-value= 1.84E-03, Cohen’s d= 0.313; Figure 1). MCI and AD did not differ significantly for either “PC3” or “PC5.” “PC5” consists of six long-chain, polyunsaturated TGs (PUTGs), with all species containing 8 or more double bonds. Lower levels of PUTGs belonging to “PC5” are seen in MCI and AD compared to CN (**Figure e-1; Table e-3**). In addition, among the six PUTGs, four (TG 60:11, TG 58:9, TG 58:8, and TG 56:8) were significantly lower in MCI and AD compared to CN (*p*-value < 0.05) (**Figure e-1**). We also identified suggestive group differences in“PC3” between CN and AD (*p*-value=0.0533, Cohen’s d= 0.179; Figure 1). “PC3” consisted of seven PUTGs, with almost all species containing 2 or more double bonds. Similar to “PC5”, AD patients compared to CN showed lower component scores for “PC3” (**Table e-4**).

**Figure 1:**
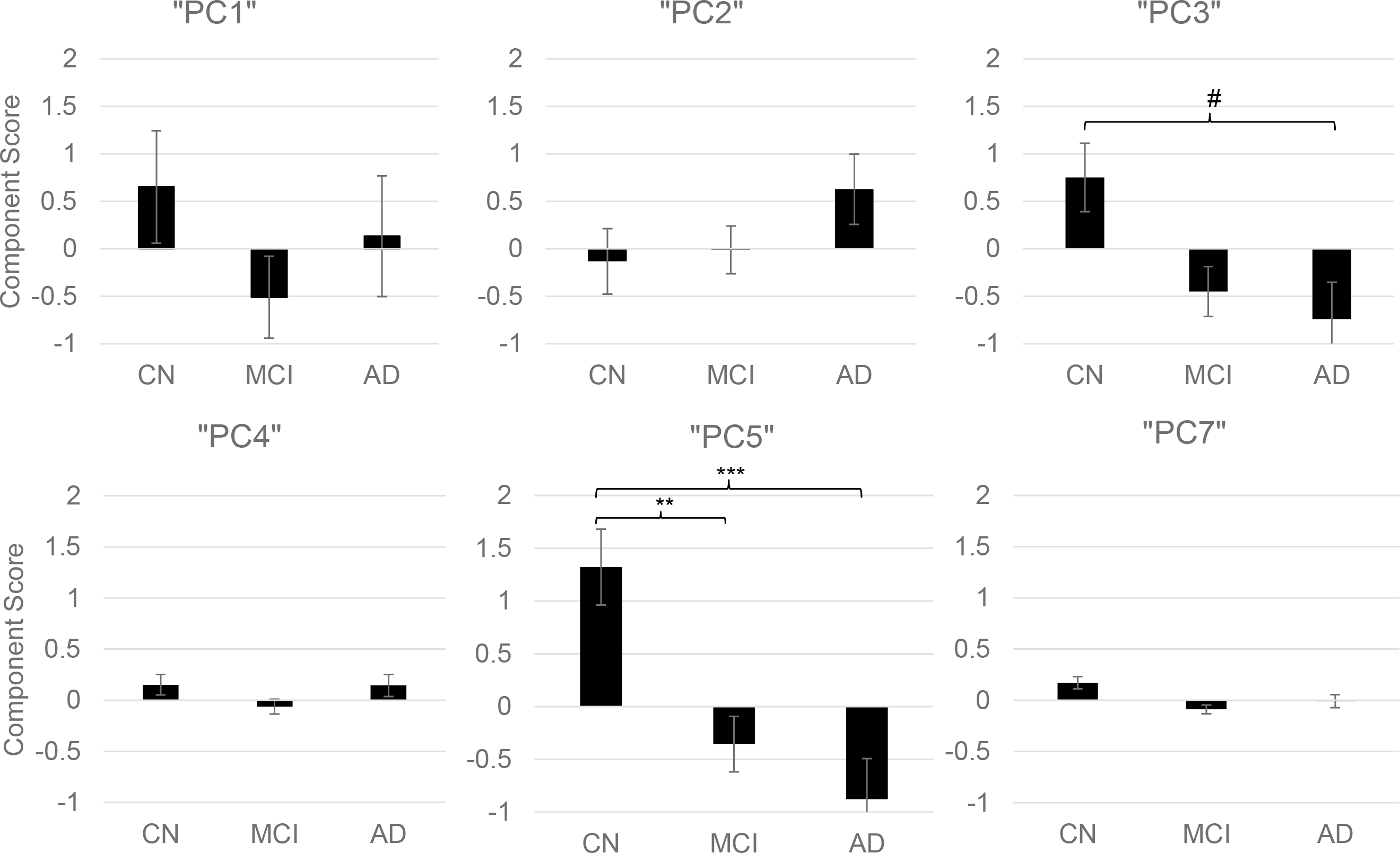
Group differences of principal components of triglycerides with diagnosis groups (CN, MCI, and AD) (# = 0.0533, * < 0.05, ** <0.01, ***<0.001) Multivariate GLM of diagnostic (CN, MCI, and AD) group differences in principal components of triglycerides. Covariates included: age, sex, body mass index (BMI), total triglycerides, and APOE ε4 status. mean +/− standard error. *Abbreviations*: PC: principal component; CN: cognitively normal older adult controls; MCI: mild cognitive impairment; AD: Alzheimer’s disease.

### Association of TG principal components with AD biomarkers

The two principal components that showed significant (“PC5”) or suggestive (“PC3”) diagnosis group differences were further investigated to assess their associations with continuous A/T/N/V biomarkers for AD. Figure 2 shows associations of “PC3” and “PC5” with AD biomarkers. Linear regression analysis indicated a significant association between “PC5” and hippocampal volume (“N”) (*p*-value =0.0243, standardized β=0.135, adjusted R^2^ = 0.0683; Figure 2) and “PC3” and entorhinal thickness (“N”) (*p-*value=0.00363, standardized β=0.132, adjusted R^2^ = 0.188; Figure 2). Lower “PC5” scores were associated with greater brain atrophy. As “PC5” is associated with hippocampal volume, we investigated the association of “PC5” with cognitive performance using ADAS-Cog13 scores and ADNI-MEM. The analysis revealed associations between ADAS-Cog13 (*p*-value =0.012, standardized β = −0.099, adjusted R^2^ = 0.0579) and ADNI-MEM (*p*-value =0.00443, standardized β = 0.115, adjusted R^2^ = 0.0613) with “PC5.” We then performed a detailed whole-brain surface-based analysis of cortical thickness to investigate the effects of “PC3” and “PC5” on cortical atrophy in a spatially unbiased manner. Lower component scores of “PC3” were significantly associated with reduced cortical thickness in bilateral frontal and parietal lobes and right temporal lobe including the entorhinal cortex (*p*-value < 0.05; Figure 3(A)). Also, lower component scores of “PC5” were significantly associated with reduced cortical thickness in right temporal lobe including the entorhinal cortex (*p*-value < 0.05; Figure 3(B)).

**Figure 2:**
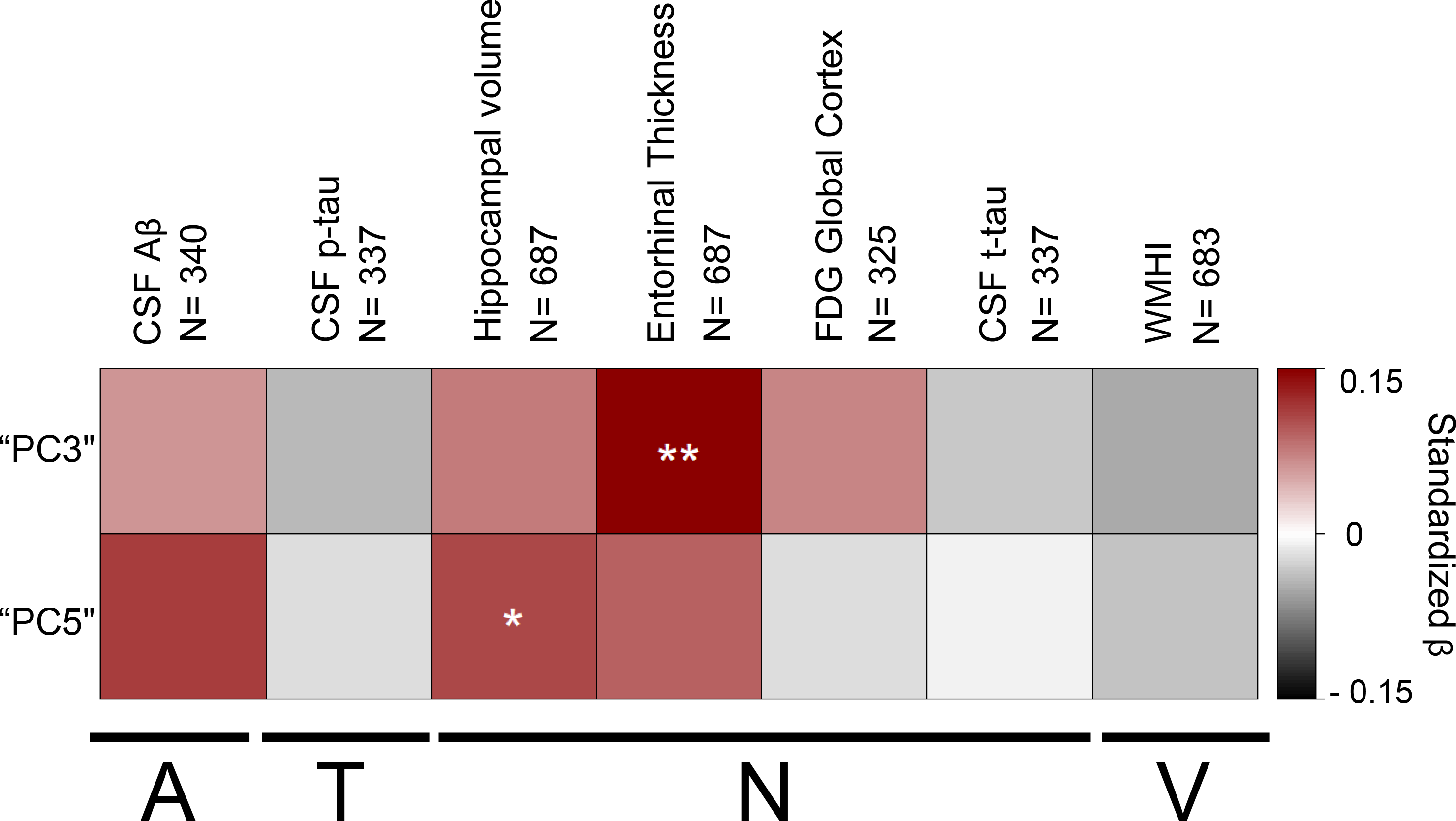
Association of triglyceride “PC3” and “PC5” with A/T/N/V biomarkers for AD. (* < 0.05, ** <0.01) Linear regression performed for “PC3” and “PC5” with AD endophenotypes. Covariates included: age, sex, body mass index (BMI), total triglycerides, and APOE ε4 status for all A/T/N/V phenotypes. For MRI biomarkers, we also included years of education and intracranial volume (ICV) as additional covariates. The y-axis colors represent standardized β values from the linear regression analysis, with shades of red indicating a positive standardized β value and gray scale a negative standardized β value. *Abbreviations*: PC: principal component; CSF: cerebrospinal fluid; CSF Aβ: CSF amyloid β 1-42 peptide (Aβ_1-42_); CSF p.tau: CSF tau phosphorylated at threonine 181 (CSF p-tau_181P_); CSF t.tau: CSF total tau (CSF t-tau); WMHI: white matter hyperintensity total volume, FDG Global cortex: cortical glucose SUVR measured from [^18^F]FDG PET scans; Hippocampal.volume: hippocampal volume; Entorhinal.thickness: entorhinal cortical thickness, “A”= Aβ_1-42_ levels as a biomarker of amyloid-β, “T”= CSF p-tau levels as a biomarker of tau, “N”= structural atrophy on MRI, FDG PET metabolism, and CSF t-tau levels as biomarkers of neurodegeneration, and “V”= white matter hyperintensity volume as a biomarker for microvascular disease burden.

**Figure 3:**
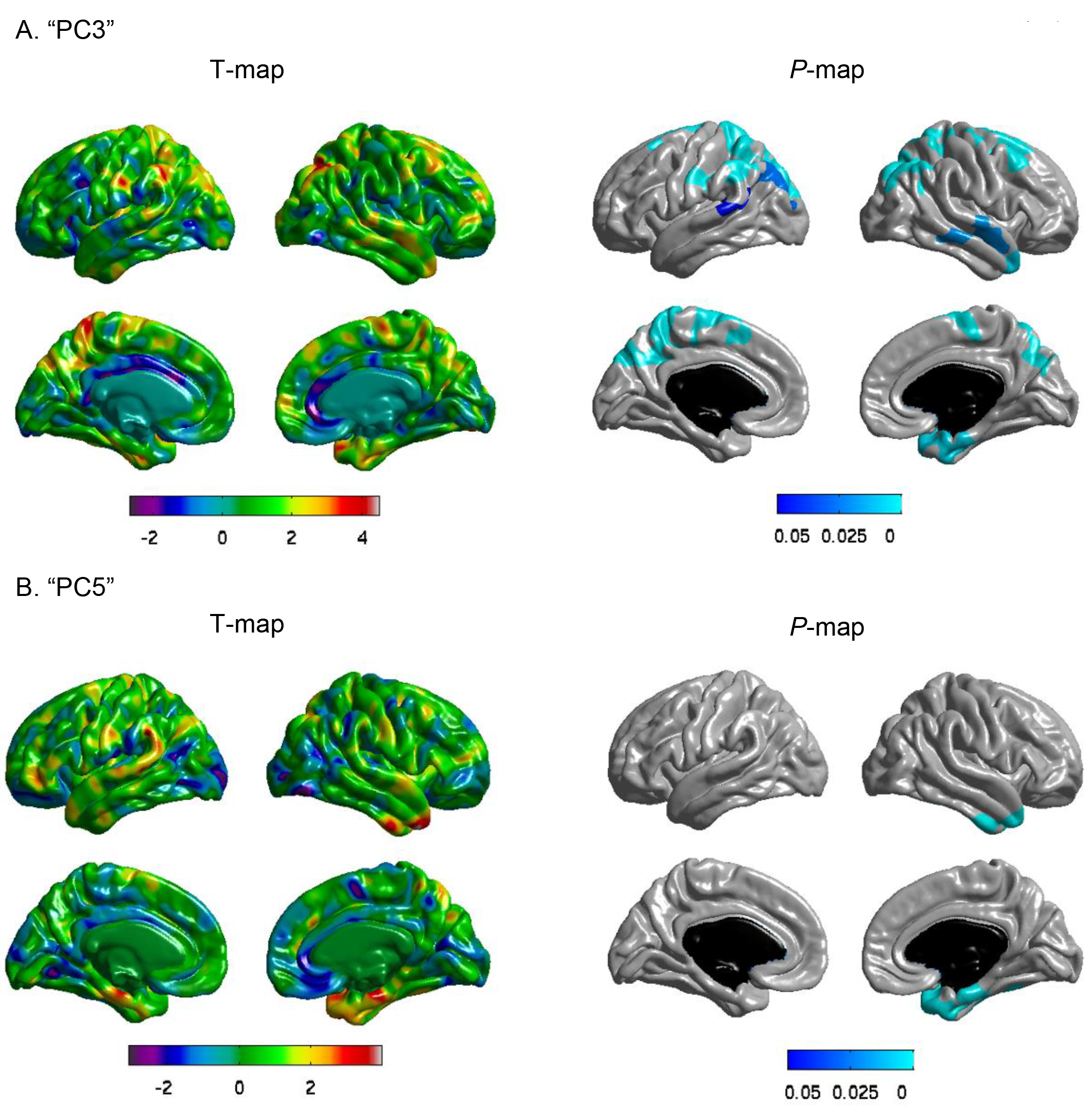
Whole-brain surface-based analysis of cortical thickness for “PC3” and “PC5”. A whole-brain multivariate analysis of cortical thickness across the brain surface was performed to identify the association of two principal components ((a) “PC3” and (b) “PC5”) with brain structure shown as a T-value map and a P-value map. Statistical maps were thresholded using a random field theory for a multiple testing adjustment to a significance level of 0.05. Positive t values (red, yellow) indicate thicker cortical thickness. The *p*-value for clusters indicates significant *p* values with the lightest blue color. Covariates included: age, sex, body mass index (BMI), total triglycerides, APOE ε4 status, years of education, and intracranial volume (ICV).

### Effect of *APOE* ε4 on TGs

In order to investigate the effect of *APOE* ε4 on TGs, we first investigated the presence of an interaction between *APOE* ε4 status and diagnosis and A/T/N/V biomarkers for AD with principal components. We did not find evidence of significant interactions for any principal components with APOE ε4 carrier status. We then performed an association analysis of principal components with diagnosis and A/T/N/V biomarkers for AD after stratifying on *APOE* ε4 carrier status. In both the *APOE* ε4 carrier group and *APOE* ε4 non-carrier group, we did not find any significant associations of principal components with diagnosis (**Figure e-2**). However, in the *APOE* ε4 carrier group, “PC5” was significantly associated with CSF Aβ_1-42_ levels (*p*-value= 0.0359, standardized β =0.228, adjusted R2=0.101; Figure 4) and marginally associated with entorhinal cortical thickness (*p*-value=0.0537, standardized β =0.156, adjusted R2= 0.073; Figure 4). “PC3” was also significantly associated with entorhinal cortical thickness (*p*-value= 9.66E-04, standardized β =0.192, adjusted R2= 0.267) (Figure 4). In the *APOE* ε4 non-carrier group, we did not identify any significant associations of principal components with A/T/N/V biomarkers for AD.

**Figure 4:**
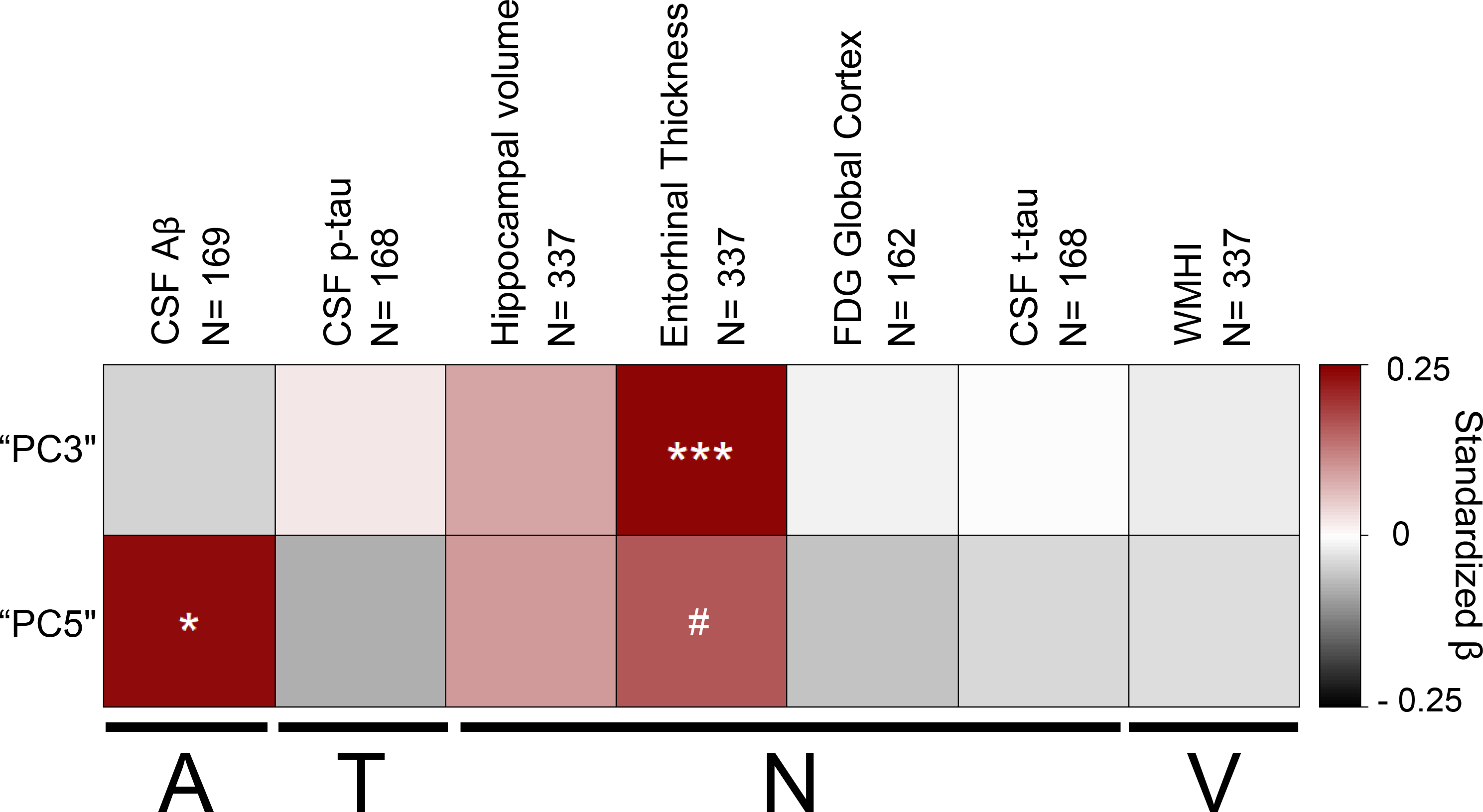
Association of “PC3” and “PC5” with A/T/N/V biomarkers for AD in the *APOE* ε4 carrier group. (# = 0.0537, * < 0.05, ** <0.01, ***<0.001) Linear regression performed for “PC3” and “PC5” with AD endophenotypes in *APOE* ε4 carrier stratified groups. Covariates included: age, sex, body mass index (BMI), total triglycerides, and APOE ε4 carrier status for all A/T/N/V phenotypes. For MRI biomarkers, we also included years of education and intracranial volume (ICV) as additional covariates. The y-axis colors represent standardized β values from the linear regression analysis, with shades of red indicating a positive standardized β value and gray scale a negative standardized β value. *Abbreviations*: PC: principal component; CSF: cerebrospinal fluid; CSF Aβ: CSF amyloid β 1-42 peptide (Aβ_1-42_); CSF p.tau: CSF tau phosphorylated at threonine 181 (CSF p-tau_181P_); CSF t.tau:CSF total tau (CSF t-tau); WMHI: white matter hyperintensity volume, FDG Global cortex:: cortical glucose SUVR measured from [^18^F]FDG PET scans; Hippocampal.volume: hippocampal volume; Entorhinal.thickness: entorhinal cortical thickness, “A”= Aβ_1-42_ levels as a biomarker of amyloid-β, “T”= CSF p-tau levels as a biomarker of tau, “N”= structural atrophy on MRI, FDG PET metabolism, and CSF t-tau levels as biomarkers of neurodegeneration, and “V”= white matter hyperintensity volume as a biomarker for microvascular disease burden.

### Effect of medication use on TGs

Using TG values adjusted for the effect of medication use at baseline as a potential confounder, we repeated all analyses. All key findings remained significant after adjustment for medication use (see **Figures e-3 and e-4**).

## Discussion

In this study, we found that long-chain, polyunsaturated FA-containing triglycerides (PUTGs) showed significant differences between diagnostic groups (CN, MCI, AD; **Table e-3** and **e-4**). Lower PUTG component scores in MCI suggest that the changes occur during prodromal stages of disease, though longitudinal studies will be required to confirm this cross-sectional analysis. PUTG component scores were significantly associated with early-AD biomarkers, including hippocampal volume and entorhinal cortical thickness measured from MRI scans. In addition, we observed a significant *APOE* ε4 effect on PUTG components. In *APOE* ε4 carriers, we found significant, positive associations between component scores of PUTGs and entorhinal cortical thickness and CSF Aβ_1-42_ levels, but no significant associations in *APOE ε4* non-carriers. The pattern of observed principal component scores suggest that reduction of PUTGs are associated with early stage changes in AD.

The association of PUTGs with atrophy in the entorhinal cortex and hippocampus is noteworthy as these regions are affected in early stages of AD pathophysiology. Aβ accumulation is an important early change in AD and is associated with *APOE* ε4 carrier status. Triglycerides (TGs) have been shown to be particularly associated with Aβ. For example, mouse studies showed that serum TG levels were elevated prior to amyloid deposition,^25^ while human studies showed increased serum TGs were associated with increased amyloidosis in cognitively normal individuals.^3^ Longitudinal studies showed that increased midlife TGs predicted amyloidosis and tau pathology 20 years later,^2^ providing additional evidence for the early influence of TGs in AD. Thus, it is particularly interesting that we found a selective association of PUTGs with CSF Aβ only in *APOE* ε4 carriers. The specific mechanistic role underlying the association of decreased PUTGs and early processes involved in AD pathogenesis remains to be determined, but our results suggest a relationship between a decrease in component scores of PUTGs and early-stage biomarkers in AD. Results remained significant after adjusting for medication use. Thus, exposure to medication by AD and some MCI patients does not appear to account for these results, although some roles cannot be entirely ruled out.

The relationship between TGs and amyloid-β may be explained through role of TGs in the lipoprotein peripheral transport of Aβ. Apolipoprotein E (ApoE) is a plasma protein that is involved in lipid transport and metabolism in lipoproteins, including TG-rich lipoproteins.^26^ Studies suggest that TG-rich lipoproteins may participate in peripheral Aβ transport and delivery, as evidenced by a study that showed Aβ accumulated in TG-rich lipoproteins.^27^ As discussed above, ApoE exists in three major isoforms, ApoE ε4, ApoE ε3, and ApoE ε2. ApoE ε4 has been shown unable to complex with Aβ and, peripherally, to associate with triglyceride-rich, very low-density lipoproteins (VLDLs) in contrast to high-density lipoproteins (HDL).^28,29^ These studies suggest an important relationship between Aβ and peripheral TG lipoprotein carriers that may provide further insight into aberrant serum PUTG levels in AD.

Omega-3 polyunsaturated fatty acids such as docosahexaenoic acid (DHA) and eicosapentaenoic acid (EPA), have anti-inflammatory and neuroprotective roles, as evidenced by their associations with cognitive improvement in the elderly.^30^ In blood, polyunsaturated fatty acids travel in different states: esterified, bound to complex lipids (such as TGs) or lipoproteins, or free-floating. Triglycerides are created by incorporating three fatty acids onto a carbon backbone, which can then serve to transport and store fatty acids. A previous study, using the same TG dataset, identified EPA and DHA as the primary polyunsaturated fatty acids contributing to PUTG fatty acids of “PC5”.^31^ Recently published studies have shown that serum DHA and EPA levels were decreased in AD,^32^ and that polyunsaturated fatty acid intake acted to reduced risk for AD.^33^ We found that lower component scores of PUTGs were associated with poorer cognitive performance. Due to the importance of these polyunsaturated fatty acids in cognition and neuroinflammation, we believe our findings need further study to determine the overlapping mechanism that may underlie PUTG and polyunsaturated fatty acid aberrations. PUTGs and, by extension, polyunsaturated fatty acids represent links in the gut-liver-brain axis. Triglycerides are synthesized in the liver or absorbed from the gut and have recently been found to cross the blood-brain barrier and accumulate in the brain.^34^ PUFAs are rapidly and preferentially incorporated into TGs following postprandial ingestion, suggesting an important role for TGs in PUFA supplementation.^35^ The gut-liver-brain communicate through neuronal, neuroimmune, and neuroendocrine pathways. These pathways overlap polyunsaturated fatty acids and AD and may serve an interesting direction to study PUTGs.

A Lipid Hypothesis has been proposed, suggesting that lipid oxidation is the initiating factor for late-onset AD.^36^ Lipid oxidation is a key early event in AD that precedes amyloid and neurofibrillary tangle deposition. When lipids are exposed to free radicals, they progress through autoperoxidation, where reactive oxidative species are released. Previous studies have shown that Aβ aggregation occurred more readily in membranes composed of oxidized lipids,^37^ and that polyunsaturated lipids were most vulnerable to oxidative stress.^38^ Early-stage AD is characterized by an accumulation of Aβ, potentially resulting from impaired clearance of pathogenic species by microglia. Early microglial activity is neuroprotective, but progressive cytokine production from microglia, due to aging and disease progression, can reduce Aβ clearance and promote its accumulation. Peripheral inflammatory cytokines have been shown to interact with or pass through endothelial cells on the blood-brain barrier to activate microglia. It is well established that polyunsaturated fatty acid intake is associated with an anti-inflammatory effect. Polyunsaturated fatty acids have been directly linked to the regulation of cytokines by decreasing the expression of proinflammatory pathways. Moreover, polyunsaturated fatty acid intake has been shown to decrease microglial inflammatory activation of Aβ, supporting a neuroprotective role.^39^ Decreases in PUTGs could presumably result in decreased availability for polyunsaturated fatty acids and loss of neuroprotective metabolites, thereby suggesting the need for further study into this relationship.

Neural connections between the brain and the gut exist through the vagus nerve and enteric system. The vagus nerve originates in the brain stem and complexes onto the enteric plexus in the gut. These neural connections are important for gastric motility and have been implicated in relaying inflammatory, microbial and nutrient information from the gut to the brain.^40^ Interestingly, dietary fat has shown to activate the vagus nerve and neuroimmunologic pathways.^41^ Recent work in Parkinson’s disease found that severance of the vagal trunk was protective against PD, suggesting that central nervous system (CNS) invasion may occur from the gut, through the vagus nerve.^42^ This study provides an interesting avenue of study for AD due to many links between PD and AD.

The gastrointestinal tract is home to more than 10 times the number of bacteria than cells in the human body, functioning in a mutualistic relationship with their human host.^43^ Gut microbiota are known to function in immune responses, nutrient absorption, and regulate gut motility. ^44,45^ When gut microbiota is no longer homeostatic (gut dysbiosis), the CNS receives signals to activate inflammatory processes. Alterations in gut microbiota have been linked to neurological conditions, including AD,^46^ where gut dysbiosis was associated with memory dysfunction and decreased hippocampal neurogenesis.^47^ Polyunsaturated fatty acid intake has been associated with changes in gut microbiota and intestinal excretion of mucosal defense factors that exhibit anti-inflammatory effects.^48^ The data suggests an interesting relationship between polyunsaturated fatty acids, gut microbiota and AD and an interesting direction to study PUTGs. We did not see a significant association between saturated TGs and AD, although we did find a significant difference in PUTG component scores between diagnostic groups. A recent meta-analysis revealed that, over time, saturated fat intake was associated with an increase in AD risk.^49^ It is possible that blood TG levels do not accurately represent long-term saturated TG intake and, rather, BMI may better represent the effects of highly saturated diets over time.^50^ Further studies are required to investigate these findings.

There are some limitations to our study. The ADNI observational cohort was designed to be typical of participants who enroll in clinical trials, but is not necessarily representative of the broader community as would be found in epidemiologically derived samples. As ADNI is a largely white sample with high mean education, the present results should not be generalized to community-based populations without further investigation. It will be important to repeat these analyses in more socioeconomically, educationally and racially diverse samples. In addition, this is a cross-sectional study investigating early changes in AD. Longitudinal and model system studies are required to determine the role and mechanisms of specific classes of TGs in AD initiation and progression. Mechanistic investigations into the cause of decreased PUTG component scores using mouse models of AD may provide insight relevant to early detection and treatment for AD. Replication in independent samples and longitudinal follow-up of the present cohort will also be important.

In summary, our study investigated the relationship between TG species, AD risk and biomarkers for AD. To our knowledge, this is the first study to show decreased levels of highly unsaturated, long-chain triglycerides in MCI and AD compared to cognitively normal older adults. We also observed an association between decreased PUTG component scores with increased brain atrophy, decreased CSF amyloid-β concentration, and the effect of APOE ε4 carrier status on PUTG components in AD. Our findings identify a specific subcategory of TGs, namely the PUTGs, which appear mechanistically relevant and provide the foundation for future work in therapeutic development. We provide evidence that PUTGs are associated with an early prodromal stage of cognitive impairment (i.e., MCI) and early stage biomarkers for AD. These results suggest a potential role for PUTGs as a target for an early detection biomarker as well as for therapeutic development. In addition to the need for independent replication, longitudinal designs are needed to elucidate causal directionality.

## Data Availability

Data use restrictions prohibit the distribution of any ADNI clinical or demographic data outside of LONI. Researchers can apply for access to the ADNI data at http://adni.loni.usc.edu/data-samples/access-data/. Data for the ADNI-1 cohort is accessible via http://dx.doi.org/10.7303/syn5592519.

## Disclosures of all authors’ financial relationships

J.Q.T. may accrue revenue in the future on patents submitted by the University of Pennsylvania wherein he is a co-inventor and he received revenue from the sale of Avid to Eli Lily as a co-inventor on imaging-related patents submitted by the University of Pennsylvania. L.M.S. receives research funding from NIH (U01 AG024904; R01 MH 098260; R01 AG 046171; 1RF AG 051550); MJFox Foundation for PD Research and is a consultant for Eli Lilly; Novartis; Roche; he provides QC over-sight for the Roche Elecsys immunoassay as part of responsibilities for the ADNI study. A.J.S. reports investigator-initiated research support from Eli Lilly unrelated to the work reported here. He has received consulting fees and travel expenses from Eli Lilly and Siemens Healthcare and is a consultant to Arkley BioTek. He also receives support from Springer-Nature Publishing as Editor in Chief of *Brain Imaging and Behavior*. M.W.W. reports stock/stock options from Elan, Synarc, travel expenses from Novartis, Tohoku University, Fundacio Ace, Travel eDreams, MCI Group, NSAS, Danone Trading, ANT Congress, NeuroVigil, CHRU-Hopital Roger Salengro, Siemens, AstraZeneca, Geneva University Hospitals, Lilly, University of California, San Diego–ADNI, Paris University, Institut Catala de Neurociencies Aplicades, University of New Mexico School of Medicine, Ipsen, Clinical Trials on Alzheimer’s Disease, Pfizer, AD PD meeting. PMD has received research grants from Lilly, Neuronetrix, Avanir, Alzheimer’s Drug Discovery Foundation, ADNI, DOD, ONR and NIH. PMD has received speaking or advisory fees from Anthrotronix, Neuroptix, Genomind, MindLink and CEOs Against Alzheimer’s. PMD owns shares in Muses Labs, Anthrotronix, Evidation Health, Turtle Shell Technologies and Advera Health Analytics whose products are not discussed here. PMD serves on the board of Baycrest, Apollo Hospitals, Goldie Hawn Foundation and Live Love Laugh Foundation. PMD is a co-inventor (through Duke) on patents relating to dementia biomarkers and therapies. R.K.D. is inventor on key patents in the field of metabolomics including applications for Alzheimer disease. All other authors report no disclosures.

## Appendix: authors

| Name | Location | Role | Contribution |
| --- | --- | --- | --- |
| Megan Bernath, BS | Indiana University,<br>Indianapolis | Author | Study design, statistical analysis, drafted and revised manuscript |
| Sudeepa Bhattacharyya, PhD | University of Arkansas for<br>Medical Sciences | Author | Initial analysis of data and edited manuscript |
| Kwangsik Nho, PhD | Indiana University,<br>Indianapolis | Author | Design and conceptualization of study, edited manuscript |
| Shannon L Risacher, PhD | Indiana University,<br>Indianapolis | Author | MRI and PET image processing |
| Dinesh Kumar Barupal, PhD | Genome Center,<br>University of California,<br>Davis, California | Author | Contributed lipidomics data and edited manuscript. |
| Oliver Fiehn, PhD | Genome Center,<br>University of California,<br>Davis, California | Author | Contributed lipidomics data and edited manuscript. |
| Rebecca Baillie, PhD | Rosa & Co LLC, San<br>Carlos, CA | Author | Interpretation of data and edited manuscript |
| Matthias Arnold, PhD | Helmholtz Zentrum<br>munchen, Neuherberg,<br>Germany | Author | Contributed to quality control<br>of data set |
| Tanner Jacobson, BS | Indiana University,<br>Indianapolis | Author | Contribution to data analysis<br>and interpretation |
| John Q. Trojanowski, MD,<br>PhD | University of<br>Pennsylvania,<br>Philadelphia | Author | Biomarker core coleader,<br>provided samples and data<br>interpretation |
| Leslie M. Shaw, PhD | University of<br>Pennsylvania,<br>Philadelphia | Author | Biomarker core coleader,<br>provided samples and data<br>interpretation |
| Michael W. Weiner, MD | University of California,<br>San Francisco and San<br>Francisco VAMC | Author | PI of ADNI, study design, data<br>interpretation |
| P. Murali Doraiswamy | Duke University Medical<br>Center | Author | Study design, data<br>interpretation |
| Rima Kaddurah Daouk, PhD | Duke University Medical<br>Center | Author | PI of ADMC the consortium<br>that conceptualized design,<br>secured funding and executed<br>study |
| Andrew J. Saykin, PsyD | Indiana University,<br>Indianapolis | Author | Design and conceptualization of<br>study, statistical analysis and interpretation, edited<br>manuscript |

## References

1. Østergaard SD, Mukherjee S, Sharp SJ, et al. Associations between Potentially Modifiable Risk Factors and Alzheimer Disease: A Mendelian Randomization Study. PLoS Med. 2015;12:e1001841; discussion e1001841.

2. Nägga K, Gustavsson A-M, Stomrud E, et al. Increased midlife triglycerides predict brain β-amyloid and tau pathology 20 years later. Neurology. 2018;90:e73–e81.

3. Choi HJ, Byun MS, Yi D, et al. Association Between Serum Triglycerides and Cerebral Amyloidosis in Cognitively Normal Elderly. The American Journal of Geriatric Psychiatry. 2016;24:604–612.

4. Lepara O, Valjevac A, Alajbegović A, Zaćiragić A, Nakas-Ićindić E. Decreased serum lipids in patients with probable Alzheimer’s disease. Bosn J Basic Med Sci. 2009;9:215–220.

5. Hall K, Murrell J, Ogunniyi A, et al. Cholesterol, APOE genotype, and Alzheimer disease: an epidemiologic study of Nigerian Yoruba. Neurology. 2006;66:223–227.

6. Saykin AJ, Shen L, Foroud TM, et al. Alzheimer’s Disease Neuroimaging Initiative biomarkers as quantitative phenotypes: Genetics core aims, progress, and plans. Alzheimers Dement. 2010;6:265–273.

7. Saykin AJ, Shen L, Yao X, et al. Genetic Studies of Quantitative MCI and AD Phenotypes in ADNI: Progress, Opportunities, and Plans. Alzheimers Dement. 2015;11:792–814.

8. Cajka T, Fiehn O. LC-MS-Based Lipidomics and Automated Identification of Lipids Using the LipidBlast In-Silico MS/MS Library. Methods Mol Biol. 2017;1609:149–170.

9. Kind T, Liu K-H, Lee DY, DeFelice B, Meissen JK, Fiehn O. LipidBlast in silico tandem mass spectrometry database for lipid identification. Nat Methods. 2013;10:755–758.

10. Toledo JB, Arnold M, Kastenmüuller G, et al. Metabolic network failures in Alzheimer’s disease: A biochemical road map. Alzheimers Dement. 2017;13:965–984.

11. Jack CR, Bernstein MA, Fox NC, et al. The Alzheimer’s Disease Neuroimaging Initiative (ADNI): MRI methods. J Magn Reson Imaging. 2008;27:685–691.

12. Jack CR, Bernstein MA, Borowski BJ, et al. Update on the magnetic resonance imaging core of the Alzheimer’s disease neuroimaging initiative. Alzheimers Dement. 2010;6:212–220.

13. Risacher SL, Kim S, Shen L, et al. The role of apolipoprotein E (APOE) genotype in early mild cognitive impairment (E-MCI). Front Aging Neurosci. 2013;5:11.

14. Fischl B, Sereno MI, Dale AM. Cortical Surface-Based Analysis: II: Inflation, Flattening, and a Surface-Based Coordinate System. NeuroImage. 1999;9:195–207.

15. Dale AM, Fischl B, Sereno MI. Cortical Surface-Based Analysis: I. Segmentation and Surface Reconstruction. NeuroImage. 1999;9:179–194.

16. Jagust WJ, Bandy D, Chen K, et al. The Alzheimer’s Disease Neuroimaging Initiative positron emission tomography core. Alzheimer’s & Dementia. 2010;6:221–229.

17. Carmichael O, Schwarz C, Drucker D, et al. Longitudinal changes in white matter disease and cognition in the first year of the Alzheimer disease neuroimaging initiative. Arch Neurol. 2010;67:1370–1378.

18. Schwarz C, Fletcher E, DeCarli C, Carmichael O. Fully-automated white matter hyperintensity detection with anatomical prior knowledge and without FLAIR. Inf Process Med Imaging. 2009;21:239–251.

19. Hansson O, Seibyl J, Stomrud E, et al. CSF biomarkers of Alzheimer’s disease concord with amyloid-β PET and predict clinical progression: A study of fully automated immunoassays in BioFINDER and ADNI cohorts. Alzheimers Dement. Epub 2018 Mar 1.

20. Mohs RC, Knopman D, Petersen RC, et al. Development of cognitive instruments for use in clinical trials of antidementia drugs: additions to the Alzheimer’s Disease Assessment Scale that broaden its scope. The Alzheimer’s Disease Cooperative Study. Alzheimer Dis Assoc Disord. 1997;11 Suppl 2:S13–21.

21. Crane PK, Carle A, Gibbons LE, et al. Development and assessment of a composite score for memory in the Alzheimer’s Disease Neuroimaging Initiative (ADNI). Brain Imaging Behav. 2012;6:502–516.

22. Gibbons LE, Carle AC, Mackin RS, et al. A composite score for executive functioning, validated in Alzheimer’s Disease Neuroimaging Initiative (ADNI) participants with baseline mild cognitive impairment. Brain Imaging Behav. 2012;6:517–527.

23. Jack CR, Bennett DA, Blennow K, et al. NIA-AA Research Framework: Toward a biological definition of Alzheimer’s disease. Alzheimers Dement. 2018;14:535–562.

24. Nho K, Risacher SL, Crane PK, et al. Voxel and Surface-Based Topography of Memory and Executive Deficits in Mild Cognitive Impairment and Alzheimer’s Disease. Brain Imaging Behav. 2012;6:551–567.

25. Burgess BL, McIsaac SA, Naus KE, et al. Elevated plasma triglyceride levels precede amyloid deposition in Alzheimer’s disease mouse models with abundant A beta in plasma. Neurobiol Dis. 2006;24:114–127.

26. Mensenkamp AR, Jong MC, van Goor H, et al. Apolipoprotein E participates in the regulation of very low density lipoprotein-triglyceride secretion by the liver. J Biol Chem. 1999;274:35711–35718.

27. James AP, Pal S, Gennat HC, Vine DF, Mamo JCL. The incorporation and metabolism of amyloid-beta into chylomicron-like lipid emulsions. J Alzheimers Dis. 2003;5:179–188.

28. Weisgraber KH. Apolipoprotein E distribution among human plasma lipoproteins: role of the cysteine-arginine interchange at residue 112. J Lipid Res. 1990;31:1503–1511.

29. Zhou Z, Smith JD, Greengard P, Gandy S. Alzheimer amyloid-beta peptide forms denaturant-resistant complex with type epsilon 3 but not type epsilon 4 isoform of native apolipoprotein E. Mol Med. 1996;2:175–180.

30. Kesse-Guyot E, Péneau S, Ferry M, et al. Thirteen-year prospective study between fish consumption, long-chain n-3 fatty acids intakes and cognitive function. J Nutr Health Aging. 2011;15:115–120.

31. Barupal DK, Fan S, Wancewicz B, et al. Generation and quality control of lipidomics data for the alzheimer’s disease neuroimaging initiative cohort. Scientific Data. 2018;5:180263.

32. Shang J, Yamashita T, Fukui Y, et al. Different Associations of Plasma Biomarkers in Alzheimer’s Disease, Mild Cognitive Impairment, Vascular Dementia, and Ischemic Stroke. J Clin Neurol. 2018;14:29–34.

33. Zhang Y, Chen J, Qiu J, Li Y, Wang J, Jiao J. Intakes of fish and polyunsaturated fatty acids and mild-to-severe cognitive impairment risks: a dose-response meta-analysis of 21 cohort studies. Am J Clin Nutr. 2016;103:330–340.

34. Banks WA, Farr SA, Salameh TS, et al. Triglycerides cross the blood–brain barrier and induce central leptin and insulin receptor resistance. Int J Obes (Lond). 2018;42:391–397.

35. Plourde M, Chouinard-Watkins R, Vandal M, et al. Plasma incorporation, apparent retroconversion and β-oxidation of 13C-docosahexaenoic acid in the elderly. Nutr Metab (Lond). 2011;8:5.

36. Cooper JL. Dietary Lipids in the Aetiology of Alzheimer’s Disease. Drugs Aging. 2003;20:399–418.

37. Koppaka V, Axelsen PH. Accelerated accumulation of amyloid beta proteins on oxidatively damaged lipid membranes. Biochemistry. 2000;39:10011–10016.

38. Buettner GR. The Pecking Order of Free Radicals and Antioxidants: Lipid Peroxidation, α-Tocopherol, and Ascorbate. Archives of Biochemistry and Biophysics. 1993;300:535–543.

39. Hopperton KE, Trépanier M-O, Giuliano V, Bazinet RP. Brain omega-3 polyunsaturated fatty acids modulate microglia cell number and morphology in response to intracerebroventricular amyloid-β 1-40 in mice. J Neuroinflammation [online serial]. 2016;13. Accessed at: https://www.ncbi.nlm.nih.gov/pmc/articles/PMC5041295/. Accessed August 16, 2018.

40. Miao FJ-P, Green PG, Levine JD. Mechanosensitive duodenal afferents contribute to vagal modulation of inflammation in the rat. J Physiol (Lond). 2004;554:227–235.

41. Luyer MD, Greve JWM, Hadfoune M, Jacobs JA, Dejong CH, Buurman WA. Nutritional stimulation of cholecystokinin receptors inhibits inflammation via the vagus nerve. J Exp Med. 2005;202:1023–1029.

42. Svensson E, Horváth-Puhó E, Thomsen RW, et al. Vagotomy and subsequent risk of Parkinson’s disease. Ann Neurol. 2015;78:522–529.

43. Bäckhed F, Ley RE, Sonnenburg JL, Peterson DA, Gordon JI. Host-bacterial mutualism in the human intestine. Science. 2005;307:1915–1920.

44. Olszak T, An D, Zeissig S, et al. Microbial exposure during early life has persistent effects on natural killer T cell function. Science. 2012;336:489–493.

45. Bäckhed F, Ding H, Wang T, et al. The gut microbiota as an environmental factor that regulates fat storage. Proc Natl Acad Sci USA. 2004;101:15718–15723.

46. Zhuang Z-Q, Shen L-L, Li W-W, et al. Gut Microbiota is Altered in Patients with Alzheimer’s Disease. J Alzheimers Dis. 2018;63:1337–1346.

47. Desbonnet L, Clarke G, Traplin A, et al. Gut microbiota depletion from early adolescence in mice: Implications for brain and behaviour. Brain Behav Immun. 2015;48:165–173.

48. Kaliannan K, Wang B, Li X-Y, Kim K-J, Kang JX. A host-microbiome interaction mediates the opposing effects of omega-6 and omega-3 fatty acids on metabolic endotoxemia. Sci Rep. 2015;5:11276.

49. Ruan Y, Tang J, Guo X, Li K, Li D. Dietary Fat Intake and Risk of Alzheimer’s Disease and Dementia: A Meta-Analysis of Cohort Studies. Curr Alzheimer Res. Epub 2018 Apr 27.

50. Hariri N, Gougeon R, Thibault L. A highly saturated fat-rich diet is more obesogenic than diets with lower saturated fat content. Nutr Res. 2010;30:632–643.

