## Supplement data for "Serum triglycerides in Alzheimer’s disease: Relation to neuroimaging and CSF biomarkers"

**Figure e-1: Group differences for TGs of “PC5” between diagnosis groups (CN, MCI, and AD)**

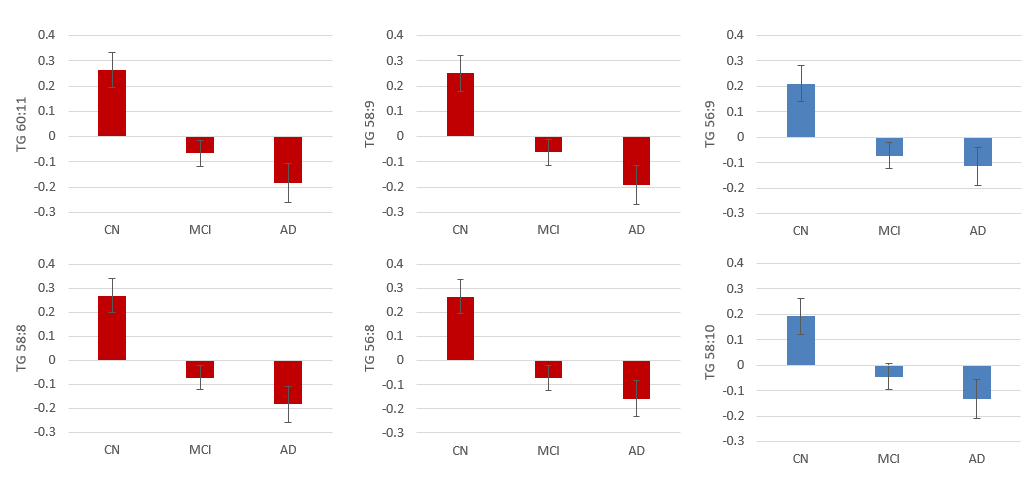

Normalized Peak Height

Shown are 6 PUTGs that contribute to “PC5.” ANCOVA performed for TGs with diagnosis; mean +/- standard error. We see the similar pattern in each PUTG to that was observed with “PC5.” Red graphs represent PUTGs that have a significant association with diagnosis. Bonferroni corrected, p-value cutoff: 5.95E-04 (=0.05/84). (TG=triglyceride; CN= cognitively normal; MCI=mild cognitive impairment; AD=Alzheimer’s disease).

**Figure e-2: Group differences for “PC5” between diagnosis groups (CN, MCI, and AD) in *APOE* ε4 carriers and non-carriers**

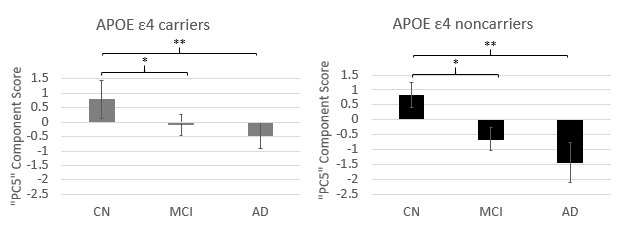

(* < 0.05, **< 5.56E-3)

This figure shows a similar pattern in both *APOE* ε4 carriers and non-carriers. ANCOVA performed for “PC5” with diagnosis in *APOE* ε4 stratified groups; mean +/- standard error. In “PC3,” there is a non-significant association that exists between *APOE* ε4 non-carriers (not shown). In “PC5,” both *APOE* ε4 carriers and non-carriers follow the same significance pattern. *APOE* ε4 carriers are those that have at least one *APOE* ε4 allele, and non-carriers have no *APOE* ε4 alleles. (CN= cognitively normal; MCI=mild cognitive impairment; AD= Alzheimer’s disease).

**Figure e-3:** **Group differences for“PC5” between diagnosis groups (CN, MCI, and AD) using data with and without medication adjustment**

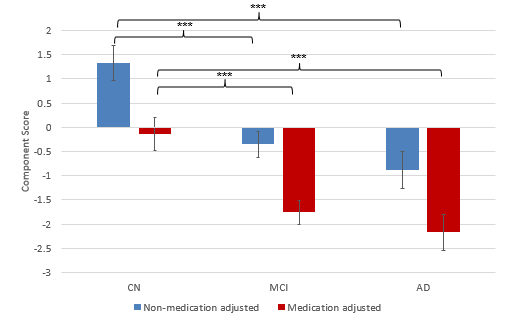

(* > 0.05, **> 5.56E-3 ***>0.001)

Shown are data with medication adjustment (red) and without medication adjustment (blue). ANCOVA performed for “PC5” and diagnosis using data with and without medication adjustment; mean +/- standard error. In “PC3,” we identified a significant association between CN and AD groups (p-value=4.94E-03) that was not present in data without medication adjustment (not shown). In “PC5,” we identified significance associations between CN and MCI or AD in data both with medication adjustment and without medication adjustment. (CN= cognitively normal; MCI=mild cognitive impairment; AD= Alzheimer’s disease).

**Figure e-4: Association of two principal components (PC3 and PC5) with A/T/N/V biomarkers using data with medication adjustment**

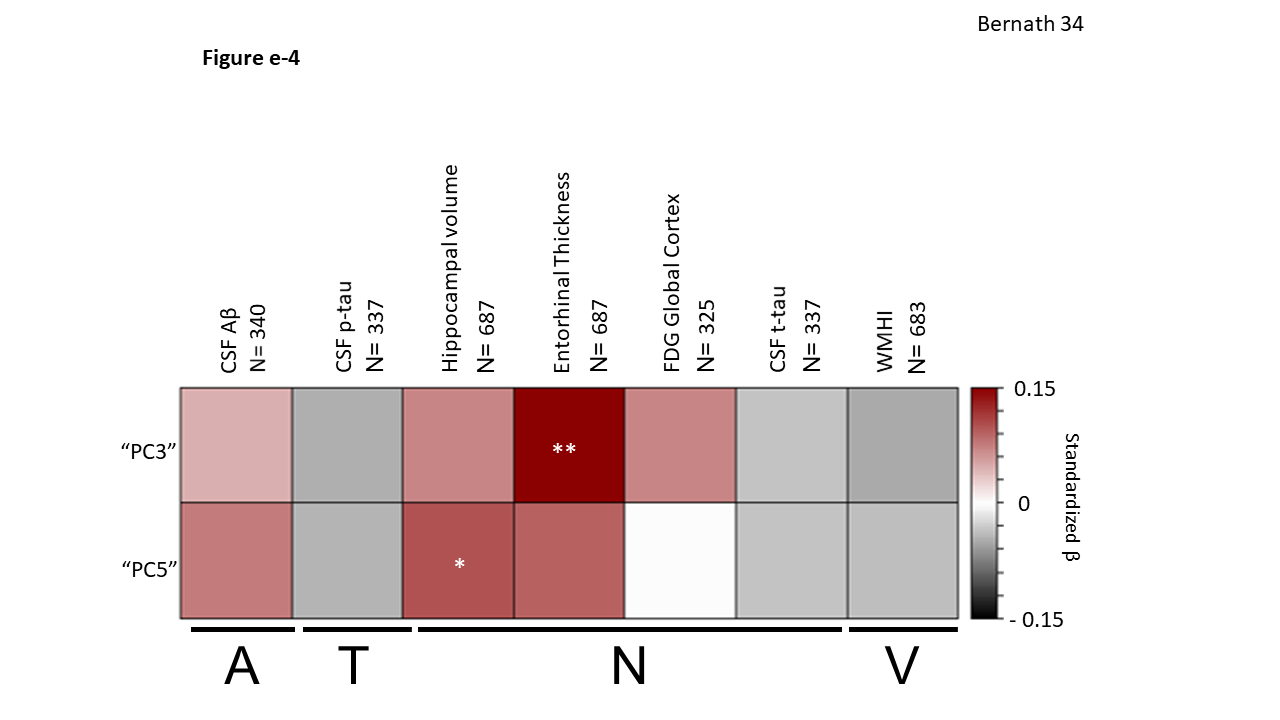

(* > 0.05, ** > 0.01)

Linear regression performed for “PC3” and “PC5” with AD endophenotypes using data with medication adjustment. We see a significant association between entorhinal volume and medication adjusted “PC3” (p-value: 3.136E-03, β: 0.149). In medication adjusted “PC5,” we see a significant association with hippocampal volume (p-value: 0.0407, β: 0.101). *Abbreviations*: PC: principal component; CSF: cerebrospinal fluid; CSF Aβ: CSF amyloid β 1-42 peptide (Aβ_1-42_); CSF p.tau: CSF tau phosphorylated at threonine 181 (CSF p-tau_181P_); CSF t.tau: CSF total tau (CSF t-tau); WMHI: white matter hyperintensity volume, FDG Global cortex: cortical glucose SUVR measured from [^18^F]FDG PET scans; Hippocampal.volume: hippocampal volume; Entorhinal.thickness: entorhinal cortical thickness, “A”= Aβ_1-42_ levels as a biomarker of amyloid-β, “T”= CSF p-tau levels as a biomarker of tau, “N”= structural atrophy on MRI, FDG PET metabolism, and CSF t-tau levels as biomarkers of neurodegeneration, and “V”= white matter hyperintensity volume as a biomarker for microvascular disease burden. The y-axis colors represent standardized β values from the linear regression analysis, with shades of red indicating a positive standardized β value and shades of black a negative standardized β value.

Table e-1: Rotation Sums of Squares Loading Eigenvalues.

| Component | Eigenvalue |
| --- | --- |
| 1 | 21.451 |
| 2 | 14.131 |
| 3 | 13.901 |
| 4 | 12.905 |
| 5 | 6.711 |
| 6 | 2.454 |
| 7 | 1.678 |
| 8 | 1.317 |
| 9 | 1.317 |

Table e-2: Component score contribution of TGs to each principal component. TG = Triglyceride

| Rotated Component Matrix | | | | | | | | | |
| --- | --- | --- | --- | --- | --- | --- | --- | --- | --- |
| Metabolite | Components | | | | | | | | |
|  | 1 | 2 | 3 | 4 | 5 | 6 | 7 | 8 | 9 |
| TG 42:0 | 0.900 | 0.051 | 0.270 | 0.044 | 0.040 | 0.138 | 0.121 | -0.078 | -0.012 |
| TG 40:0 | 0.812 | 0.031 | 0.265 | -0.035 | 0.013 | 0.114 | 0.196 | -0.187 | 0.015 |
| TG 40:1 | 0.056 | 0.078 | 0.048 | 0.012 | -0.053 | 0.244 | 0.118 | -0.022 | 0.795 |
| TG 42:1 | 0.684 | -0.089 | 0.184 | 0.038 | -0.017 | 0.052 | 0.118 | -0.218 | 0.082 |
| TG 42:2 | 0.638 | 0.038 | 0.148 | 0.029 | 0.066 | 0.051 | 0.675 | -0.069 | -0.094 |
| TG 42:3 | 0.067 | -0.034 | -0.067 | 0.086 | 0.021 | 0.088 | 0.862 | 0.025 | 0.070 |
| TG 44:0 | 0.916 | 0.045 | 0.250 | 0.173 | 0.087 | 0.091 | 0.055 | 0.042 | -0.029 |
| TG 44:1 | 0.936 | 0.072 | 0.240 | 0.176 | 0.053 | 0.012 | 0.032 | 0.017 | -0.037 |
| TG 44:2 | 0.917 | 0.132 | 0.272 | 0.037 | -0.031 | 0.053 | 0.005 | -0.078 | 0.010 |
| TG 46:0 | 0.873 | -0.008 | 0.265 | 0.237 | 0.037 | 0.222 | -0.055 | 0.091 | 0.011 |
| TG 46:1 | 0.893 | 0.121 | 0.178 | 0.327 | 0.076 | -0.007 | 0.008 | 0.077 | -0.046 |
| TG 46:2 | 0.917 | 0.216 | 0.198 | 0.163 | -0.059 | 0.012 | -0.014 | -0.031 | -0.034 |
| TG 46:3 | 0.875 | 0.318 | 0.223 | 0.157 | -0.046 | -0.047 | 0.059 | -0.031 | -0.063 |
| TG 46:4 | 0.810 | 0.363 | 0.309 | 0.011 | 0.067 | 0.034 | 0.109 | -0.044 | -0.043 |
| TG 48:0 | 0.785 | -0.039 | 0.233 | 0.258 | 0.058 | 0.402 | -0.022 | 0.130 | -0.005 |
| TG 48:1 | 0.818 | 0.083 | 0.157 | 0.476 | 0.033 | 0.084 | -0.071 | 0.103 | -0.038 |
| TG 48:2 | 0.797 | 0.302 | 0.146 | 0.429 | 0.065 | 0.005 | -0.034 | 0.127 | -0.026 |
| TG 48:3 | 0.786 | 0.442 | 0.115 | 0.302 | -0.048 | -0.089 | -0.007 | -0.012 | -0.030 |
| TG 48:4 | 0.701 | 0.614 | 0.194 | 0.056 | -0.014 | -0.047 | -0.044 | -0.128 | -0.003 |
| TG 49:0 | 0.759 | 0.018 | 0.328 | 0.355 | 0.000 | 0.281 | 0.015 | 0.230 | 0.019 |
| TG 49:1 | 0.718 | 0.149 | 0.204 | 0.568 | -0.040 | 0.110 | -0.007 | 0.246 | -0.020 |
| TG 49:2 | 0.690 | 0.270 | 0.130 | 0.567 | -0.006 | 0.014 | 0.060 | 0.260 | -0.008 |
| TG 49:3 | 0.610 | 0.513 | 0.119 | 0.474 | 0.122 | -0.039 | 0.039 | 0.253 | -0.013 |
| TG 50:0 | 0.650 | 0.017 | 0.302 | 0.314 | 0.088 | 0.565 | 0.015 | 0.063 | 0.006 |
| TG 50:1 | 0.680 | 0.031 | 0.167 | 0.577 | -0.003 | 0.185 | -0.091 | 0.148 | -0.001 |
| TG 50:2 | 0.636 | 0.241 | 0.096 | 0.654 | 0.021 | 0.060 | -0.053 | 0.104 | -0.008 |
| TG 50:3 | 0.527 | 0.569 | 0.072 | 0.553 | 0.084 | -0.037 | 0.000 | 0.101 | -0.014 |
| TG 50:4 | 0.455 | 0.800 | 0.136 | 0.287 | -0.074 | -0.016 | -0.021 | 0.036 | -0.008 |
| TG 50:5 | 0.529 | 0.779 | 0.156 | 0.161 | 0.048 | -0.015 | -0.015 | 0.059 | -0.066 |
| TG 50:6 | 0.745 | 0.374 | 0.172 | 0.111 | 0.390 | -0.049 | 0.027 | 0.016 | -0.031 |
| TG 51:1 | 0.653 | 0.111 | 0.262 | 0.622 | -0.015 | 0.173 | -0.023 | 0.204 | -0.033 |
| TG 51:2 | 0.511 | 0.286 | 0.148 | 0.758 | -0.012 | 0.008 | 0.042 | 0.179 | -0.037 |
| TG 51:3 | 0.361 | 0.627 | 0.131 | 0.632 | 0.025 | 0.013 | 0.041 | 0.149 | 0.003 |
| TG 51:4 | 0.160 | 0.829 | 0.102 | 0.312 | -0.157 | 0.019 | 0.004 | 0.160 | 0.032 |
| TG 51:5 | 0.391 | 0.705 | 0.204 | 0.198 | 0.343 | -0.016 | 0.048 | 0.158 | 0.006 |
| TG 52:0 | 0.441 | 0.036 | 0.324 | 0.166 | 0.041 | 0.741 | 0.146 | -0.009 | -0.020 |
| TG 52:1 | 0.619 | 0.036 | 0.343 | 0.585 | 0.048 | 0.274 | -0.089 | 0.056 | -0.052 |
| TG 52:2 | 0.302 | 0.269 | 0.164 | 0.848 | 0.006 | 0.043 | -0.008 | -0.041 | 0.016 |
| TG 52:3 | 0.175 | 0.655 | 0.140 | 0.641 | 0.035 | 0.081 | -0.035 | -0.061 | 0.072 |
| TG 52:4 | 0.070 | 0.854 | 0.130 | 0.362 | 0.066 | 0.088 | -0.037 | -0.021 | 0.052 |
| TG 52:5 | 0.195 | 0.844 | 0.150 | 0.260 | 0.282 | 0.050 | -0.008 | 0.071 | -0.022 |
| TG 52:6 | 0.395 | 0.777 | 0.116 | 0.151 | 0.213 | 0.028 | -0.057 | 0.061 | -0.044 |
| TG 53:1 | 0.606 | 0.067 | 0.380 | 0.570 | -0.008 | 0.210 | 0.018 | 0.166 | -0.042 |
| TG 53:2 | 0.358 | 0.269 | 0.264 | 0.807 | 0.007 | 0.054 | 0.084 | 0.098 | -0.048 |
| TG 53:3 | 0.240 | 0.584 | 0.248 | 0.696 | 0.039 | 0.042 | 0.040 | 0.071 | 0.022 |
| TG 53:4 | 0.186 | 0.771 | 0.215 | 0.513 | 0.110 | 0.024 | 0.018 | 0.085 | 0.024 |
| TG 53:5 | 0.188 | 0.817 | 0.203 | 0.347 | 0.263 | -0.012 | 0.052 | 0.111 | 0.009 |
| TG 54:0 | 0.382 | 0.064 | 0.415 | 0.062 | -0.008 | 0.725 | 0.092 | -0.030 | 0.098 |
| TG 54:1 | 0.542 | 0.029 | 0.549 | 0.393 | 0.020 | 0.353 | -0.053 | -0.009 | -0.121 |
| TG 54:2 | 0.366 | 0.189 | 0.452 | 0.726 | 0.057 | 0.166 | 0.013 | -0.095 | -0.067 |
| TG 54:3 | -0.036 | 0.543 | 0.393 | 0.636 | -0.009 | 0.013 | 0.073 | -0.267 | -0.032 |
| TG 54:4 | -0.212 | 0.757 | 0.329 | 0.401 | 0.013 | -0.021 | 0.032 | -0.255 | 0.049 |
| TG 54:5 | -0.284 | 0.833 | 0.295 | 0.040 | -0.028 | -0.002 | -0.013 | -0.232 | 0.063 |
| TG 54:6 | -0.159 | 0.917 | 0.232 | 0.003 | 0.026 | 0.013 | 0.028 | -0.111 | -0.004 |
| TG 54:8 | 0.120 | 0.790 | 0.200 | -0.145 | 0.380 | 0.012 | -0.009 | 0.034 | -0.118 |
| TG 56:1 | 0.459 | 0.061 | 0.775 | 0.203 | 0.021 | 0.247 | -0.165 | -0.021 | 0.011 |
| TG 56:2 | 0.375 | 0.138 | 0.765 | 0.389 | 0.077 | 0.168 | 0.000 | -0.096 | -0.058 |
| TG 56:3 | 0.194 | 0.293 | 0.522 | 0.704 | 0.055 | 0.001 | 0.118 | -0.178 | -0.043 |
| TG 56:4 | 0.079 | 0.518 | 0.404 | 0.635 | 0.127 | 0.013 | 0.047 | -0.208 | -0.012 |
| TG 56:5 | 0.145 | 0.309 | 0.219 | 0.746 | 0.358 | 0.058 | 0.025 | -0.093 | -0.023 |
| TG 56:6 | 0.194 | 0.406 | 0.207 | 0.642 | 0.447 | 0.003 | 0.026 | -0.053 | -0.018 |
| TG 56:7 | 0.133 | 0.587 | 0.209 | 0.331 | 0.587 | 0.060 | -0.013 | -0.006 | 0.046 |
| TG 56:8 | 0.030 | 0.177 | 0.073 | 0.155 | 0.862 | 0.041 | 0.028 | 0.032 | -0.038 |
| TG 56:9 | 0.079 | 0.315 | 0.111 | -0.063 | 0.899 | 0.001 | -0.052 | 0.049 | -0.017 |
| TG 57:1 | 0.494 | 0.060 | 0.686 | 0.209 | 0.011 | 0.174 | 0.072 | 0.228 | 0.065 |
| TG 57:2 | 0.436 | 0.090 | 0.704 | 0.248 | 0.073 | 0.092 | 0.114 | 0.258 | -0.034 |
| TG 58:1 | 0.457 | 0.037 | 0.800 | 0.213 | 0.040 | 0.138 | -0.106 | 0.012 | 0.007 |
| TG 58:10 | -0.054 | 0.147 | 0.075 | -0.175 | 0.910 | 0.020 | -0.044 | -0.003 | 0.047 |
| TG 58:2 | 0.336 | 0.133 | 0.848 | 0.183 | 0.087 | 0.108 | -0.131 | -0.077 | 0.005 |
| TG 58:3 | 0.225 | 0.242 | 0.854 | 0.125 | 0.092 | 0.085 | -0.098 | -0.102 | -0.061 |
| TG 58:4 | 0.235 | 0.364 | 0.632 | 0.492 | 0.173 | 0.088 | -0.026 | -0.128 | 0.070 |
| TG 58:6 | 0.269 | 0.350 | 0.262 | 0.656 | 0.341 | -0.006 | 0.135 | -0.019 | -0.078 |
| TG 58:8 | -0.023 | 0.021 | 0.164 | 0.304 | 0.882 | -0.027 | 0.059 | -0.055 | 0.002 |
| TG 58:9 | -0.103 | -0.069 | 0.075 | 0.002 | 0.952 | -0.004 | 0.051 | -0.057 | 0.043 |
| TG 59:2 | 0.436 | 0.158 | 0.783 | 0.249 | 0.082 | 0.098 | 0.063 | 0.135 | 0.032 |
| TG 59:3 | 0.205 | 0.196 | 0.713 | 0.208 | 0.110 | 0.002 | 0.157 | 0.148 | -0.022 |
| TG 60:11 | 0.127 | 0.018 | 0.089 | 0.132 | 0.954 | 0.005 | 0.016 | 0.052 | 0.022 |
| TG 60:2 | 0.259 | 0.112 | 0.893 | 0.190 | 0.040 | 0.064 | -0.061 | -0.095 | -0.008 |
| TG 60:3 | 0.169 | 0.286 | 0.885 | 0.095 | 0.066 | 0.061 | -0.130 | -0.129 | 0.030 |
| TG 60:4 | 0.140 | 0.445 | 0.821 | 0.001 | 0.109 | 0.035 | -0.086 | -0.092 | 0.070 |
| TG 60:6 | -0.180 | -0.057 | 0.038 | -0.109 | 0.111 | -0.214 | -0.077 | 0.020 | 0.721 |
| TG 62:3 | 0.173 | 0.206 | 0.905 | 0.124 | 0.103 | -0.010 | 0.090 | -0.015 | 0.052 |
| TG 62:4 | 0.159 | 0.318 | 0.798 | 0.174 | 0.208 | -0.013 | 0.129 | 0.026 | 0.005 |
| TG 64:4 | 0.091 | 0.198 | 0.731 | 0.095 | 0.128 | -0.048 | 0.241 | 0.253 | 0.067 |

Table e-3: Adjusted values for top contributors to “PC5”
TG values were adjusted for the following covariates: Age, Sex, APOE status, body mass index (BMI), and total triglycerides. Units are normalized peak height.

DXGrp= Diagnostic group; CN= control; MCI= mild cognitive impairment; AD= Alzheimer’s disease

| Table e-3: Adjusted values for top contributors to “PC5” | | | | | | |
| --- | --- | --- | --- | --- | --- | --- |
| DXGrp | TG 56:8 | TG 56:9 | TG 58:8 | TG 58:9 | TG 58:10 | TG 60:11 |
| CN | 2.413 | 3.088 | 3.093 | 2.607 | 2.406 | 3.151 |
| CN | 2.52 | 2.812 | 2.352 | 2.076 | 1.962 | 2.894 |
| MCI | 2.476 | 2.149 | 2.168 | 1.967 | 1.71 | 2.687 |
| MCI | 2.389 | 2.804 | 2.698 | 2.881 | 2.427 | 2.659 |
| MCI | 2.197 | 3.094 | 2.677 | 2.101 | 2.864 | 2.575 |
| MCI | 2.382 | 2.452 | 2.713 | 2.416 | 1.891 | 2.528 |
| CN | 1.606 | 1.812 | 1.336 | 1.888 | 2.193 | 2.328 |
| MCI | 2.634 | 2.567 | 2.699 | 2.929 | 2.718 | 2.273 |
| MCI | 1.362 | 1.769 | 2.258 | 2.531 | 1.679 | 2.263 |
| MCI | 1.227 | 2.534 | 2.092 | 2.202 | 2.335 | 2.207 |
| AD | 2.522 | 1.7 | 2.117 | 2.139 | 1.921 | 2.166 |
| MCI | 1.494 | 1.606 | 2.338 | 1.916 | 1.892 | 2.161 |
| CN | 2.101 | 2.098 | 1.814 | 2.098 | 1.942 | 2.112 |
| MCI | 2.702 | 2.16 | 2.058 | 2.24 | 1.964 | 2.095 |
| CN | 1.04 | 1.054 | 1.643 | 1.599 | 1.494 | 2.08 |
| MCI | 1.827 | 1.492 | 0.69 | 0.99 | 1.272 | 2.004 |
| MCI | 1.639 | 1.725 | 2.017 | 1.416 | 1.334 | 1.973 |
| CN | 1.521 | 2.221 | 0.975 | 1.264 | 1.969 | 1.955 |
| CN | 2.478 | 1.527 | 1.768 | 1.637 | 1.344 | 1.891 |
| MCI | 0.937 | 1.786 | 1.139 | 0.186 | 0.562 | 1.853 |
| CN | 1.53 | 1.152 | 2.136 | 1.323 | 0.664 | 1.846 |
| MCI | 1.947 | 1.698 | 2.251 | 2.27 | 1.783 | 1.844 |
| CN | 1.308 | 1.253 | 1.778 | 1.454 | 1.427 | 1.829 |
| MCI | 2.273 | 1.789 | 2.404 | 2.66 | 1.976 | 1.823 |
| MCI | 1.958 | 1.861 | 1.859 | 2.118 | 1.713 | 1.801 |
| CN | 2.146 | 2.235 | 2.221 | 2.399 | 2.652 | 1.77 |
| CN | 1.782 | 1.522 | 1.688 | 1.187 | 1.166 | 1.767 |
| MCI | 1.344 | 1.312 | 1.385 | 1.383 | 1.196 | 1.718 |
| AD | 1.822 | 1.634 | 1.392 | 1.221 | 0.962 | 1.715 |
| AD | 1.346 | 2.227 | 1.46 | 1.939 | 1.905 | 1.694 |
| CN | 2.43 | 1.417 | 2.296 | 2.361 | 1.738 | 1.688 |
| AD | 2.19 | 0.735 | 2.039 | 2.052 | 0.968 | 1.679 |
| AD | 2.02 | 1.744 | 1.814 | 1.312 | 0.529 | 1.67 |
| MCI | 1.752 | 0.805 | 1.656 | 1.709 | 0.942 | 1.611 |
| MCI | 2.075 | 0.907 | 1.621 | 1.553 | 0.796 | 1.61 |
| CN | 1.774 | 1.219 | 1.717 | 1.41 | 1.235 | 1.608 |
| CN | 1.277 | 1.357 | 2.27 | 2.12 | 1.25 | 1.593 |
| MCI | 1.379 | 1.416 | 1.568 | 1.375 | 1.232 | 1.59 |
| AD | 1.963 | 1.343 | 1.106 | 1.644 | 1.895 | 1.583 |
| CN | 0.943 | 1.539 | 0.651 | 1.283 | 1.837 | 1.563 |
| MCI | 0.834 | 1.002 | 0.922 | 1.004 | 1.354 | 1.563 |
| CN | 1.581 | 0.908 | 0.929 | 0.957 | 0.715 | 1.545 |
| MCI | 1.064 | 1.818 | 1.24 | 0.806 | 1.422 | 1.54 |
| MCI | 0.776 | 1.386 | 1.428 | 0.416 | 0.724 | 1.537 |
| MCI | 0.889 | 0.914 | 1.715 | 1.566 | 1.491 | 1.528 |
| MCI | 1.887 | 1.505 | 1.702 | 1.503 | 0.871 | 1.517 |
| MCI | 1.274 | 1.665 | 1.745 | 1.819 | 1.878 | 1.497 |
| MCI | 0.979 | 1.231 | 0.854 | 0.813 | 1.139 | 1.492 |
| MCI | 1.635 | 0.82 | 1.659 | 2.135 | 1.254 | 1.471 |
| MCI | 1.097 | 1.042 | 1.21 | 0.67 | 0.824 | 1.47 |
| CN | 1.283 | 1 | 1.839 | 1.208 | 0.848 | 1.459 |
| MCI | 1.613 | 1.156 | 1.15 | 0.732 | 0.276 | 1.457 |
| CN | 0.799 | 0.534 | 0.89 | 0.63 | 0.631 | 1.429 |
| MCI | 1.188 | 0.86 | 1.126 | 1.497 | 1.315 | 1.426 |
| AD | 1.766 | 1.187 | 1.614 | 1.968 | 1.403 | 1.412 |
| CN | 1.589 | 1.631 | 1.736 | 1.391 | 1.163 | 1.412 |
| CN | 0.937 | 1.117 | 1.223 | 1.037 | 0.988 | 1.392 |
| MCI | 1.75 | 0.971 | 1.497 | 1.532 | 0.94 | 1.378 |
| CN | 1.573 | 1.552 | 1.26 | 1.701 | 1.658 | 1.377 |
| MCI | 1.748 | 1.663 | 1.515 | 1.298 | 1.098 | 1.371 |
| MCI | 1.401 | 1.223 | 1.347 | 1.045 | 0.993 | 1.363 |
| CN | 1.923 | 0.914 | 2.249 | 1.993 | 0.992 | 1.358 |
| AD | 1.036 | 1.061 | 0.692 | 1.259 | 1.014 | 1.351 |
| CN | 1.634 | 1.073 | 1.453 | 1.209 | 0.453 | 1.339 |
| MCI | 1.314 | 0.751 | 1.094 | 0.836 | 0.928 | 1.336 |
| CN | 1.27 | 0.519 | 1.023 | 1.266 | 0.725 | 1.333 |
| AD | 0.975 | 0.904 | 1.139 | 0.854 | 0.645 | 1.332 |
| CN | 0.832 | 0.324 | 0.916 | 0.707 | -0.022 | 1.331 |
| AD | 1.794 | 1.261 | 1.151 | 1.157 | 1.027 | 1.316 |
| MCI | 1.508 | 1.703 | 0.98 | 1.345 | 1.507 | 1.307 |
| CN | 0.88 | 1.229 | 1.046 | 0.962 | 1.12 | 1.294 |
| CN | 1.366 | 0.847 | 1.198 | 0.684 | 0.374 | 1.277 |
| CN | 1.581 | 1.044 | 0.899 | 0.554 | 0.506 | 1.268 |
| CN | 0.826 | 1.087 | 1.194 | 1.188 | 0.819 | 1.267 |
| CN | 1.754 | 1.341 | 1.984 | 1.963 | 1.145 | 1.262 |
| CN | 1.239 | 0.548 | 0.456 | 0.565 | 0.814 | 1.21 |
| AD | 0.927 | 2.268 | 1.275 | 1.34 | 1.884 | 1.191 |
| MCI | 1.142 | 0.969 | 0.679 | 1.119 | 1.186 | 1.169 |
| CN | 0.405 | 0.51 | 0.701 | 1.26 | 1.376 | 1.143 |
| CN | 0.942 | 0.991 | 0.896 | 0.175 | 0.272 | 1.128 |
| MCI | 0.866 | 0.59 | 0.911 | 0.797 | 0.47 | 1.122 |
| CN | 1.045 | 0.912 | 1.764 | 0.58 | 0.216 | 1.111 |
| CN | 0.999 | 0.534 | 0.78 | 0.401 | 0.284 | 1.089 |
| MCI | 1.002 | 0.007 | 1.049 | 0.527 | -0.002 | 1.087 |
| AD | 1.132 | 0.825 | 1.522 | 1.371 | 1.117 | 1.085 |
| AD | 0.944 | 1.007 | 1.402 | 1.002 | 0.66 | 1.079 |
| CN | 1.079 | 0.87 | 0.679 | 1.31 | 1.312 | 1.078 |
| MCI | 1.181 | 0.304 | 1.097 | 0.755 | 0.152 | 1.068 |
| CN | 0.581 | 0.628 | 0.658 | 0.567 | 1.131 | 1.053 |
| AD | 1.384 | 0.951 | 0.948 | 1.443 | 1.184 | 1.037 |
| MCI | 1.827 | 1.199 | 1.721 | 2.141 | 1.845 | 1.003 |
| MCI | 0.287 | 0.409 | 1.008 | 1.279 | 1.331 | 1.001 |
| MCI | 1.114 | 0.798 | 1.007 | 0.882 | 0.754 | 0.992 |
| MCI | 1.084 | 0.82 | 1.121 | 0.413 | -0.1 | 0.99 |
| MCI | 0.962 | 0.698 | 1.489 | 1.169 | 0.912 | 0.987 |
| AD | 0.459 | 1.152 | 0.155 | 0.271 | 0.89 | 0.985 |
| CN | 0.926 | 0.388 | 1.299 | 0.869 | 0.419 | 0.982 |
| MCI | 0.103 | 0.542 | 0.776 | 0.72 | 0.772 | 0.975 |
| CN | -0.492 | 0.145 | 0.27 | 0.996 | 0.992 | 0.962 |
| CN | 1.168 | 0.44 | 0.729 | 0.672 | 0.535 | 0.961 |
| CN | 0.325 | -0.86 | 0.678 | 0.941 | 0.305 | 0.961 |
| CN | 1.531 | 0.439 | 0.758 | 0.741 | 0.491 | 0.959 |
| CN | 0.42 | 0.655 | 0.609 | 1.112 | 1.201 | 0.958 |
| MCI | 1.048 | 0.587 | 0.809 | 0.644 | 0.671 | 0.955 |
| CN | 1.343 | 0.783 | 0.981 | 0.863 | 0.699 | 0.953 |
| MCI | 0.113 | -0.896 | 0.825 | 0.83 | 0.182 | 0.947 |
| AD | 1.335 | 0.655 | 0.701 | 0.252 | -0.012 | 0.945 |
| MCI | 0.79 | 1.1 | 0.758 | 0.658 | 0.768 | 0.944 |
| CN | 1.103 | 1.02 | 0.878 | 1.119 | 1.055 | 0.93 |
| CN | 0.023 | 0.747 | 0.613 | 0.968 | 1.241 | 0.92 |
| AD | 1.962 | 1.115 | 1.122 | 1.103 | 1.045 | 0.913 |
| CN | 1.074 | 2.357 | 0.299 | 0.376 | 1.255 | 0.911 |
| CN | 0.939 | 0.6 | 0.797 | 0.638 | 0.433 | 0.907 |
| CN | 1.161 | 0.385 | 0.91 | 0.907 | 0.322 | 0.89 |
| CN | 0.476 | 1 | 0.56 | 0.326 | 0.304 | 0.888 |
| MCI | 0.892 | 0.787 | 0.857 | 0.87 | 0.822 | 0.886 |
| AD | 0.844 | 0.982 | 1.095 | 1.095 | 1.104 | 0.878 |
| AD | 1.357 | 0.142 | 1.713 | 1.349 | 0.399 | 0.873 |
| AD | 0.736 | 0.656 | 0.033 | -0.167 | 0.24 | 0.861 |
| AD | 0.417 | 1.022 | 1.019 | 0.885 | 1.07 | 0.858 |
| CN | 1.009 | 1.011 | 1.004 | 0.461 | 0.413 | 0.836 |
| CN | 0.903 | 1.387 | 0.768 | 0.281 | 0.682 | 0.803 |
| AD | 1.25 | 0.55 | 0.816 | 1.257 | 1.156 | 0.8 |
| MCI | 0.847 | 0.688 | 0.853 | 0.668 | 0.689 | 0.8 |
| CN | 1.189 | 0.385 | 0.803 | 0.354 | 0.046 | 0.799 |
| AD | 0.286 | 1.139 | 0.737 | 1.195 | 1.181 | 0.792 |
| CN | 0.764 | 0.571 | 0.553 | 0.585 | 0.661 | 0.787 |
| CN | 0.491 | 0.807 | 0.617 | 0.192 | 0.195 | 0.785 |
| CN | 1.358 | 0.673 | 1.009 | 0.625 | 0.198 | 0.778 |
| MCI | 0.329 | 1.067 | 1.212 | 1.339 | 1.379 | 0.773 |
| AD | 0.231 | -0.153 | 0.637 | -0.173 | 0.064 | 0.771 |
| MCI | 0.021 | 0.471 | 0.791 | 0.639 | 0.695 | 0.767 |
| CN | 0.709 | 0.853 | 0.874 | 0.481 | 0.519 | 0.766 |
| MCI | -0.119 | -0.195 | 1.136 | 0.361 | 0.065 | 0.765 |
| MCI | 0.927 | 0.532 | 0.962 | 0.881 | 0.318 | 0.764 |
| MCI | 0.937 | 0.335 | 1.031 | 0.271 | -0.176 | 0.764 |
| MCI | 0.73 | 0.488 | 0.205 | 0.814 | 1.219 | 0.756 |
| AD | 0.818 | 0.123 | 0.931 | 0.533 | 0.07 | 0.749 |
| MCI | 1.491 | 1.774 | 0.407 | 0.754 | 1.485 | 0.748 |
| CN | -0.031 | 0.493 | -0.523 | 0.461 | 1.146 | 0.743 |
| CN | 0.654 | 0.234 | 0.394 | -0.245 | -0.534 | 0.735 |
| CN | -0.004 | 0.657 | 0.435 | 0.524 | 0.704 | 0.725 |
| CN | 0.489 | -0.192 | 1.11 | 0.718 | -0.503 | 0.722 |
| MCI | 0.711 | 0.104 | 1.343 | 0.636 | -0.013 | 0.716 |
| AD | 1.325 | 0.34 | 0.802 | 0.41 | 0.227 | 0.711 |
| MCI | 1.116 | 0.693 | 0.538 | 0.583 | 0.808 | 0.71 |
| MCI | 0.605 | 0.702 | 0.842 | 1.06 | 1.153 | 0.706 |
| AD | 0.395 | 1.206 | 0.042 | 0.754 | 1.53 | 0.702 |
| AD | 1.377 | 0.624 | 0.582 | 0.16 | -0.035 | 0.701 |
| MCI | 0.813 | 1.014 | 0.498 | 0.904 | 0.971 | 0.697 |
| MCI | 1.116 | 0.348 | 0.221 | -0.087 | 0.002 | 0.684 |
| MCI | 0.498 | 0.504 | 0.758 | 0.575 | 0.387 | 0.683 |
| CN | 1.025 | 0.365 | 1.136 | 0.723 | 0.338 | 0.683 |
| MCI | 1.071 | 0.477 | 0.261 | 0.342 | 0.598 | 0.681 |
| AD | 0.396 | 1.658 | 0.028 | 0.748 | 1.584 | 0.655 |
| MCI | 1.125 | 0.753 | 0.728 | 0.93 | 0.988 | 0.655 |
| MCI | 0.821 | 0.612 | 0.577 | 0.214 | -0.029 | 0.645 |
| CN | 0.9 | 0.512 | 0.362 | 0.244 | 0.322 | 0.643 |
| MCI | -0.082 | 1.579 | -0.087 | 0.03 | 0.723 | 0.625 |
| AD | -0.106 | 0.473 | 0.104 | 1.13 | 1.363 | 0.624 |
| AD | 0.616 | 1.016 | 0.061 | 0.498 | 0.927 | 0.622 |
| CN | 0.821 | 0.58 | 0.381 | 0.69 | 1.245 | 0.62 |
| MCI | 0.009 | 0.638 | 0.12 | 0.789 | 1.477 | 0.618 |
| CN | 0.687 | 0.119 | 0.447 | 0.03 | -0.222 | 0.616 |
| MCI | 0.855 | 0.091 | 0.766 | 0.689 | 0.341 | 0.615 |
| MCI | 0.373 | 0.512 | 0.095 | 0.25 | 0.624 | 0.613 |
| CN | 0.353 | -0.049 | 0.512 | 0.334 | -0.076 | 0.61 |
| MCI | 0.25 | -0.355 | 0.459 | 0.079 | -0.232 | 0.61 |
| AD | 0.176 | 0.528 | 0.721 | 0.801 | 0.861 | 0.605 |
| MCI | 0.642 | 0.609 | 0.7 | 0.647 | 0.632 | 0.603 |
| MCI | 0.345 | 0.247 | 0.538 | 0.618 | 0.56 | 0.598 |
| MCI | 0.165 | 1.084 | 0.655 | 0.159 | 0.574 | 0.597 |
| AD | 0.76 | -0.184 | 0.726 | 1.189 | 0.516 | 0.597 |
| CN | 0.629 | 0.74 | 1.34 | 1.428 | 0.821 | 0.591 |
| AD | 0.522 | 0.594 | 0.451 | 0.28 | 0.284 | 0.589 |
| MCI | 0.621 | -0.014 | 1.076 | 0.885 | -0.024 | 0.586 |
| AD | 0.832 | 0.726 | 0.335 | 0.054 | 0.087 | 0.583 |
| MCI | 0.838 | 0.272 | 0.744 | 0.415 | 0.143 | 0.582 |
| MCI | 0.797 | -0.516 | 0.512 | 0.184 | -0.581 | 0.57 |
| CN | 0.086 | -0.703 | 0.564 | 0.573 | 0.114 | 0.566 |
| MCI | 0.175 | 0.519 | -0.349 | 0.25 | 0.691 | 0.565 |
| MCI | 0.539 | 1.064 | -0.051 | 0.212 | 0.853 | 0.564 |
| MCI | 0.754 | 0.623 | 0.484 | 0.526 | 0.534 | 0.564 |
| AD | 0.529 | 1.389 | 0.096 | 0.283 | 0.477 | 0.556 |
| MCI | 0.313 | 0.433 | 0.643 | 0.634 | 0.979 | 0.555 |
| MCI | 0.264 | 0.233 | -0.061 | 0.268 | 0.615 | 0.55 |
| MCI | 0.26 | 2.222 | -0.161 | 1.518 | 2.531 | 0.525 |
| MCI | 0.956 | 2.134 | 0.39 | 0.782 | 1.38 | 0.523 |
| AD | 0.484 | 0.165 | 0.351 | 0.647 | 0.305 | 0.517 |
| MCI | 0.384 | 0.551 | 0.444 | 0.01 | 0.267 | 0.514 |
| CN | 0.331 | 0.443 | 0.814 | 0.745 | 0.218 | 0.51 |
| CN | 0.249 | 0.181 | 0.221 | 0.364 | 0.352 | 0.506 |
| MCI | 0.033 | 0.834 | -0.202 | 0.096 | 0.65 | 0.505 |
| MCI | -0.009 | 0.364 | 0.771 | 0.768 | 0.739 | 0.499 |
| AD | 0.492 | 0.464 | 0.43 | 0.382 | 0.42 | 0.496 |
| MCI | -0.097 | 0.559 | 0.529 | 0.466 | 0.528 | 0.494 |
| AD | -0.336 | 0.569 | 0.267 | 0.458 | 0.488 | 0.492 |
| CN | 0.322 | 0.823 | 0.078 | 0.152 | 0.674 | 0.491 |
| MCI | -0.289 | 0.613 | 0.131 | 1.271 | 1.058 | 0.483 |
| MCI | -0.427 | 0.076 | 0.988 | 0.731 | 0.476 | 0.483 |
| AD | 0.74 | -0.039 | 0.507 | 0.291 | 0.309 | 0.48 |
| AD | 0.997 | 0.186 | 0.773 | 0.738 | 0.094 | 0.479 |
| CN | 0.174 | 0.463 | 0.646 | 0.597 | 0.626 | 0.476 |
| MCI | 0.499 | 0.788 | 0.306 | 0.279 | 0.422 | 0.472 |
| MCI | 0.396 | 0.828 | 0.502 | 1.198 | 1.17 | 0.464 |
| MCI | 0.027 | 0.174 | 0.069 | 0.156 | 0.393 | 0.459 |
| MCI | 1.405 | 0.761 | 0.565 | 0.887 | 1.375 | 0.454 |
| MCI | -0.533 | 0.356 | 0.375 | 0.36 | 0.497 | 0.444 |
| CN | 0.513 | 0.656 | 1.128 | 1.924 | 1.662 | 0.439 |
| MCI | 0.461 | -0.426 | 0.5 | 0.424 | -0.325 | 0.438 |
| MCI | 0.816 | 0.131 | 0.473 | 0.116 | -0.22 | 0.434 |
| MCI | -0.11 | 0.174 | 0.215 | 0.356 | 0.519 | 0.433 |
| CN | 0.09 | -0.187 | -0.44 | 0.304 | 0.556 | 0.429 |
| CN | 1.223 | 0.099 | 1.32 | 1.558 | 0.844 | 0.424 |
| MCI | 0.483 | -0.471 | 0.108 | 0.029 | 0.19 | 0.419 |
| CN | 0.296 | 0.085 | 0.546 | 0.87 | 0.706 | 0.417 |
| CN | 0.908 | 0.333 | 0.891 | 0.545 | 0.331 | 0.414 |
| MCI | 0.435 | 0.338 | 0.507 | 0.715 | 0.523 | 0.402 |
| CN | 0.324 | 0.004 | 0.619 | 0.215 | -0.145 | 0.397 |
| CN | 0.13 | 0.085 | 0.458 | 0.199 | -0.05 | 0.396 |
| CN | 0.322 | 0.55 | 0.291 | 0.363 | 0.427 | 0.388 |
| CN | 0.085 | 0.425 | 0.169 | 0.253 | 0.638 | 0.385 |
| MCI | 0.531 | 0.796 | 0.49 | 0.346 | 0.356 | 0.373 |
| MCI | 0.106 | 0.915 | 0.431 | 0.241 | 0.615 | 0.372 |
| MCI | 0.101 | 0.501 | 0.244 | 0.24 | 0.27 | 0.372 |
| CN | 0.511 | 0.442 | 0.51 | 0.137 | -0.152 | 0.37 |
| MCI | 0.11 | 0.671 | 0.062 | 0.325 | 0.614 | 0.369 |
| CN | -0.061 | 0.912 | 0.246 | -0.118 | 0.23 | 0.36 |
| AD | 0.418 | 0.157 | 0.232 | 0.451 | 0.384 | 0.353 |
| MCI | 0.195 | 0.073 | 0.486 | -0.035 | 0.069 | 0.351 |
| AD | 0.447 | 0.171 | -0.07 | 0.661 | 1.151 | 0.346 |
| CN | 0.237 | 0.198 | 0.138 | 0.162 | 0.081 | 0.34 |
| MCI | -0.192 | 0.946 | -0.203 | 0.592 | 1.069 | 0.338 |
| MCI | -0.033 | 0.1 | 0.237 | 0.36 | 0.355 | 0.333 |
| AD | 0.54 | 0.47 | 0.112 | 0.254 | 0.351 | 0.333 |
| MCI | 0.619 | 0.411 | 0.548 | 0.585 | 0.243 | 0.329 |
| MCI | 0.049 | 0.429 | 0.4 | 0.162 | 0.234 | 0.326 |
| AD | -0.11 | -0.182 | 0.342 | -0.298 | -0.118 | 0.323 |
| MCI | 0.039 | 0.468 | 0.041 | 0.114 | 0.502 | 0.32 |
| MCI | 0.675 | 1.207 | 1.647 | 2.159 | 1.414 | 0.307 |
| MCI | -0.248 | 0.911 | -0.439 | 0.608 | 1.13 | 0.305 |
| CN | -0.212 | 1.363 | -0.038 | 0.132 | 0.761 | 0.301 |
| MCI | 0.251 | 0.665 | 0.03 | -0.179 | 0.028 | 0.3 |
| MCI | 0.137 | 0.212 | 0.286 | 0.177 | 0.371 | 0.293 |
| CN | 0.644 | 0.595 | 0.286 | 0.618 | 0.589 | 0.29 |
| MCI | 0.449 | 1.085 | -0.299 | 0.167 | 0.817 | 0.281 |
| AD | -0.384 | -0.371 | -0.33 | 0.207 | 0.028 | 0.269 |
| CN | -0.03 | -0.05 | -0.026 | 0.279 | -0.026 | 0.269 |
| MCI | -0.417 | 0.875 | -0.254 | 1.119 | 1.364 | 0.26 |
| MCI | 0.329 | 0.581 | 0.437 | 0.081 | 0.171 | 0.26 |
| AD | -0.486 | -0.047 | -0.008 | 0.053 | 0.057 | 0.254 |
| AD | -0.131 | 0.144 | 0.457 | -0.303 | -0.046 | 0.254 |
| AD | -0.09 | -0.475 | -0.13 | 0.093 | 0.419 | 0.253 |
| CN | 1.035 | 0.339 | 0.507 | 0.127 | -0.314 | 0.252 |
| CN | 0.81 | 0.351 | 0.666 | 0.501 | 0.112 | 0.248 |
| AD | 0.592 | 0.334 | 0.578 | 0.351 | 0.067 | 0.235 |
| AD | -0.117 | 0.856 | 0.15 | 0.637 | 1.12 | 0.231 |
| MCI | 0.211 | 0.513 | 0.119 | 0.089 | 0.516 | 0.219 |
| AD | 0.167 | 1.281 | 0.146 | 0.241 | 0.608 | 0.204 |
| MCI | -0.239 | -0.329 | 0.208 | 0.153 | -0.186 | 0.201 |
| MCI | 0.06 | -0.098 | 0.822 | 0.51 | 0.036 | 0.197 |
| AD | 0.172 | -0.083 | 0.319 | -0.066 | -0.249 | 0.197 |
| CN | 1.054 | 0.449 | 0.7 | 0.067 | 0.116 | 0.196 |
| CN | 0.278 | -0.279 | -0.345 | -0.088 | -0.113 | 0.194 |
| AD | 0.335 | -0.201 | 0.02 | -0.112 | -0.383 | 0.189 |
| MCI | 0.086 | 0.458 | -0.318 | -0.089 | 0.191 | 0.188 |
| MCI | -0.195 | 0.998 | -1.037 | 0.389 | 1.052 | 0.186 |
| MCI | 0.303 | 0.334 | 0.399 | -0.057 | 0.012 | 0.181 |
| AD | 0.637 | -0.016 | -0.056 | -0.087 | -0.222 | 0.181 |
| MCI | -0.414 | -0.376 | -0.602 | -0.963 | -0.81 | 0.176 |
| CN | -0.205 | 0.438 | -0.209 | 0.379 | 0.808 | 0.168 |
| CN | -0.202 | -0.694 | -0.013 | -0.111 | -0.295 | 0.163 |
| MCI | 0.229 | -0.276 | 0.243 | 0.091 | -0.434 | 0.158 |
| CN | 0.067 | 0.602 | 0.04 | -0.067 | 0.249 | 0.155 |
| MCI | 0.206 | 0.224 | 0.442 | -0.341 | -0.253 | 0.154 |
| AD | -0.591 | -0.193 | 0.143 | 0.8 | 0.83 | 0.15 |
| MCI | 1.097 | 0.319 | 0.834 | 0.435 | 0.076 | 0.149 |
| AD | -0.437 | 0.808 | -0.226 | 0.278 | 0.876 | 0.141 |
| AD | -0.23 | -0.061 | -0.119 | 0.103 | -0.028 | 0.141 |
| MCI | 0.482 | 0.15 | 0.198 | -0.388 | -0.415 | 0.141 |
| MCI | 1.011 | -0.141 | 1.022 | 0.793 | 0.067 | 0.139 |
| CN | 0.986 | 0.76 | 0.398 | -0.019 | -0.333 | 0.139 |
| MCI | 0.336 | -0.697 | 0.003 | 0.105 | -0.28 | 0.138 |
| CN | 0.727 | 0.126 | 0.589 | 0.161 | -0.316 | 0.136 |
| MCI | -0.247 | -0.021 | 0.149 | -0.189 | -0.175 | 0.134 |
| CN | 0.744 | 0.306 | 0.933 | 0.709 | 0.245 | 0.133 |
| AD | 0.255 | -0.172 | -0.293 | -0.338 | -0.346 | 0.131 |
| MCI | 0.547 | 0.307 | 0.591 | 0.281 | -0.56 | 0.131 |
| AD | -0.212 | 0.353 | 0.218 | 0.174 | 0.316 | 0.13 |
| CN | 0.052 | 0.058 | 0.36 | -0.003 | 0.15 | 0.127 |
| MCI | -0.104 | -0.019 | -0.009 | 0.026 | -0.038 | 0.127 |
| MCI | 0.018 | 0.526 | 0.335 | -0.28 | 0.037 | 0.126 |
| CN | -0.209 | -0.493 | 0.139 | 0.066 | 0.064 | 0.125 |
| CN | -0.028 | 0.739 | 0.229 | 0.799 | 1.176 | 0.123 |
| MCI | -0.063 | 0.56 | -0.157 | -0.052 | 0.554 | 0.118 |
| AD | 0.581 | 1.126 | 0.054 | 0.491 | 0.882 | 0.117 |
| MCI | 0.023 | -0.728 | 0.259 | 0.004 | -0.637 | 0.115 |
| MCI | -0.19 | 0.368 | 0.125 | 0.244 | 0.751 | 0.114 |
| CN | 0.751 | 0.887 | 0.632 | 0.115 | 0.324 | 0.11 |
| AD | -0.957 | -0.736 | 0.092 | -0.004 | -0.285 | 0.102 |
| MCI | 0.365 | -0.102 | -0.387 | -0.33 | 0.413 | 0.1 |
| CN | 0.369 | -0.106 | 0.427 | 0.306 | -0.161 | 0.1 |
| MCI | -0.277 | 0.464 | -0.548 | -0.012 | 0.423 | 0.098 |
| AD | 0.286 | 0.239 | 0.394 | 0.13 | 0.123 | 0.097 |
| CN | 0.526 | -0.268 | 0.669 | 0.13 | -0.933 | 0.097 |
| MCI | 0.093 | -0.551 | 0.365 | 0.206 | -0.362 | 0.092 |
| MCI | 0.102 | 0.354 | -0.189 | -0.048 | 0.437 | 0.091 |
| CN | 0.498 | 0.132 | -0.206 | 0.039 | 0.637 | 0.089 |
| AD | -0.409 | 0.523 | 0.339 | 0.609 | 0.648 | 0.088 |
| MCI | -0.406 | -0.307 | -0.051 | 0.268 | -0.159 | 0.088 |
| CN | 0.249 | -0.034 | 0.777 | 0.352 | -0.243 | 0.088 |
| CN | -0.262 | 0.401 | 0.087 | -0.103 | -0.297 | 0.081 |
| AD | 0.988 | 1.446 | 0.019 | -0.592 | 0.151 | 0.08 |
| MCI | -0.764 | 0.076 | 0.261 | 0.143 | 0.031 | 0.08 |
| MCI | 0.375 | -0.603 | 0.213 | -0.162 | -0.962 | 0.08 |
| MCI | -0.306 | 0.235 | -0.312 | 0.078 | 0.571 | 0.078 |
| CN | 0.333 | -0.615 | 0.242 | -0.438 | -1.541 | 0.063 |
| AD | -0.295 | 0.186 | -0.237 | 0.183 | 0.526 | 0.062 |
| CN | 0.354 | -0.788 | -0.016 | -0.214 | -1.061 | 0.062 |
| MCI | 0.149 | 0.371 | -0.39 | -0.159 | 0.183 | 0.059 |
| MCI | -0.254 | -0.055 | -0.191 | -0.377 | 0.195 | 0.056 |
| CN | 0.168 | -0.102 | 0.211 | -0.16 | -0.344 | 0.056 |
| CN | -0.604 | 0.093 | -0.113 | 0.251 | 0.507 | 0.051 |
| MCI | -0.948 | 0.251 | -0.468 | 0.576 | 0.851 | 0.048 |
| CN | 0.017 | 0.164 | -0.158 | 0.148 | 0.785 | 0.043 |
| AD | 0.121 | -0.531 | 0.027 | 0.075 | 0.054 | 0.042 |
| CN | -0.506 | -0.309 | 0.057 | 0.248 | 0.061 | 0.04 |
| MCI | 0.752 | 0.882 | 0.369 | 0.977 | 1.497 | 0.034 |
| CN | -0.847 | 0.263 | 0.122 | 0.361 | 0.519 | 0.032 |
| AD | 0.043 | -0.147 | 0.085 | -0.116 | -0.061 | 0.027 |
| MCI | 0.199 | -0.455 | -0.441 | -0.442 | -0.739 | 0.02 |
| CN | -0.086 | -0.155 | 0.166 | 0.05 | -0.159 | 0.019 |
| CN | 0.213 | -0.177 | -0.069 | 0.017 | 0.043 | 0.012 |
| MCI | -0.71 | -0.2 | -0.215 | 0.459 | 0.37 | 0.008 |
| MCI | 0.505 | -0.18 | 0.221 | -0.273 | -0.609 | 0.005 |
| MCI | 0.167 | -0.382 | 0.361 | -0.252 | -0.645 | 0.005 |
| MCI | -0.097 | 0.444 | 0.283 | 0.629 | 0.889 | 0 |
| MCI | 0.212 | 0.562 | 0.55 | 0.625 | 0.597 | -0.011 |
| MCI | -0.088 | -0.423 | 0.325 | -0.116 | -0.825 | -0.011 |
| CN | 0.123 | -0.979 | 0.177 | 0.037 | -1.129 | -0.012 |
| AD | -0.082 | 0.075 | -0.251 | -0.359 | 0.22 | -0.023 |
| MCI | -0.077 | -0.317 | -0.021 | 0.121 | 0.255 | -0.024 |
| MCI | 0.153 | 0.778 | -0.367 | -0.065 | 0.593 | -0.027 |
| CN | -0.782 | -0.493 | -0.425 | 0.028 | 0.144 | -0.028 |
| CN | 0.384 | -0.126 | 0.675 | 0.524 | -0.546 | -0.031 |
| CN | -0.725 | -0.081 | -0.293 | -0.261 | 0.13 | -0.033 |
| CN | -0.496 | 0.154 | 0.1 | -0.036 | 0.153 | -0.034 |
| AD | -0.35 | -0.177 | -0.281 | 0.649 | 1.019 | -0.039 |
| MCI | 0.216 | 0.018 | 0.144 | -0.078 | -0.403 | -0.04 |
| MCI | -0.505 | -0.069 | -0.086 | 0.469 | 0.217 | -0.043 |
| AD | -0.535 | -0.382 | -0.026 | -0.281 | -0.696 | -0.044 |
| MCI | 0.879 | 0.084 | 0.196 | 0.358 | -0.638 | -0.046 |
| MCI | -0.065 | -0.096 | -0.307 | -0.082 | 0.039 | -0.05 |
| MCI | -0.102 | -0.06 | 0.369 | 0.175 | -0.009 | -0.05 |
| MCI | 0.055 | -0.23 | 0.12 | 0.109 | -0.345 | -0.052 |
| MCI | 0.055 | -0.375 | 0.006 | -0.068 | 0.405 | -0.063 |
| CN | -0.901 | 0.402 | -0.394 | 0.236 | 0.372 | -0.064 |
| CN | -0.187 | -0.062 | -0.751 | -0.051 | 0.454 | -0.068 |
| MCI | -0.203 | -0.436 | -0.527 | -0.805 | -0.427 | -0.068 |
| AD | -0.139 | -0.688 | 0.225 | -0.591 | -1.11 | -0.076 |
| MCI | -0.252 | -0.533 | 0.088 | -0.653 | -0.838 | -0.08 |
| MCI | -1.597 | -1.464 | 0.045 | -0.51 | -0.814 | -0.084 |
| MCI | -0.408 | -0.065 | -0.352 | -0.009 | 0.329 | -0.086 |
| MCI | 0.245 | -0.247 | 0.256 | 0.32 | 0.031 | -0.087 |
| AD | -0.344 | 0.292 | 0.106 | -0.301 | -0.471 | -0.088 |
| MCI | 0.733 | 0.544 | 0.395 | 0.185 | 0.374 | -0.091 |
| MCI | 0.825 | -0.244 | 0.382 | 0.787 | 0.167 | -0.094 |
| AD | -0.521 | -0.118 | -0.031 | 0.036 | 0.131 | -0.096 |
| MCI | 0.199 | 0.137 | -0.492 | -0.186 | 0.309 | -0.098 |
| CN | -0.26 | 0.395 | -0.5 | 0.26 | 0.658 | -0.103 |
| MCI | -0.493 | -0.819 | -0.46 | 0.209 | 0.23 | -0.103 |
| MCI | -0.125 | 0.526 | -0.103 | 0.124 | 0.507 | -0.107 |
| AD | -0.173 | 0.307 | 0.153 | -0.284 | 0.062 | -0.107 |
| MCI | -0.459 | -0.215 | -0.126 | -1.052 | -1.389 | -0.107 |
| MCI | 0.439 | -0.457 | 0.401 | 0.27 | -0.629 | -0.108 |
| AD | -0.025 | 0.53 | 0.02 | 0.628 | 0.604 | -0.113 |
| AD | -0.671 | -0.304 | 0.236 | 0.03 | -0.142 | -0.117 |
| CN | -0.776 | -0.243 | -0.256 | 0.076 | 0.509 | -0.118 |
| MCI | -0.104 | 0.128 | -0.256 | -0.247 | -0.104 | -0.118 |
| MCI | 0.181 | 0.127 | 0.018 | -0.13 | 0.011 | -0.119 |
| AD | -0.029 | -0.406 | 0.319 | -0.208 | 0.03 | -0.126 |
| MCI | -0.01 | -0.776 | -0.122 | -0.086 | -0.275 | -0.126 |
| MCI | -0.75 | 0.749 | -0.486 | 0.895 | 1.207 | -0.128 |
| CN | -0.447 | -0.003 | -0.7 | -0.037 | 0.363 | -0.133 |
| MCI | 0.017 | -0.408 | 0.553 | 1.066 | -0.066 | -0.135 |
| CN | -0.223 | -0.008 | -0.697 | 0.068 | 0.429 | -0.145 |
| MCI | -0.782 | -1.024 | -0.263 | -0.71 | -0.952 | -0.146 |
| MCI | -1.076 | -0.704 | -0.444 | -0.394 | -0.272 | -0.151 |
| AD | 0.405 | -0.395 | -0.312 | -0.364 | -0.267 | -0.153 |
| MCI | -0.193 | -0.07 | -0.059 | -0.022 | -0.135 | -0.157 |
| AD | 0.133 | 0.654 | -0.323 | -0.378 | 0.285 | -0.159 |
| MCI | 0.313 | -0.355 | 0.23 | 0.02 | -0.681 | -0.16 |
| AD | -0.425 | -0.037 | 0.249 | 0.188 | 0.018 | -0.161 |
| AD | -0.083 | 0.132 | -0.028 | -0.384 | -0.555 | -0.167 |
| MCI | 0.416 | -0.355 | -0.218 | -0.278 | -0.629 | -0.167 |
| MCI | -0.359 | -0.237 | 0.193 | -0.34 | -0.501 | -0.171 |
| AD | -0.718 | -0.213 | 0.29 | -0.25 | -0.351 | -0.172 |
| MCI | 0.341 | -0.415 | 0.051 | 0.038 | 0.204 | -0.174 |
| MCI | 1.158 | 0.181 | 0.544 | 0.767 | 0.576 | -0.176 |
| CN | -0.374 | -0.401 | 0.014 | -0.074 | 0.035 | -0.178 |
| AD | 0.029 | -0.837 | -0.114 | -0.318 | -0.506 | -0.179 |
| MCI | 0.153 | -0.496 | 0.196 | -0.011 | -0.329 | -0.183 |
| AD | 0.082 | -0.082 | -0.051 | -0.201 | -0.411 | -0.185 |
| CN | 0.131 | -0.541 | -0.065 | -0.019 | 0.549 | -0.19 |
| MCI | 0.521 | -0.032 | -0.083 | -0.701 | -1.443 | -0.193 |
| AD | -0.837 | -0.442 | -0.112 | 0.191 | 0.029 | -0.2 |
| MCI | -0.35 | -0.895 | -0.508 | -0.63 | -1.162 | -0.201 |
| MCI | -0.03 | 0.077 | -0.209 | -0.26 | 0.004 | -0.203 |
| CN | -0.351 | -0.245 | -0.424 | -0.168 | -0.075 | -0.203 |
| CN | -0.218 | 0.103 | -0.288 | 0.249 | 0.361 | -0.21 |
| MCI | -0.751 | -0.74 | -0.245 | -0.455 | -0.857 | -0.211 |
| MCI | -0.538 | 0.672 | -0.698 | -0.241 | 0.257 | -0.213 |
| MCI | -0.182 | -0.704 | -0.387 | -0.32 | -0.329 | -0.216 |
| AD | 0.185 | -0.327 | -0.766 | -0.307 | -0.018 | -0.219 |
| CN | 0.591 | 0.223 | 0.368 | -0.158 | -0.227 | -0.229 |
| AD | 0.142 | 0.055 | 0.167 | -0.245 | -0.204 | -0.235 |
| AD | -0.472 | -0.819 | -0.182 | -0.176 | -0.403 | -0.239 |
| AD | -1.109 | -0.58 | -0.449 | -1.203 | -1.088 | -0.241 |
| AD | -0.206 | -0.778 | -0.44 | -0.458 | -0.619 | -0.244 |
| MCI | -0.04 | -0.315 | -0.058 | -0.02 | -0.157 | -0.25 |
| MCI | -0.906 | -0.737 | -0.272 | -0.055 | -0.157 | -0.252 |
| CN | -0.155 | 0.059 | 0.385 | 0.116 | 0.232 | -0.265 |
| MCI | -0.786 | -0.581 | -0.413 | -0.26 | -0.523 | -0.266 |
| MCI | 0.299 | 0.461 | 0.506 | 0.672 | 0.529 | -0.272 |
| CN | 0.048 | -0.221 | -0.3 | -0.58 | -0.377 | -0.273 |
| MCI | 0.449 | -0.514 | -0.614 | -0.344 | -0.2 | -0.278 |
| MCI | -0.341 | -0.765 | -1.129 | -0.717 | -0.642 | -0.287 |
| AD | -0.267 | -0.249 | 0.174 | -0.274 | -0.326 | -0.291 |
| MCI | -0.213 | -0.439 | -0.295 | -0.193 | -0.118 | -0.3 |
| CN | 0.111 | -0.498 | -0.606 | -0.615 | -0.888 | -0.3 |
| MCI | -0.417 | -0.621 | 0.146 | 0.064 | -0.216 | -0.307 |
| MCI | 0.054 | -0.611 | -0.414 | -0.504 | -0.698 | -0.314 |
| AD | -0.874 | -0.335 | -0.367 | -0.276 | -0.147 | -0.317 |
| MCI | 0.038 | 0.339 | -0.605 | 0.045 | 0.584 | -0.32 |
| MCI | -0.448 | 0.236 | -0.385 | -0.128 | 0.002 | -0.321 |
| AD | 0.025 | -0.456 | -0.025 | 0.447 | 0.326 | -0.322 |
| AD | 0.295 | -0.386 | -0.103 | 0.139 | 0.168 | -0.323 |
| MCI | -0.43 | -0.424 | -0.499 | -0.089 | -0.131 | -0.327 |
| MCI | 0.15 | -0.463 | -0.199 | -0.433 | -0.471 | -0.328 |
| CN | -0.474 | -0.145 | -0.677 | -0.038 | 0.165 | -0.331 |
| MCI | -0.452 | -0.295 | -0.353 | -0.491 | -0.075 | -0.331 |
| MCI | 0.268 | 0.124 | -0.282 | 0.183 | 0.286 | -0.338 |
| MCI | 0.093 | -0.476 | -0.424 | -0.828 | -0.437 | -0.354 |
| AD | -0.909 | -0.883 | -0.16 | -0.817 | -0.743 | -0.355 |
| AD | -1.177 | -1.036 | -0.351 | -0.954 | -1.37 | -0.355 |
| MCI | -1.218 | -0.592 | -0.767 | -0.365 | -0.164 | -0.362 |
| CN | -0.248 | -0.695 | 0.374 | -0.522 | -1.18 | -0.365 |
| CN | 0.13 | -0.849 | -0.107 | -0.266 | -0.541 | -0.369 |
| MCI | -0.772 | -0.561 | -0.636 | -0.004 | 0.476 | -0.37 |
| MCI | -0.934 | -0.512 | -0.59 | -0.724 | -0.416 | -0.373 |
| CN | -0.679 | 0.03 | -0.093 | -0.135 | 0.042 | -0.375 |
| AD | -0.848 | -1.033 | -0.225 | -0.825 | -1.074 | -0.379 |
| MCI | -1.166 | -0.219 | -0.683 | 0.112 | 0.467 | -0.387 |
| AD | -0.11 | -0.607 | -0.321 | -1.085 | -1.676 | -0.387 |
| MCI | 0.006 | -0.255 | -0.752 | -0.246 | 0.248 | -0.388 |
| MCI | -0.261 | 0.214 | -0.118 | -0.202 | 0.112 | -0.394 |
| MCI | -0.579 | -1.528 | -0.277 | -0.341 | -0.608 | -0.396 |
| CN | 0.11 | -0.504 | -0.702 | -0.586 | -0.317 | -0.404 |
| AD | -0.588 | -0.41 | -0.629 | -0.905 | -0.422 | -0.408 |
| MCI | 0.09 | -0.263 | -0.223 | -0.208 | -0.305 | -0.41 |
| CN | -0.037 | 0.025 | -0.097 | -0.019 | -0.455 | -0.411 |
| MCI | 0.003 | -0.712 | -0.672 | -0.387 | -0.225 | -0.413 |
| CN | -0.749 | -0.594 | -0.717 | -0.337 | -0.144 | -0.416 |
| MCI | -0.532 | -0.272 | -0.608 | -0.542 | 0.071 | -0.418 |
| AD | -0.742 | 1.03 | -0.977 | 0.184 | 0.703 | -0.42 |
| MCI | -0.755 | -0.72 | -0.508 | -0.85 | -1.091 | -0.42 |
| AD | -0.413 | -0.51 | -0.46 | -0.159 | -0.18 | -0.422 |
| CN | -0.522 | -0.634 | -0.137 | -0.544 | -0.85 | -0.422 |
| AD | -0.265 | -0.035 | -0.52 | -0.501 | 0.239 | -0.423 |
| MCI | -0.761 | -0.329 | -0.953 | 0.167 | 0.428 | -0.424 |
| MCI | -0.727 | -0.107 | -0.756 | -0.807 | -0.176 | -0.426 |
| CN | -0.671 | 0.315 | -0.314 | -0.052 | 0.015 | -0.427 |
| MCI | -0.651 | -0.457 | 0.088 | -0.159 | -0.783 | -0.43 |
| MCI | -0.649 | 0.093 | -0.009 | -0.119 | 0.243 | -0.435 |
| CN | -0.586 | 0 | -0.779 | -0.277 | 0.194 | -0.437 |
| CN | -0.586 | -0.543 | -0.189 | -0.619 | -0.543 | -0.441 |
| AD | -0.829 | -0.293 | -0.681 | -0.466 | -0.191 | -0.444 |
| MCI | -1.211 | -0.826 | -0.792 | -0.081 | 0.283 | -0.445 |
| MCI | -0.848 | -0.934 | -0.259 | -0.712 | -0.92 | -0.446 |
| MCI | -0.366 | -0.212 | -0.317 | -0.497 | 0.043 | -0.447 |
| MCI | -0.333 | -0.777 | -0.415 | -0.315 | -0.04 | -0.454 |
| AD | -0.409 | -0.582 | -0.484 | -0.873 | -1.241 | -0.459 |
| MCI | -0.824 | -0.214 | -0.776 | -0.308 | -0.1 | -0.469 |
| CN | -0.042 | -1.101 | -0.257 | -0.42 | -0.876 | -0.475 |
| AD | -0.515 | -0.553 | -0.173 | -0.57 | -0.849 | -0.484 |
| CN | -0.108 | 0.31 | -0.927 | 0.614 | 1.627 | -0.486 |
| AD | -0.237 | -1.373 | 0.142 | -0.069 | -0.723 | -0.486 |
| MCI | -0.965 | -0.457 | -0.536 | 0.071 | 0.393 | -0.492 |
| MCI | -0.157 | 0.944 | -0.955 | -0.172 | 0.591 | -0.501 |
| AD | 0.398 | 0.496 | 0.431 | -0.038 | 0.041 | -0.501 |
| CN | -0.424 | -0.138 | -0.365 | -0.332 | 0.009 | -0.501 |
| CN | -0.849 | -0.466 | -0.668 | -0.331 | -0.188 | -0.507 |
| CN | -0.28 | -0.043 | -0.441 | -0.221 | -0.253 | -0.511 |
| MCI | -0.344 | -0.378 | -0.31 | -0.587 | -0.282 | -0.513 |
| CN | 0.155 | 0.129 | -0.263 | -0.458 | -0.503 | -0.515 |
| MCI | 0.18 | -0.222 | -0.539 | 0.225 | 0.449 | -0.517 |
| CN | -1.012 | -1.039 | -0.874 | -0.311 | -0.526 | -0.517 |
| MCI | -0.505 | -1.178 | -0.133 | -0.138 | -0.869 | -0.527 |
| CN | -0.738 | 0.693 | -1.162 | -0.083 | 0.786 | -0.528 |
| MCI | -0.305 | 0.051 | -0.495 | -1.001 | -0.744 | -0.534 |
| MCI | -0.65 | -0.321 | -0.657 | -0.524 | -0.385 | -0.539 |
| CN | -1.238 | -0.735 | -0.342 | -0.162 | -0.649 | -0.544 |
| MCI | -0.104 | -0.652 | 0.021 | -0.645 | -1.147 | -0.545 |
| AD | -0.638 | -0.547 | -0.54 | -0.678 | -0.569 | -0.555 |
| MCI | -1.025 | -0.614 | -0.304 | -0.206 | -0.338 | -0.563 |
| CN | -0.166 | -0.982 | 0.005 | -0.031 | -0.549 | -0.567 |
| MCI | -0.366 | -0.825 | -0.329 | -0.98 | -0.949 | -0.572 |
| MCI | -0.544 | 1.179 | -1.112 | 0.774 | 1.711 | -0.573 |
| CN | -0.693 | -0.709 | 0.921 | 0.587 | -0.727 | -0.576 |
| MCI | 0.529 | -0.541 | -0.405 | -0.38 | -0.597 | -0.579 |
| MCI | -0.655 | -0.984 | -1.079 | -0.741 | -0.828 | -0.581 |
| MCI | -0.779 | -1.281 | -0.403 | -0.731 | -0.99 | -0.59 |
| AD | -0.931 | -0.131 | -0.874 | -0.02 | 0.386 | -0.604 |
| MCI | -0.492 | -0.303 | -0.298 | -0.816 | -0.568 | -0.613 |
| MCI | -0.782 | -0.567 | -0.536 | -0.887 | -0.762 | -0.613 |
| MCI | -1.105 | -0.844 | -0.516 | -1.027 | -1.318 | -0.614 |
| AD | -0.637 | -1.44 | -0.506 | -0.624 | -1.117 | -0.622 |
| MCI | -0.922 | -0.879 | -0.997 | -0.756 | -0.465 | -0.626 |
| MCI | -0.461 | -0.736 | -0.737 | -0.819 | -1.007 | -0.632 |
| CN | -0.259 | 0.32 | -0.288 | -0.585 | -0.347 | -0.635 |
| AD | 0.032 | -0.453 | -0.145 | -0.672 | -0.968 | -0.642 |
| MCI | -1.212 | -1.585 | -0.211 | -0.686 | -1.853 | -0.645 |
| AD | -0.573 | -0.302 | -0.722 | -0.488 | -0.115 | -0.662 |
| AD | -0.567 | -0.282 | -0.164 | -0.19 | -0.165 | -0.67 |
| MCI | -1.096 | -0.522 | -0.547 | -0.78 | -0.12 | -0.675 |
| AD | -1.063 | -0.386 | -0.639 | -0.671 | -0.273 | -0.679 |
| AD | 0.215 | -1.332 | -0.183 | -0.109 | -0.81 | -0.682 |
| AD | -0.126 | -0.459 | -0.389 | -0.704 | -0.528 | -0.684 |
| CN | -0.707 | -0.431 | -0.346 | -0.675 | -0.73 | -0.685 |
| AD | -0.957 | -0.18 | -1.051 | -0.955 | -0.443 | -0.687 |
| AD | -0.517 | -0.515 | -0.511 | -0.539 | -0.028 | -0.701 |
| MCI | -1.471 | -0.491 | -1.1 | -0.877 | -0.25 | -0.701 |
| CN | -0.227 | 0.802 | -1.47 | 0.031 | 1.344 | -0.71 |
| MCI | -0.368 | -0.163 | -0.593 | -0.921 | -0.805 | -0.714 |
| MCI | -0.874 | 0.395 | -0.468 | 0.106 | 0.191 | -0.717 |
| MCI | -0.267 | -0.425 | -0.96 | -0.765 | -0.434 | -0.717 |
| AD | -1.089 | 0.113 | -1.168 | 0.025 | 0.374 | -0.719 |
| CN | -0.531 | -0.023 | -0.982 | -1.169 | -0.364 | -0.72 |
| MCI | -0.315 | -1.029 | -0.951 | -0.971 | -1.092 | -0.724 |
| AD | -0.326 | -0.979 | -0.395 | -0.606 | -1.049 | -0.725 |
| MCI | -1.029 | -1.096 | -1.197 | -0.902 | -0.796 | -0.729 |
| CN | -0.75 | -1.498 | -0.408 | -0.56 | -1.087 | -0.739 |
| MCI | -0.589 | -1.345 | -0.425 | -0.447 | -0.895 | -0.74 |
| AD | -1.61 | -1.324 | -0.313 | -1.176 | -1.428 | -0.741 |
| MCI | -1.189 | -0.717 | -0.556 | -0.667 | -0.464 | -0.749 |
| CN | -0.553 | -0.44 | -0.606 | -1.039 | -0.889 | -0.752 |
| CN | -1.038 | -0.462 | -0.134 | -0.281 | -0.428 | -0.754 |
| MCI | -1.55 | -1.115 | -0.657 | -0.671 | -0.975 | -0.763 |
| AD | -0.651 | -0.579 | -1.12 | -1.042 | -0.572 | -0.764 |
| CN | 0.103 | -0.421 | -0.321 | -0.993 | -0.446 | -0.77 |
| CN | -1.123 | -0.873 | -0.563 | -0.803 | -0.916 | -0.772 |
| AD | -0.547 | 0.479 | -1.014 | -0.879 | 0.014 | -0.776 |
| CN | -0.298 | -0.435 | -0.419 | -0.64 | -0.547 | -0.776 |
| CN | -0.062 | -0.838 | -0.464 | -0.618 | -0.469 | -0.777 |
| CN | -0.515 | -1.447 | -0.712 | -0.717 | -0.609 | -0.785 |
| MCI | -0.823 | -1.445 | -0.704 | -0.805 | -1.139 | -0.788 |
| MCI | -0.587 | -0.32 | -0.881 | -0.593 | -0.439 | -0.793 |
| MCI | -1.317 | -1.042 | -1.036 | -0.826 | -0.61 | -0.795 |
| AD | -1.081 | -1.572 | -1.012 | -1.002 | -0.749 | -0.801 |
| MCI | -1.41 | -1.131 | -1.005 | -0.688 | -0.646 | -0.805 |
| CN | -0.905 | -0.56 | -0.776 | -0.442 | -0.043 | -0.821 |
| CN | -0.607 | -0.718 | -0.589 | -0.931 | -0.951 | -0.823 |
| MCI | -1.356 | -0.538 | -1.474 | -0.459 | 0.249 | -0.825 |
| CN | 0.018 | 0.601 | -0.589 | -0.695 | 0.028 | -0.832 |
| CN | -1.016 | -0.201 | -1.181 | -1.015 | -0.494 | -0.834 |
| AD | -1.108 | -0.56 | -0.729 | -0.611 | -0.536 | -0.851 |
| AD | -0.729 | -0.684 | -0.561 | -0.999 | -0.945 | -0.87 |
| MCI | -1.001 | -1.505 | -0.858 | -0.663 | -0.798 | -0.875 |
| MCI | -0.286 | 0.819 | -0.935 | -1.468 | -0.364 | -0.878 |
| AD | -1.026 | -1.034 | -0.816 | -1.039 | -1.102 | -0.883 |
| CN | -0.309 | -0.434 | -0.865 | -0.66 | -0.2 | -0.887 |
| MCI | -1.141 | -0.993 | -0.625 | -0.731 | -1.198 | -0.887 |
| MCI | -0.795 | -1.302 | -0.969 | -0.865 | -1.074 | -0.889 |
| MCI | -0.644 | -0.856 | -0.403 | -0.917 | -0.854 | -0.89 |
| MCI | -0.964 | -0.764 | -0.54 | -0.284 | -0.624 | -0.896 |
| AD | -1.249 | -0.836 | -0.601 | -1.066 | -1.227 | -0.896 |
| MCI | -1.166 | -0.531 | -1.069 | -0.872 | -0.272 | -0.909 |
| MCI | -0.659 | -1.378 | -0.052 | -0.65 | -1.374 | -0.921 |
| CN | -1.271 | -0.684 | -1.271 | -1.002 | -0.551 | -0.924 |
| MCI | -1.252 | -1.019 | -1.051 | -1.365 | -1.115 | -0.924 |
| AD | -1.385 | -1.289 | -0.973 | -1.323 | -1.094 | -0.931 |
| AD | -0.362 | -0.773 | -0.614 | -0.734 | -0.716 | -0.933 |
| MCI | -1.055 | -0.544 | -0.951 | -0.961 | -0.372 | -0.943 |
| CN | -0.82 | -0.766 | -0.724 | -0.743 | -0.694 | -0.949 |
| CN | -0.911 | -0.62 | -0.794 | -0.997 | -1.121 | -0.959 |
| MCI | -0.301 | -1.305 | -0.385 | -0.432 | -1.224 | -0.969 |
| CN | -0.982 | -0.636 | -1.093 | -1.099 | -0.737 | -0.972 |
| MCI | -0.736 | 0.229 | -0.77 | -0.648 | 0.135 | -0.975 |
| MCI | -0.52 | -0.898 | -1.187 | -1.12 | -0.406 | -0.976 |
| MCI | -0.953 | -0.964 | -0.901 | -1.229 | -1.086 | -0.996 |
| AD | -0.813 | -0.027 | -1.378 | -0.803 | -0.085 | -0.997 |
| AD | -1.277 | -1.416 | -0.993 | -0.941 | -0.831 | -1.004 |
| MCI | -0.989 | -0.649 | -1.01 | -1.252 | -0.818 | -1.016 |
| MCI | -0.712 | -0.243 | -0.688 | -1.046 | -0.54 | -1.029 |
| MCI | -1.001 | -0.593 | -0.751 | -1.251 | -0.525 | -1.032 |
| MCI | -1.163 | -1.531 | -0.817 | -0.895 | -1.528 | -1.032 |
| MCI | -1.396 | -1.033 | -0.948 | -0.982 | -1.22 | -1.037 |
| MCI | -1.082 | -0.899 | -1.336 | -1.045 | -0.247 | -1.038 |
| MCI | -0.287 | -1.416 | -0.683 | -1.136 | -0.708 | -1.039 |
| MCI | -0.796 | -1.011 | -1.463 | -1.152 | -0.776 | -1.044 |
| CN | -0.835 | -1.162 | -1.169 | -0.782 | -0.983 | -1.046 |
| MCI | -0.723 | -0.465 | -0.87 | -1.229 | -1.376 | -1.048 |
| AD | -1.816 | -0.682 | -1.668 | -0.86 | -0.29 | -1.051 |
| MCI | -0.914 | -0.481 | -1.171 | -1.165 | -0.762 | -1.072 |
| MCI | -1.642 | -0.996 | -1.037 | -0.78 | -0.636 | -1.078 |
| CN | -0.019 | 0.332 | -1.012 | -0.031 | 0.487 | -1.085 |
| MCI | -1.062 | -0.829 | -0.807 | -0.77 | -0.964 | -1.108 |
| AD | -0.336 | -0.644 | -1.261 | -1.111 | -0.721 | -1.109 |
| CN | -0.902 | -0.808 | -0.97 | -0.699 | -0.287 | -1.13 |
| MCI | -0.855 | -0.644 | -1.59 | -1.189 | -0.115 | -1.138 |
| AD | -0.91 | -0.16 | -1.205 | -0.931 | -0.51 | -1.141 |
| MCI | -1.204 | -1.292 | -1.051 | -1.1 | -1.45 | -1.143 |
| MCI | -0.488 | -0.97 | -0.485 | -0.336 | -0.883 | -1.145 |
| MCI | -0.369 | -1.176 | -0.68 | -1.21 | -1.071 | -1.145 |
| MCI | -0.845 | -0.31 | -1.14 | -0.954 | -0.553 | -1.165 |
| AD | -0.357 | -0.253 | -1.902 | -0.652 | 0.46 | -1.176 |
| AD | -0.938 | -0.305 | -1.332 | -1.09 | -0.58 | -1.177 |
| MCI | -0.515 | -1.057 | -1.427 | -0.96 | -0.217 | -1.182 |
| AD | -0.234 | -1.053 | -0.755 | -1.134 | -1.431 | -1.188 |
| AD | -0.326 | 0.189 | -1.002 | -1.114 | -0.548 | -1.195 |
| MCI | -0.297 | -1.003 | -0.685 | -0.94 | -0.653 | -1.203 |
| AD | -1.246 | -1.819 | -1.109 | -0.996 | -0.907 | -1.219 |
| MCI | -0.268 | -0.981 | -0.978 | -1.305 | -1.5 | -1.221 |
| AD | -1.486 | -0.87 | -0.82 | -0.268 | -0.209 | -1.236 |
| MCI | -1.299 | -0.804 | -1.496 | -0.999 | -0.599 | -1.247 |
| MCI | -1.144 | -1.006 | -1.103 | -1.476 | -1.734 | -1.25 |
| MCI | -0.647 | -0.903 | -1.037 | -1.163 | -0.697 | -1.256 |
| CN | -0.583 | -1.693 | -0.844 | -1.006 | -1.785 | -1.26 |
| MCI | -1.587 | -1.865 | -0.512 | -1.203 | -2.398 | -1.274 |
| AD | -0.807 | -0.61 | -1.148 | -1.102 | -0.66 | -1.299 |
| MCI | -0.461 | -1.374 | -0.777 | -1.538 | -2.075 | -1.3 |
| CN | -1.134 | -2.045 | -0.917 | -1.288 | -2.21 | -1.322 |
| AD | -0.927 | -1.471 | -0.866 | -1.139 | -1.138 | -1.333 |
| MCI | -1.153 | -1.123 | -1.081 | -1.744 | -1.747 | -1.338 |
| AD | -0.795 | -1.768 | -1.419 | -1.274 | -0.759 | -1.342 |
| MCI | -1.178 | -1.322 | -1.379 | -1.501 | -1.225 | -1.348 |
| MCI | -0.83 | -1.856 | -1.198 | -1.129 | -2.019 | -1.352 |
| MCI | -2.322 | -1.757 | -1.59 | -1.612 | -1.785 | -1.353 |
| MCI | -1.651 | -1.545 | -1.461 | -1.449 | -1.123 | -1.365 |
| MCI | -1.176 | -1.748 | -0.788 | -0.937 | -1.339 | -1.367 |
| MCI | -1.017 | 0.044 | -1.535 | 0.107 | -0.001 | -1.376 |
| MCI | -1.753 | -1.765 | -1.793 | -1.551 | -1.85 | -1.389 |
| CN | -1.518 | -1.755 | -1.156 | -1.713 | -2.553 | -1.406 |
| MCI | -0.598 | -0.993 | -1.372 | -0.808 | -0.116 | -1.418 |
| MCI | -1.993 | -1.414 | -1.518 | -1.631 | -1.83 | -1.441 |
| AD | -0.639 | -0.133 | -1.482 | -1.573 | -1.059 | -1.45 |
| AD | -1.843 | -1.025 | -1.906 | -1.61 | -1.359 | -1.456 |
| MCI | -1.841 | -2.553 | -1.42 | -1.829 | -2.535 | -1.477 |
| CN | -0.114 | 0.024 | -1.213 | -1.5 | -0.859 | -1.5 |
| CN | -1.14 | -1.213 | -1.102 | -1.564 | -1.578 | -1.508 |
| MCI | -0.93 | -0.898 | -0.939 | -1.155 | -0.973 | -1.518 |
| AD | -1.056 | -0.88 | -1.637 | -0.653 | 0.014 | -1.534 |
| CN | -1.694 | -0.96 | -1.51 | -1.123 | -1.168 | -1.536 |
| MCI | -0.689 | -0.724 | -1.342 | -1.368 | -0.711 | -1.543 |
| MCI | -1.304 | -1.543 | -1.135 | -1.056 | -1.027 | -1.557 |
| CN | -0.618 | -1.272 | -1.211 | -1.733 | -1.035 | -1.596 |
| MCI | -0.709 | -1.255 | -1.57 | -1.632 | -1.757 | -1.597 |
| AD | -0.664 | -0.949 | -1.419 | -1.669 | -1.197 | -1.626 |
| CN | -1.701 | -1.54 | -1.32 | -1.935 | -1.634 | -1.634 |
| AD | -1.642 | -1.363 | -1.506 | -1.872 | -2.533 | -1.657 |
| AD | -1.409 | -1.322 | -1.618 | -2.062 | -1.722 | -1.668 |
| MCI | -1.402 | -2.26 | -1.628 | -2.254 | -1.806 | -1.681 |
| CN | -2.55 | -1.938 | -1.895 | -1.443 | -1.907 | -1.697 |
| AD | -2.138 | -1.09 | -1.954 | -2.106 | -1.93 | -1.721 |
| MCI | -0.259 | -0.346 | -2.177 | -1.682 | -0.592 | -1.726 |
| MCI | -1.653 | -1.294 | -1.518 | -1.919 | -1.704 | -1.742 |
| MCI | -1.1 | -1.018 | -1.846 | -1.74 | -1.184 | -1.795 |
| MCI | -2.037 | -1.73 | -1.847 | -1.944 | -1.961 | -1.811 |
| MCI | -1.211 | -1.337 | -2.048 | -2.185 | -1.422 | -1.831 |
| MCI | -1.836 | -1.434 | -2.002 | -1.698 | -1.108 | -1.853 |
| AD | -1.776 | -1.411 | -1.555 | -2.206 | -2.075 | -1.877 |
| AD | -1.748 | -1.52 | -1.875 | -1.867 | -2.155 | -1.877 |
| MCI | -0.88 | -0.623 | -1.957 | -1.659 | -1.077 | -1.881 |
| AD | -1.763 | -2.106 | -1.669 | -2.327 | -2.363 | -1.889 |
| AD | -0.847 | -1.879 | -1.848 | -1.766 | -2.123 | -1.89 |
| MCI | -1.912 | -2.313 | -1.933 | -2.512 | -2.733 | -1.905 |
| AD | -1.303 | -1.146 | -1.44 | -2.11 | -1.534 | -1.911 |
| MCI | -1.463 | -1.905 | -1.728 | -2.261 | -2.029 | -1.949 |
| AD | -2.086 | -1.552 | -1.799 | -1.61 | -1.147 | -2.003 |
| MCI | -1.005 | -1.616 | -1.607 | -1.87 | -1.838 | -2.025 |
| MCI | -1.725 | -2.104 | -1.684 | -2.283 | -2.71 | -2.085 |
| AD | -1.636 | -1.514 | -2.021 | -2.174 | -2.534 | -2.119 |
| AD | -1.414 | -1.393 | -1.479 | -1.994 | -1.962 | -2.129 |
| MCI | -1.379 | -2.376 | -1.945 | -2.299 | -2.342 | -2.175 |
| MCI | -1.321 | -1.813 | -1.982 | -2.191 | -1.981 | -2.247 |
| MCI | -2.291 | -1.798 | -1.934 | -2.418 | -2.143 | -2.25 |
| MCI | -2.617 | -2.302 | -2.127 | -2.821 | -2.775 | -2.357 |
| CN | -2.161 | -2.196 | -2.042 | -2.635 | -2.962 | -2.401 |
| MCI | -2.472 | -1.92 | -2.237 | -0.94 | -1.074 | -2.419 |

Table e-4: Adjusted values for top contributors to “PC3”
TG values were adjusted for the following covariates: Age, Sex, APOE status, body mass index (BMI), and total triglycerides. Units are normalized peak height.

DXGrp= Diagnostic group; CN= control; MCI= mild cognitive impairment; AD= Alzheimer’s disease

| Table e-4: Adjusted values for top contributors to “PC3” | | | | | | | |
| --- | --- | --- | --- | --- | --- | --- | --- |
| DXGrp | TG 58:1 | TG 58:2 | TG 58:3 | TG 60:2 | TG 60:3 | TG 60:4 | TG 62:3 |
| MCI | 2.635 | 3.200 | 3.170 | 3.156 | 3.123 | 3.105 | 3.097 |
| CN | 3.025 | 3.043 | 3.093 | 3.061 | 3.085 | 3.099 | 3.106 |
| AD | 3.071 | 3.058 | 3.123 | 3.077 | 3.070 | 3.038 | 3.054 |
| CN | 2.271 | 2.778 | 3.021 | 2.812 | 3.004 | 3.011 | 2.697 |
| AD | 0.684 | 0.838 | 1.166 | 0.907 | 1.396 | 2.937 | 1.806 |
| CN | 2.432 | 2.487 | 2.608 | 2.387 | 2.545 | 2.801 | 3.061 |
| MCI | 1.178 | 1.661 | 2.621 | 1.404 | 2.261 | 2.767 | 1.894 |
| CN | 2.665 | 2.659 | 2.675 | 2.707 | 2.685 | 2.668 | 2.711 |
| MCI | 2.131 | 2.371 | 2.492 | 2.341 | 2.660 | 2.580 | 2.268 |
| MCI | 2.288 | 2.310 | 2.412 | 2.483 | 2.515 | 2.482 | 2.497 |
| CN | 2.625 | 2.772 | 2.825 | 2.811 | 2.845 | 2.472 | 2.882 |
| CN | 2.306 | 3.137 | 2.932 | 3.287 | 2.983 | 2.295 | 2.884 |
| MCI | 2.295 | 2.514 | 2.694 | 2.306 | 2.331 | 2.270 | 1.972 |
| MCI | 2.421 | 2.421 | 2.545 | 2.540 | 2.542 | 2.256 | 2.576 |
| MCI | 2.114 | 2.113 | 2.146 | 2.275 | 2.263 | 2.251 | 2.334 |
| MCI | 2.253 | 3.022 | 1.756 | 3.084 | 2.296 | 2.141 | 2.460 |
| CN | 2.341 | 2.531 | 2.514 | 2.770 | 2.671 | 2.121 | 2.565 |
| AD | 1.086 | 1.490 | 1.951 | 1.400 | 1.719 | 2.114 | 1.565 |
| AD | 0.328 | 0.650 | 1.204 | 0.586 | 1.251 | 2.045 | 1.355 |
| MCI | 1.313 | 1.516 | 1.727 | 1.536 | 1.840 | 1.977 | 1.722 |
| CN | 2.022 | 1.732 | 1.821 | 1.827 | 1.762 | 1.824 | 2.135 |
| MCI | -0.197 | 0.132 | 0.641 | 0.362 | 1.008 | 1.709 | 0.798 |
| MCI | 2.004 | 1.851 | 1.659 | 1.868 | 1.763 | 1.708 | 1.756 |
| MCI | 0.883 | 1.102 | 1.250 | 1.113 | 1.408 | 1.707 | 1.199 |
| CN | 2.036 | 2.012 | 2.047 | 1.730 | 1.676 | 1.694 | 1.868 |
| MCI | 0.614 | 0.813 | 1.186 | 0.679 | 1.093 | 1.692 | 0.824 |
| CN | 2.411 | 2.023 | 1.631 | 1.877 | 1.507 | 1.627 | 1.626 |
| AD | 1.432 | 2.209 | 2.627 | 1.510 | 1.681 | 1.607 | 1.871 |
| CN | 1.578 | 1.526 | 1.232 | 1.259 | 1.252 | 1.580 | 1.483 |
| CN | 1.418 | 1.496 | 1.723 | 1.467 | 1.570 | 1.533 | 1.241 |
| CN | -0.528 | 0.021 | 0.506 | -0.452 | 0.327 | 1.513 | 0.005 |
| MCI | 0.321 | 0.559 | 0.980 | 0.667 | 1.115 | 1.512 | 1.009 |
| AD | 0.689 | 0.982 | 1.309 | 1.002 | 1.318 | 1.510 | 1.335 |
| MCI | 0.135 | 0.663 | 0.919 | 0.807 | 1.247 | 1.504 | 1.146 |
| AD | 1.137 | 1.412 | 1.404 | 1.639 | 1.835 | 1.494 | 1.720 |
| MCI | 1.905 | 1.974 | 1.939 | 2.384 | 2.052 | 1.492 | 2.222 |
| MCI | 1.680 | 1.646 | 1.632 | 1.314 | 1.545 | 1.393 | 1.140 |
| AD | 0.755 | 0.831 | 0.746 | 1.127 | 1.295 | 1.381 | 1.417 |
| AD | 1.276 | 1.546 | 1.569 | 1.774 | 1.738 | 1.366 | 1.709 |
| CN | 0.378 | 1.316 | 2.138 | 0.808 | 1.679 | 1.346 | 2.306 |
| MCI | 1.395 | 1.484 | 1.146 | 1.439 | 1.362 | 1.339 | 1.373 |
| MCI | 1.347 | 1.257 | 1.390 | 0.966 | 0.936 | 1.323 | 0.588 |
| CN | 0.353 | 0.672 | 1.009 | 0.739 | 1.079 | 1.304 | 0.733 |
| CN | 1.303 | 1.018 | 0.922 | 1.051 | 1.114 | 1.268 | 1.243 |
| MCI | 1.056 | 1.136 | 1.151 | 1.095 | 1.129 | 1.264 | 1.262 |
| AD | 0.865 | 1.210 | 1.390 | 1.384 | 1.437 | 1.236 | 1.461 |
| MCI | 1.275 | 1.060 | 0.967 | 1.136 | 1.048 | 1.235 | 1.682 |
| CN | 1.061 | 0.847 | 0.844 | 0.868 | 0.929 | 1.225 | 1.068 |
| MCI | 0.796 | 0.818 | 0.746 | 0.691 | 0.853 | 1.193 | 0.593 |
| AD | 0.555 | 0.948 | 0.925 | 1.077 | 1.134 | 1.177 | 1.508 |
| MCI | 0.817 | 0.924 | 1.354 | 0.999 | 1.222 | 1.160 | 1.662 |
| MCI | -0.070 | 0.071 | 0.027 | 0.117 | 0.577 | 1.146 | 0.790 |
| MCI | 1.588 | 1.305 | 1.346 | 1.142 | 0.944 | 1.145 | 0.894 |
| MCI | 1.232 | 1.565 | 1.322 | 1.634 | 1.461 | 1.116 | 1.577 |
| CN | 0.599 | 0.773 | 0.836 | 0.516 | 0.874 | 1.115 | 0.792 |
| MCI | 2.483 | 1.994 | 1.490 | 2.152 | 1.771 | 1.113 | 1.642 |
| AD | 1.246 | 0.989 | 0.920 | 0.968 | 0.996 | 1.103 | 1.526 |
| MCI | -0.433 | -0.072 | 0.433 | -0.370 | 0.328 | 1.095 | -0.011 |
| AD | 0.471 | 0.884 | 0.819 | 1.110 | 1.250 | 1.092 | 0.947 |
| CN | 1.084 | 0.973 | 0.738 | 0.983 | 1.101 | 1.066 | 0.827 |
| CN | 0.554 | 0.971 | 1.054 | 1.119 | 1.184 | 1.053 | 1.269 |
| MCI | 1.506 | 1.586 | 1.437 | 1.727 | 1.522 | 1.052 | 1.301 |
| MCI | 0.279 | 0.445 | 0.174 | 0.238 | 0.531 | 1.021 | 0.647 |
| MCI | 0.589 | 0.710 | 0.799 | 0.671 | 0.874 | 1.019 | 0.543 |
| CN | 1.303 | 1.019 | 0.974 | 1.002 | 0.983 | 1.018 | 1.085 |
| MCI | 0.622 | 0.761 | 0.587 | 0.594 | 0.786 | 1.016 | 0.723 |
| MCI | 1.165 | 1.624 | 1.251 | 1.974 | 1.422 | 0.999 | 1.309 |
| MCI | 2.468 | 2.759 | 1.734 | 2.972 | 2.311 | 0.997 | 2.519 |
| CN | 1.318 | 1.442 | 1.156 | 1.345 | 1.309 | 0.987 | 1.239 |
| CN | -0.376 | -0.336 | 0.162 | -0.157 | 0.332 | 0.983 | 0.413 |
| MCI | 0.186 | 0.380 | 0.265 | 0.602 | 0.938 | 0.977 | 0.721 |
| AD | 1.481 | 1.299 | 1.465 | 1.520 | 1.150 | 0.974 | 1.408 |
| AD | 0.443 | 0.447 | 0.238 | 0.608 | 0.761 | 0.965 | 0.848 |
| MCI | 0.595 | 0.308 | 0.160 | 0.472 | 0.633 | 0.963 | 1.206 |
| AD | 1.043 | 1.391 | 1.214 | 1.541 | 1.326 | 0.945 | 1.108 |
| CN | 0.208 | 0.279 | 0.414 | 0.225 | 0.380 | 0.921 | 0.368 |
| CN | 0.604 | 0.809 | 0.971 | 0.746 | 0.908 | 0.918 | 0.646 |
| MCI | 0.012 | -0.078 | 0.004 | 0.004 | 0.182 | 0.914 | 0.451 |
| MCI | 1.305 | 1.032 | 0.594 | 0.660 | 0.672 | 0.914 | 0.595 |
| MCI | 1.026 | 0.988 | 0.973 | 0.762 | 0.681 | 0.908 | 0.947 |
| AD | -0.150 | 0.033 | -0.026 | -0.087 | 0.208 | 0.900 | 0.484 |
| MCI | 0.382 | 0.531 | 0.761 | 0.647 | 0.808 | 0.864 | 0.740 |
| MCI | 0.986 | 0.876 | 0.955 | 0.517 | 0.653 | 0.863 | 0.404 |
| CN | 0.021 | 0.549 | 0.776 | 0.249 | 0.418 | 0.850 | 0.340 |
| AD | -0.155 | 0.255 | 0.436 | 0.195 | 0.524 | 0.847 | 0.380 |
| AD | 0.737 | 0.797 | 0.671 | 0.532 | 0.686 | 0.832 | 0.393 |
| CN | 0.505 | 0.446 | 0.362 | 0.544 | 0.683 | 0.828 | 0.886 |
| CN | 0.362 | 0.270 | -0.132 | 0.156 | 0.492 | 0.816 | 0.369 |
| MCI | 1.240 | 1.030 | 0.624 | 1.264 | 0.947 | 0.808 | 0.900 |
| MCI | -0.006 | 0.053 | 0.429 | -0.198 | 0.073 | 0.802 | -0.627 |
| CN | 0.879 | 0.809 | 0.462 | 1.163 | 1.078 | 0.798 | 1.494 |
| MCI | -0.212 | 0.024 | 0.254 | 0.287 | 0.559 | 0.767 | 0.575 |
| CN | 0.610 | 0.364 | 0.092 | 0.445 | 0.508 | 0.750 | 0.996 |
| MCI | 0.542 | 0.469 | 0.324 | 0.422 | 0.453 | 0.749 | 0.482 |
| AD | -0.009 | 0.053 | 0.123 | 0.167 | 0.475 | 0.711 | 0.557 |
| AD | 1.510 | 1.115 | 0.846 | 0.985 | 0.742 | 0.704 | 0.684 |
| CN | 1.617 | 1.970 | 1.129 | 2.438 | 1.533 | 0.701 | 1.632 |
| MCI | 0.264 | 0.265 | 0.066 | 0.273 | 0.432 | 0.696 | 0.421 |
| MCI | 0.327 | 0.452 | 0.314 | 0.356 | 0.560 | 0.690 | 0.176 |
| MCI | 0.610 | 0.575 | 0.320 | 0.369 | 0.400 | 0.689 | 0.373 |
| AD | -0.274 | 0.054 | 0.240 | -0.311 | 0.261 | 0.687 | -0.754 |
| CN | 1.355 | 1.191 | 0.892 | 1.209 | 0.871 | 0.686 | 0.763 |
| CN | 0.543 | 0.400 | 0.003 | 0.174 | 0.261 | 0.673 | 0.072 |
| MCI | 1.027 | 0.977 | 0.781 | 1.016 | 0.879 | 0.672 | 0.769 |
| CN | 0.272 | 0.043 | -0.556 | -0.082 | -0.032 | 0.672 | 0.765 |
| AD | 0.845 | 0.828 | 0.554 | 0.551 | 0.543 | 0.667 | 0.585 |
| CN | 0.310 | 0.398 | 0.236 | 0.276 | 0.444 | 0.663 | 0.145 |
| AD | 0.047 | 0.051 | 0.359 | 0.345 | 0.528 | 0.661 | 1.022 |
| CN | 0.437 | 0.149 | 0.154 | 0.337 | 0.433 | 0.654 | 0.676 |
| MCI | -0.098 | 0.249 | 0.844 | 0.175 | 0.487 | 0.646 | 1.460 |
| MCI | 0.031 | 0.260 | 0.185 | 0.035 | 0.311 | 0.643 | 0.061 |
| MCI | 0.313 | 0.239 | 0.099 | 0.392 | 0.604 | 0.642 | 0.444 |
| MCI | -0.183 | -0.048 | -0.177 | -0.221 | 0.179 | 0.639 | -0.329 |
| MCI | 0.161 | 0.310 | 0.369 | 0.159 | 0.468 | 0.638 | 1.258 |
| CN | 0.604 | 0.735 | 0.691 | 0.629 | 0.760 | 0.634 | 0.549 |
| CN | 0.790 | 0.720 | 0.577 | 0.714 | 0.714 | 0.631 | 0.730 |
| CN | 1.314 | 1.189 | 0.817 | 1.121 | 0.851 | 0.627 | 1.236 |
| CN | -0.019 | 0.126 | 0.155 | 0.046 | 0.357 | 0.625 | 0.171 |
| AD | 0.706 | 0.608 | 0.291 | 0.495 | 0.515 | 0.621 | 0.194 |
| MCI | 0.026 | 0.195 | 0.280 | 0.302 | 0.442 | 0.620 | 0.495 |
| MCI | -0.494 | -0.350 | 0.073 | -0.272 | 0.269 | 0.619 | 0.059 |
| CN | 0.210 | 0.467 | 0.460 | 0.310 | 0.411 | 0.611 | 0.021 |
| AD | 0.934 | 0.972 | 0.883 | 0.851 | 0.611 | 0.602 | 0.620 |
| MCI | 2.008 | 1.819 | 1.416 | 1.718 | 1.507 | 0.601 | 1.065 |
| MCI | 1.459 | 0.760 | 0.350 | 0.972 | 0.508 | 0.600 | 1.552 |
| MCI | 0.285 | 0.127 | 0.132 | 0.258 | 0.232 | 0.589 | 1.052 |
| MCI | -0.175 | 0.010 | -0.149 | -0.132 | 0.250 | 0.587 | 0.261 |
| MCI | 0.826 | 0.438 | 0.311 | 0.416 | 0.358 | 0.587 | 0.673 |
| MCI | -0.370 | 0.220 | 0.199 | -0.130 | 0.267 | 0.583 | -0.249 |
| CN | 0.839 | 0.766 | 0.448 | 0.633 | 0.561 | 0.578 | 0.537 |
| MCI | 0.979 | 0.707 | 0.708 | 0.893 | 0.691 | 0.578 | 0.780 |
| MCI | 0.337 | 0.394 | 0.542 | 0.560 | 0.603 | 0.573 | 0.684 |
| CN | 0.419 | 0.494 | 0.366 | 0.519 | 0.565 | 0.569 | 0.402 |
| CN | 1.410 | 1.090 | 0.493 | 1.361 | 0.799 | 0.566 | 1.225 |
| MCI | 0.993 | 0.977 | 0.371 | 0.826 | 0.610 | 0.562 | 0.590 |
| AD | 0.351 | 0.359 | 0.133 | 0.217 | 0.384 | 0.559 | 0.157 |
| MCI | -0.254 | 0.027 | 0.236 | -0.054 | 0.231 | 0.559 | 0.361 |
| MCI | 0.066 | 0.132 | -0.193 | 0.019 | 0.321 | 0.545 | 0.126 |
| CN | 0.575 | 0.531 | 0.163 | 0.532 | 0.624 | 0.532 | 0.723 |
| MCI | 0.183 | -0.025 | 0.301 | -0.144 | 0.158 | 0.530 | 0.044 |
| AD | 0.479 | 0.530 | 0.285 | 0.404 | 0.451 | 0.528 | 0.115 |
| MCI | 0.077 | 0.194 | 0.199 | 0.086 | 0.321 | 0.527 | 0.342 |
| CN | 0.078 | 0.185 | -0.216 | 0.092 | 0.325 | 0.526 | 0.273 |
| AD | -0.201 | -0.047 | -0.194 | -0.012 | 0.324 | 0.526 | 0.273 |
| MCI | 0.707 | 0.738 | 0.834 | 0.482 | 0.497 | 0.503 | 0.206 |
| MCI | 0.407 | 0.465 | 0.483 | 0.603 | 0.581 | 0.502 | 0.353 |
| CN | 1.086 | 0.803 | 0.645 | 0.836 | 0.591 | 0.501 | 0.461 |
| MCI | 0.950 | 0.945 | 0.874 | 0.897 | 0.720 | 0.492 | 0.420 |
| MCI | 0.606 | 0.595 | 0.688 | 0.551 | 0.450 | 0.490 | 1.033 |
| MCI | -0.585 | 0.155 | 0.559 | -0.173 | 0.377 | 0.489 | 0.585 |
| CN | 0.500 | 0.623 | 0.145 | 0.191 | 0.432 | 0.485 | -0.084 |
| AD | -0.935 | -0.512 | 0.138 | -0.367 | -0.055 | 0.484 | 0.075 |
| MCI | -0.599 | -0.286 | -0.436 | -0.175 | 0.108 | 0.477 | 0.459 |
| CN | 1.096 | 0.836 | 0.514 | 0.985 | 0.693 | 0.476 | 1.322 |
| CN | 0.154 | 0.100 | -0.077 | 0.106 | 0.293 | 0.467 | 0.679 |
| MCI | 0.523 | 0.422 | 0.545 | 0.637 | 0.537 | 0.464 | 0.466 |
| MCI | 0.043 | 0.209 | 0.469 | -0.206 | 0.085 | 0.460 | -0.317 |
| CN | -0.044 | 0.328 | -0.080 | 0.388 | 0.551 | 0.458 | 0.259 |
| MCI | 0.234 | 0.214 | -0.069 | 0.288 | 0.258 | 0.457 | 0.220 |
| MCI | -0.641 | -0.353 | 0.115 | -0.598 | 0.047 | 0.455 | -0.751 |
| MCI | -0.162 | 0.015 | 0.416 | 0.077 | 0.373 | 0.453 | 0.265 |
| MCI | 0.638 | 0.546 | 0.381 | 0.428 | 0.545 | 0.452 | 0.143 |
| AD | 0.053 | 0.133 | -0.071 | -0.066 | 0.179 | 0.452 | 0.244 |
| AD | -0.008 | 0.160 | -0.079 | -0.033 | 0.141 | 0.448 | 0.052 |
| MCI | 0.136 | 0.224 | -0.012 | 0.111 | 0.277 | 0.446 | 0.122 |
| AD | 0.834 | 0.792 | 0.752 | 0.665 | 0.435 | 0.445 | 0.733 |
| MCI | 0.676 | 0.653 | 0.503 | 1.029 | 0.772 | 0.435 | 0.900 |
| MCI | -0.052 | -0.143 | 0.056 | 0.290 | 0.450 | 0.433 | 0.603 |
| MCI | -0.310 | 0.052 | 0.335 | 0.143 | 0.351 | 0.433 | 0.256 |
| MCI | -0.381 | -0.117 | 0.101 | -0.073 | 0.210 | 0.431 | 0.241 |
| AD | 1.555 | 0.861 | 0.149 | 0.838 | 0.095 | 0.429 | 1.202 |
| MCI | 0.187 | 0.448 | 0.551 | 0.490 | 0.571 | 0.429 | 0.526 |
| CN | -0.006 | 0.233 | 0.604 | 0.531 | 0.589 | 0.423 | 0.670 |
| AD | -0.086 | 0.198 | 0.034 | 0.136 | 0.402 | 0.423 | 0.129 |
| AD | 0.660 | 0.638 | 0.115 | 0.814 | 0.644 | 0.421 | 0.647 |
| MCI | 0.659 | 0.635 | 0.657 | 0.292 | 0.486 | 0.419 | 0.120 |
| AD | -0.057 | -0.008 | -0.050 | -0.118 | 0.155 | 0.417 | -0.079 |
| AD | -0.194 | 0.027 | -0.143 | 0.015 | 0.175 | 0.414 | -0.306 |
| MCI | -0.422 | -0.219 | -0.337 | -0.180 | 0.172 | 0.406 | 0.161 |
| MCI | 0.237 | 0.464 | 0.495 | 0.495 | 0.235 | 0.404 | 0.626 |
| AD | 0.016 | -0.128 | 0.090 | 0.144 | 0.133 | 0.400 | 0.326 |
| MCI | -0.210 | 0.169 | 0.023 | -0.130 | -0.096 | 0.398 | 0.122 |
| CN | 0.253 | 0.262 | 0.385 | 0.076 | 0.173 | 0.396 | 0.739 |
| CN | 0.674 | 0.668 | 0.386 | 0.442 | 0.458 | 0.394 | 0.303 |
| MCI | -0.157 | 0.043 | -0.158 | -0.032 | 0.247 | 0.383 | 0.214 |
| MCI | 1.155 | 0.857 | 0.465 | 0.669 | 0.399 | 0.375 | 0.463 |
| MCI | 0.610 | 1.702 | 1.170 | 1.767 | 1.241 | 0.371 | 0.865 |
| MCI | 0.528 | 0.442 | 0.366 | 0.146 | 0.368 | 0.368 | -0.188 |
| MCI | 2.437 | 1.737 | 1.050 | 1.569 | 0.934 | 0.366 | 1.079 |
| CN | -0.100 | -0.010 | -0.025 | -0.113 | 0.227 | 0.359 | -0.415 |
| MCI | 0.725 | 0.560 | 0.027 | 0.250 | -0.054 | 0.359 | 0.188 |
| AD | 0.285 | 0.304 | 0.141 | 0.280 | 0.390 | 0.359 | 0.265 |
| CN | 0.050 | 0.350 | -0.067 | 0.117 | 0.334 | 0.358 | -0.124 |
| MCI | 0.135 | 0.259 | 0.450 | 0.275 | 0.471 | 0.356 | 0.662 |
| MCI | -0.690 | -0.532 | -0.063 | -0.361 | 0.133 | 0.355 | -0.377 |
| MCI | -0.659 | -0.488 | -0.065 | -0.461 | 0.055 | 0.350 | -0.383 |
| MCI | 0.736 | 0.644 | 0.179 | 0.548 | 0.414 | 0.347 | 0.477 |
| MCI | -0.100 | 0.259 | -0.123 | -0.183 | 0.137 | 0.339 | -0.255 |
| CN | -0.194 | 0.308 | -0.173 | 0.099 | 0.110 | 0.335 | -0.113 |
| MCI | 0.489 | 0.212 | -0.039 | 0.128 | 0.161 | 0.324 | 0.579 |
| MCI | -1.260 | -0.793 | -0.634 | -0.526 | -0.018 | 0.323 | 0.355 |
| CN | 0.397 | 0.488 | 0.329 | 0.615 | 0.481 | 0.323 | 0.627 |
| CN | 0.555 | 0.267 | 0.063 | 0.396 | 0.334 | 0.322 | 0.656 |
| MCI | -0.654 | -0.151 | 0.153 | -0.048 | 0.251 | 0.319 | 0.095 |
| CN | 0.319 | -0.156 | -0.018 | 0.311 | 0.273 | 0.317 | 1.050 |
| MCI | 0.177 | 0.414 | 0.624 | 0.123 | 0.342 | 0.317 | 0.041 |
| CN | 0.002 | 0.120 | -0.137 | 0.016 | 0.225 | 0.315 | -0.129 |
| MCI | -0.294 | 0.056 | -0.094 | 0.522 | 0.588 | 0.311 | 0.466 |
| CN | -0.195 | -0.009 | -0.307 | -0.240 | 0.068 | 0.308 | -0.293 |
| AD | -0.102 | 0.038 | 0.302 | 0.001 | 0.188 | 0.304 | 0.070 |
| CN | 0.801 | 0.739 | 0.419 | 0.564 | 0.404 | 0.302 | 0.152 |
| AD | 0.636 | 0.304 | -0.297 | 0.125 | 0.040 | 0.293 | 0.472 |
| AD | 0.698 | 0.637 | 0.536 | 0.829 | 0.692 | 0.291 | 0.940 |
| MCI | 0.320 | 0.219 | -0.069 | 0.031 | 0.067 | 0.287 | 0.175 |
| AD | -0.924 | -0.599 | -0.021 | -0.346 | 0.075 | 0.287 | 0.401 |
| CN | -2.503 | -1.793 | -1.030 | -1.289 | -0.538 | 0.287 | 0.514 |
| CN | 0.025 | -0.052 | 0.333 | 0.270 | 0.288 | 0.280 | 0.539 |
| AD | 0.262 | 0.048 | -0.206 | 0.327 | 0.151 | 0.278 | 0.321 |
| AD | -0.854 | -0.597 | -0.200 | -0.534 | -0.063 | 0.276 | 0.102 |
| CN | 1.368 | 0.892 | 0.474 | 0.611 | 0.242 | 0.270 | 0.396 |
| CN | 0.782 | 1.254 | 0.839 | 1.184 | 0.682 | 0.268 | 0.721 |
| MCI | -0.381 | -0.175 | 0.082 | -0.273 | 0.086 | 0.267 | -0.294 |
| MCI | 0.263 | 0.183 | 0.118 | 0.211 | 0.054 | 0.266 | 0.200 |
| CN | 0.938 | 1.008 | 0.807 | 1.054 | 0.652 | 0.265 | 0.587 |
| MCI | -0.258 | -0.174 | 0.010 | -0.425 | -0.300 | 0.257 | -0.241 |
| MCI | -0.591 | -0.264 | -0.141 | -0.079 | 0.250 | 0.254 | 0.262 |
| AD | 0.389 | 0.570 | 0.294 | 0.338 | 0.410 | 0.251 | 0.117 |
| CN | 0.763 | 0.600 | 0.363 | 0.482 | 0.336 | 0.250 | 0.076 |
| MCI | -0.458 | -0.095 | -0.308 | -0.059 | -0.034 | 0.250 | 0.291 |
| AD | 0.437 | 0.259 | 0.116 | 0.010 | -0.029 | 0.249 | -0.182 |
| MCI | -0.275 | -0.100 | 0.206 | -0.351 | 0.080 | 0.247 | -0.540 |
| AD | 1.154 | 0.812 | 0.485 | 0.628 | 0.399 | 0.245 | 0.166 |
| CN | -1.616 | -1.232 | -0.442 | -0.644 | -0.251 | 0.238 | -0.208 |
| CN | 1.361 | 1.021 | 0.661 | 0.612 | 0.295 | 0.233 | 0.439 |
| CN | 0.754 | 0.552 | 0.483 | 0.315 | 0.179 | 0.228 | 0.007 |
| MCI | 0.337 | 0.506 | 0.356 | 0.279 | 0.215 | 0.228 | 0.742 |
| MCI | 0.038 | -0.111 | 0.274 | 0.118 | 0.243 | 0.227 | -0.292 |
| MCI | -0.141 | -0.150 | 0.225 | -0.232 | 0.006 | 0.225 | -0.631 |
| CN | -0.482 | -0.296 | 0.162 | -0.381 | -0.006 | 0.215 | -0.176 |
| CN | -0.008 | 0.601 | 0.287 | 0.289 | 0.452 | 0.213 | 0.122 |
| MCI | 0.577 | 0.477 | 0.113 | 0.317 | 0.325 | 0.212 | 0.179 |
| MCI | 0.612 | 0.347 | -0.374 | 0.064 | -0.294 | 0.211 | -0.144 |
| CN | 0.768 | 0.313 | -0.025 | 0.266 | 0.214 | 0.210 | -0.149 |
| AD | 0.452 | 0.442 | 0.428 | 0.055 | 0.168 | 0.210 | -0.263 |
| CN | 1.093 | 0.886 | 0.461 | 1.047 | 0.555 | 0.210 | 0.882 |
| AD | 0.215 | 0.363 | 0.356 | 0.348 | 0.294 | 0.209 | 0.364 |
| MCI | -0.076 | 0.073 | 0.361 | 0.028 | 0.017 | 0.207 | 0.376 |
| CN | 0.023 | 0.272 | 0.646 | 0.120 | 0.336 | 0.196 | 0.152 |
| CN | 0.400 | 0.443 | 0.256 | 0.412 | 0.220 | 0.194 | 0.452 |
| CN | 0.629 | 0.907 | 0.518 | 0.880 | 0.595 | 0.193 | 0.244 |
| AD | -0.313 | -0.426 | -0.501 | -0.318 | -0.062 | 0.187 | -0.467 |
| MCI | 0.088 | 0.102 | -0.244 | 0.082 | 0.105 | 0.185 | 0.107 |
| AD | 0.548 | 0.512 | 0.396 | 0.325 | 0.219 | 0.181 | 0.480 |
| AD | 0.542 | 0.359 | -0.368 | 0.269 | 0.301 | 0.175 | 0.171 |
| CN | 0.421 | 0.402 | -0.124 | 0.148 | 0.175 | 0.174 | 0.105 |
| AD | 0.054 | 0.045 | -0.105 | -0.092 | 0.066 | 0.171 | -0.310 |
| MCI | 0.109 | 0.163 | -0.289 | 0.072 | 0.208 | 0.170 | -0.054 |
| CN | 0.468 | 0.062 | -0.692 | -0.139 | -0.617 | 0.168 | -0.149 |
| MCI | 0.976 | 0.703 | 0.468 | 0.878 | 0.589 | 0.163 | 0.584 |
| CN | -0.575 | -0.480 | -0.064 | -0.159 | -0.221 | 0.162 | 0.146 |
| AD | -0.266 | -0.394 | -0.238 | -0.216 | -0.025 | 0.161 | 0.019 |
| MCI | -0.151 | -0.141 | -0.128 | -0.291 | -0.134 | 0.159 | -0.284 |
| MCI | -0.347 | -0.286 | -0.005 | -0.310 | -0.032 | 0.155 | -0.535 |
| AD | -0.565 | -0.280 | 0.016 | -0.305 | 0.045 | 0.152 | -0.834 |
| MCI | 0.225 | 0.284 | 0.109 | -0.033 | 0.074 | 0.145 | 0.336 |
| AD | 0.250 | 0.198 | -0.015 | -0.078 | 0.037 | 0.143 | 0.095 |
| MCI | 0.077 | 0.115 | 0.073 | -0.198 | -0.100 | 0.143 | -0.045 |
| MCI | -0.241 | -0.250 | -0.155 | -0.398 | -0.220 | 0.142 | -0.044 |
| CN | 0.535 | 0.359 | -0.030 | 0.269 | 0.176 | 0.138 | 0.057 |
| MCI | -1.015 | -0.589 | -0.137 | -0.495 | -0.028 | 0.136 | -0.235 |
| MCI | -0.417 | -0.153 | 0.135 | -0.574 | -0.278 | 0.134 | -0.622 |
| CN | 0.916 | 0.931 | 0.585 | 1.023 | 0.673 | 0.122 | 0.420 |
| MCI | 0.542 | 0.366 | 0.035 | 0.368 | 0.237 | 0.121 | 0.228 |
| CN | -0.019 | 0.104 | 0.227 | 0.043 | 0.182 | 0.119 | 0.181 |
| MCI | 1.199 | 0.822 | 0.532 | 1.569 | 1.178 | 0.119 | 1.401 |
| MCI | -0.271 | -0.200 | -0.199 | -0.091 | 0.018 | 0.117 | 0.030 |
| CN | 1.060 | 0.785 | 0.212 | 0.769 | 0.367 | 0.107 | 0.500 |
| AD | -0.243 | -0.312 | -0.471 | -0.562 | -0.484 | 0.105 | -0.203 |
| AD | -0.848 | -0.626 | -0.068 | -0.832 | -0.288 | 0.104 | -0.709 |
| MCI | -0.679 | -0.423 | -0.075 | -0.513 | -0.073 | 0.103 | -0.724 |
| AD | 0.284 | 0.247 | -0.055 | 0.650 | 0.376 | 0.102 | 0.333 |
| MCI | 0.409 | 0.099 | -0.475 | 0.126 | -0.041 | 0.099 | 0.281 |
| MCI | 0.761 | 0.250 | 0.033 | 0.488 | 0.254 | 0.095 | 0.591 |
| AD | 0.345 | 0.135 | 0.209 | -0.078 | 0.022 | 0.095 | -0.218 |
| AD | 1.268 | 0.779 | 0.719 | 1.108 | 0.697 | 0.093 | 0.964 |
| MCI | -0.880 | -0.442 | -0.796 | -0.451 | -0.461 | 0.093 | -0.013 |
| CN | 0.789 | 0.386 | 0.066 | 0.517 | 0.299 | 0.087 | 0.968 |
| MCI | 0.316 | 0.357 | 0.252 | 0.230 | 0.052 | 0.087 | 0.067 |
| CN | -0.163 | -0.021 | -0.398 | -0.316 | -0.038 | 0.085 | -0.809 |
| MCI | -0.198 | -0.139 | 0.005 | -0.289 | -0.025 | 0.083 | 0.395 |
| CN | -0.038 | 0.004 | -0.109 | -0.013 | 0.171 | 0.083 | -0.219 |
| MCI | -0.935 | -1.022 | -0.652 | -0.922 | -0.646 | 0.080 | -0.473 |
| CN | 0.389 | 0.370 | -0.022 | -0.026 | 0.054 | 0.080 | -0.216 |
| CN | 0.101 | 0.422 | 0.117 | 0.337 | 0.153 | 0.080 | 0.017 |
| MCI | -1.444 | -0.972 | -0.247 | -0.730 | -0.310 | 0.078 | -0.747 |
| CN | 1.129 | 0.729 | 0.298 | 0.396 | 0.192 | 0.076 | 0.359 |
| MCI | -0.378 | -0.486 | -0.122 | -0.430 | -0.174 | 0.075 | -0.532 |
| AD | -0.222 | -0.100 | 0.008 | 0.047 | 0.177 | 0.072 | 0.439 |
| MCI | -0.934 | -0.134 | 0.362 | -0.301 | 0.145 | 0.071 | 0.009 |
| MCI | -0.149 | -0.339 | -0.055 | -0.232 | -0.258 | 0.070 | -0.524 |
| CN | -0.295 | -0.092 | 0.206 | -0.187 | 0.100 | 0.067 | 0.151 |
| AD | 0.356 | 0.134 | -0.122 | 0.272 | -0.002 | 0.062 | 0.061 |
| MCI | -0.471 | -0.510 | -0.180 | -0.661 | -0.314 | 0.061 | -0.387 |
| CN | 1.246 | 0.770 | 0.505 | 0.771 | 0.356 | 0.059 | 0.816 |
| AD | -0.915 | -0.192 | 0.402 | -0.783 | -0.245 | 0.056 | -0.578 |
| CN | 0.006 | 0.171 | -0.004 | 0.156 | 0.076 | 0.049 | 0.096 |
| CN | 0.311 | 0.216 | 0.193 | 0.322 | 0.223 | 0.047 | 0.442 |
| CN | -0.017 | 0.556 | 0.426 | 0.326 | 0.291 | 0.047 | -0.077 |
| AD | 0.328 | 0.262 | 0.052 | 0.592 | 0.318 | 0.046 | 0.579 |
| AD | 0.100 | 0.205 | -0.097 | -0.045 | 0.078 | 0.039 | -0.269 |
| CN | 0.004 | 0.019 | 0.211 | -0.099 | -0.063 | 0.039 | -0.124 |
| AD | 0.024 | 0.013 | -0.712 | -0.116 | -0.077 | 0.039 | -0.343 |
| MCI | -0.266 | 0.017 | 0.180 | -0.093 | 0.055 | 0.037 | 0.089 |
| AD | -0.024 | 0.152 | 0.395 | -0.104 | 0.104 | 0.026 | -0.092 |
| AD | -0.386 | -0.334 | -0.472 | -0.504 | -0.202 | 0.022 | -0.969 |
| MCI | -0.111 | 0.005 | 0.148 | 0.171 | 0.243 | 0.012 | 0.098 |
| MCI | 0.086 | 0.081 | -0.091 | -0.041 | -0.035 | 0.008 | -0.104 |
| MCI | -0.240 | -0.223 | -0.665 | -0.190 | 0.020 | 0.006 | -0.003 |
| MCI | 0.073 | 0.114 | 0.184 | -0.124 | 0.102 | 0.005 | -0.511 |
| AD | -0.557 | -0.554 | -0.112 | -0.396 | -0.184 | 0.004 | -0.774 |
| MCI | -0.566 | -0.284 | -0.118 | -0.210 | -0.053 | 0.003 | -0.138 |
| MCI | -0.073 | -0.139 | -0.070 | -0.211 | -0.078 | -0.001 | 0.008 |
| MCI | -0.019 | 0.002 | 0.090 | 0.067 | 0.011 | -0.001 | 0.210 |
| MCI | 0.445 | 0.260 | -0.113 | 0.187 | 0.168 | -0.003 | 0.321 |
| CN | -0.018 | -0.166 | -0.457 | -0.375 | -0.155 | -0.004 | -0.557 |
| CN | 0.398 | 0.292 | 0.318 | 0.218 | 0.209 | -0.009 | 0.073 |
| MCI | 0.116 | -0.112 | 0.173 | 0.055 | 0.083 | -0.012 | -0.234 |
| MCI | -0.185 | -0.371 | -0.106 | -0.285 | 0.006 | -0.015 | -0.186 |
| MCI | -0.143 | -0.166 | 0.111 | -0.037 | 0.095 | -0.021 | -0.001 |
| CN | 0.182 | 0.220 | 0.506 | 0.179 | 0.204 | -0.021 | 0.130 |
| CN | -0.517 | -0.528 | -0.543 | -0.415 | -0.076 | -0.022 | -0.411 |
| CN | 0.860 | 0.258 | 0.374 | 0.387 | 0.064 | -0.029 | 0.871 |
| CN | -0.453 | -0.295 | -0.615 | -0.282 | -0.065 | -0.030 | -0.201 |
| AD | 0.248 | 0.158 | -0.225 | 0.160 | 0.147 | -0.031 | -0.190 |
| MCI | -0.153 | -0.046 | -0.321 | -0.042 | 0.009 | -0.033 | -0.051 |
| MCI | -0.419 | -0.565 | -1.004 | -0.421 | -0.571 | -0.038 | -0.025 |
| MCI | 1.129 | 0.699 | 0.373 | 0.805 | 0.324 | -0.039 | 0.708 |
| MCI | -0.301 | -0.302 | -0.659 | -0.402 | -0.218 | -0.040 | -0.159 |
| MCI | 0.439 | 0.700 | 0.385 | 0.665 | 0.417 | -0.040 | 0.236 |
| CN | 0.580 | 0.500 | 0.405 | 0.428 | 0.295 | -0.041 | 0.418 |
| MCI | -0.611 | -0.277 | 0.194 | -0.555 | -0.166 | -0.045 | -0.659 |
| MCI | 0.277 | 0.357 | 0.250 | 0.129 | -0.001 | -0.048 | 0.206 |
| AD | 0.097 | 0.168 | -0.429 | -0.203 | -0.041 | -0.049 | -0.304 |
| MCI | -0.934 | -0.915 | -0.691 | -0.840 | -0.589 | -0.052 | -0.560 |
| MCI | 0.868 | 0.381 | 0.100 | 0.463 | 0.160 | -0.053 | 0.056 |
| CN | -0.232 | -0.201 | -0.103 | -0.110 | 0.083 | -0.056 | -0.198 |
| MCI | -0.136 | 0.098 | 0.148 | 0.067 | 0.020 | -0.056 | -0.010 |
| MCI | -0.568 | -0.455 | -0.205 | -0.326 | -0.270 | -0.069 | -0.158 |
| AD | 0.368 | 0.344 | 0.169 | 0.151 | 0.040 | -0.072 | -0.140 |
| MCI | 0.090 | -0.093 | -0.116 | -0.148 | -0.085 | -0.073 | -0.016 |
| CN | -0.772 | -0.804 | -0.831 | -0.495 | -0.503 | -0.079 | -0.495 |
| CN | 0.365 | 0.154 | 0.245 | -0.029 | -0.050 | -0.079 | -0.452 |
| MCI | 0.149 | 0.059 | -0.001 | -0.098 | -0.337 | -0.080 | -0.092 |
| MCI | -0.012 | -0.032 | -0.014 | -0.273 | -0.096 | -0.082 | -0.499 |
| MCI | -0.475 | -0.329 | 0.031 | -0.384 | -0.111 | -0.082 | -1.480 |
| CN | -0.266 | -0.124 | -0.481 | -0.318 | -0.099 | -0.083 | -0.434 |
| MCI | -0.066 | -0.165 | -0.185 | -0.049 | -0.136 | -0.083 | 0.233 |
| CN | -0.919 | -0.568 | 0.014 | -0.326 | -0.008 | -0.083 | 0.055 |
| CN | 0.492 | 0.441 | 0.331 | 0.337 | 0.157 | -0.083 | 0.402 |
| AD | -0.666 | -0.251 | 0.028 | -0.198 | -0.123 | -0.084 | -0.403 |
| AD | -0.284 | -0.267 | -0.062 | -0.267 | -0.335 | -0.087 | -0.336 |
| CN | -0.129 | 0.041 | -0.398 | -0.086 | 0.087 | -0.091 | -0.230 |
| CN | 0.170 | 0.043 | -0.322 | 0.158 | 0.189 | -0.093 | 0.113 |
| CN | -0.658 | -0.569 | -0.184 | -0.466 | -0.278 | -0.095 | -0.475 |
| AD | -0.183 | -0.147 | -0.432 | -0.110 | -0.064 | -0.096 | -0.357 |
| CN | 0.057 | -0.958 | -0.622 | -0.531 | -0.798 | -0.103 | 0.150 |
| AD | -1.359 | -1.078 | -0.167 | -0.889 | -0.566 | -0.104 | -0.846 |
| CN | 0.326 | 0.245 | -0.416 | 0.117 | -0.028 | -0.105 | -0.267 |
| AD | 0.084 | -0.073 | 0.010 | -0.023 | -0.113 | -0.108 | -0.034 |
| CN | -2.002 | -1.607 | -0.551 | -0.892 | -0.700 | -0.109 | -0.502 |
| AD | -0.900 | -0.682 | -0.455 | -0.739 | -0.413 | -0.114 | -0.436 |
| MCI | 0.185 | -0.085 | -0.442 | -0.096 | -0.168 | -0.116 | 0.183 |
| MCI | -0.288 | -0.379 | -0.338 | -0.393 | -0.483 | -0.118 | -0.408 |
| CN | 0.843 | 0.419 | 0.124 | 0.898 | 0.607 | -0.119 | 1.024 |
| CN | -0.187 | -0.255 | -0.762 | -0.297 | -0.076 | -0.120 | -0.298 |
| AD | 0.936 | 0.214 | 0.122 | 0.246 | -0.084 | -0.121 | 0.614 |
| MCI | -0.537 | -0.425 | -0.412 | -0.385 | -0.157 | -0.125 | -0.347 |
| MCI | 0.444 | 0.209 | 0.007 | 0.242 | 0.075 | -0.129 | -0.005 |
| MCI | -0.862 | -0.621 | -0.314 | -0.662 | -0.159 | -0.133 | -0.753 |
| AD | 0.229 | 0.281 | 0.090 | 0.031 | 0.015 | -0.135 | -0.270 |
| MCI | -0.276 | -0.331 | -0.873 | -0.648 | -0.704 | -0.137 | -0.452 |
| AD | 0.750 | 0.295 | -0.296 | -0.067 | -0.363 | -0.138 | -0.146 |
| AD | -0.813 | -0.703 | -0.574 | -0.571 | -0.542 | -0.138 | -0.049 |
| MCI | 0.750 | 0.547 | 0.008 | -0.124 | -0.148 | -0.139 | -0.500 |
| MCI | -0.154 | -0.423 | -0.905 | -0.473 | -0.817 | -0.140 | 0.366 |
| MCI | -0.344 | -0.380 | -0.266 | -0.276 | -0.267 | -0.146 | 0.060 |
| CN | -0.208 | -0.104 | -0.450 | -0.140 | 0.026 | -0.154 | -0.441 |
| CN | 0.078 | 0.081 | -0.338 | -0.124 | -0.201 | -0.155 | -0.277 |
| MCI | -1.355 | -0.991 | -0.855 | -1.560 | -1.327 | -0.158 | -1.226 |
| MCI | -0.532 | -0.205 | 0.346 | -0.122 | -0.012 | -0.158 | -0.074 |
| MCI | 0.864 | 0.493 | -0.064 | 0.392 | 0.144 | -0.161 | -0.188 |
| AD | -0.285 | 0.201 | -0.098 | -0.116 | -0.261 | -0.166 | -0.237 |
| CN | 0.489 | 0.321 | 0.175 | 0.798 | 0.380 | -0.168 | 0.632 |
| MCI | -0.795 | -0.557 | -0.159 | -0.583 | -0.213 | -0.168 | -0.336 |
| MCI | -0.323 | -0.295 | -0.783 | -0.446 | -0.192 | -0.174 | -0.649 |
| AD | 0.033 | 0.034 | -0.527 | -0.296 | -0.280 | -0.176 | -0.329 |
| CN | -0.148 | -0.314 | -0.499 | -0.183 | -0.272 | -0.180 | 0.085 |
| AD | 0.766 | 0.590 | -0.124 | 0.782 | 0.177 | -0.181 | 0.192 |
| AD | 0.894 | 0.637 | 0.351 | 0.576 | 0.128 | -0.182 | 0.540 |
| CN | -0.176 | -0.097 | -0.347 | -0.130 | 0.128 | -0.183 | -0.413 |
| MCI | -0.740 | -0.600 | -0.201 | -0.428 | -0.201 | -0.184 | -0.720 |
| MCI | -0.800 | -0.547 | -0.756 | -0.618 | -0.298 | -0.185 | -0.729 |
| CN | 0.469 | 0.182 | 0.048 | 0.192 | -0.024 | -0.187 | 0.267 |
| MCI | 0.096 | 0.026 | -0.206 | -0.162 | -0.183 | -0.190 | -0.057 |
| AD | 0.113 | 0.092 | -0.066 | 0.046 | -0.132 | -0.191 | 0.011 |
| MCI | -1.646 | -1.569 | -1.067 | -1.163 | -0.758 | -0.194 | -0.469 |
| MCI | -0.755 | -0.452 | 0.345 | -0.189 | -0.006 | -0.194 | 0.280 |
| AD | 0.165 | -0.163 | -0.436 | -0.137 | -0.244 | -0.198 | 0.222 |
| MCI | -0.832 | -0.630 | -0.232 | -0.846 | -0.266 | -0.204 | -1.235 |
| CN | -0.781 | -0.662 | -0.742 | -0.746 | -0.316 | -0.205 | -0.823 |
| AD | 0.247 | 0.158 | 0.125 | 0.048 | 0.236 | -0.211 | -0.164 |
| AD | 0.706 | 0.147 | -0.193 | 0.006 | -0.265 | -0.215 | -0.141 |
| AD | -0.134 | -0.044 | -0.500 | -0.157 | -0.101 | -0.217 | -0.366 |
| MCI | -0.415 | -0.683 | -0.686 | -0.373 | -0.301 | -0.225 | 0.012 |
| MCI | -0.277 | -0.080 | -0.008 | -0.147 | -0.253 | -0.227 | -0.280 |
| CN | 0.274 | 0.151 | -0.224 | 0.100 | 0.045 | -0.228 | 0.070 |
| MCI | 0.694 | 0.076 | -0.555 | 0.083 | -0.360 | -0.228 | 0.511 |
| MCI | -0.436 | -0.339 | 0.145 | -0.290 | -0.175 | -0.232 | -0.224 |
| CN | -0.261 | -0.262 | -0.029 | -0.246 | -0.116 | -0.235 | -0.236 |
| CN | -0.879 | -1.019 | -0.712 | -0.716 | -0.482 | -0.244 | -0.473 |
| MCI | -0.520 | -0.608 | -0.173 | -0.335 | -0.279 | -0.245 | -0.084 |
| MCI | 0.211 | 0.207 | 0.161 | 0.222 | 0.015 | -0.254 | 0.339 |
| MCI | -0.467 | -0.319 | -0.305 | -0.679 | -0.190 | -0.255 | -1.464 |
| AD | -0.675 | -0.539 | -0.953 | -0.769 | -0.466 | -0.256 | -1.284 |
| CN | -0.744 | -0.468 | -0.190 | -0.735 | -0.282 | -0.258 | -1.721 |
| MCI | -0.923 | -0.571 | -0.155 | 0.051 | -0.049 | -0.264 | 0.146 |
| CN | 0.077 | 0.056 | 0.009 | -0.088 | 0.009 | -0.267 | -0.198 |
| CN | 0.523 | 0.005 | -0.239 | 0.097 | -0.238 | -0.272 | 0.218 |
| MCI | -0.779 | -0.372 | -0.169 | -0.724 | -0.528 | -0.276 | -0.671 |
| MCI | -0.419 | -0.478 | -0.834 | -0.676 | -0.504 | -0.277 | -0.998 |
| MCI | 0.088 | -0.055 | -0.147 | 0.095 | -0.207 | -0.277 | 0.174 |
| MCI | -0.422 | -0.476 | -0.279 | -0.560 | -0.293 | -0.278 | -0.372 |
| MCI | -0.384 | -0.294 | -0.124 | -0.578 | -0.394 | -0.279 | -0.651 |
| AD | -0.659 | -0.530 | -0.824 | -0.550 | -0.348 | -0.280 | -0.728 |
| MCI | -0.073 | -0.057 | -0.762 | -0.213 | -0.030 | -0.291 | -0.626 |
| MCI | -0.410 | -0.348 | -0.715 | -0.328 | -0.209 | -0.291 | -0.255 |
| CN | 0.439 | 0.030 | -0.238 | -0.067 | -0.238 | -0.295 | -0.524 |
| CN | -0.489 | -0.337 | -0.771 | -0.629 | -0.413 | -0.301 | -0.865 |
| CN | -0.627 | -0.538 | -0.094 | -0.515 | -0.350 | -0.303 | -0.298 |
| AD | -0.716 | -0.721 | -0.921 | -0.944 | -0.558 | -0.304 | -1.013 |
| MCI | -0.759 | -0.856 | -0.328 | -0.790 | -0.763 | -0.308 | -0.580 |
| MCI | -0.191 | -0.356 | -0.121 | -0.215 | -0.245 | -0.313 | 0.127 |
| AD | 0.038 | 0.171 | 0.132 | 0.255 | 0.055 | -0.316 | 0.296 |
| AD | 0.609 | 0.179 | -0.186 | -0.313 | -0.750 | -0.321 | -0.537 |
| AD | -0.629 | -0.573 | -0.948 | -0.940 | -0.829 | -0.321 | -0.579 |
| MCI | 0.255 | 0.063 | -0.145 | 0.088 | -0.130 | -0.327 | 0.200 |
| CN | 0.506 | 0.486 | 0.328 | 0.011 | -0.137 | -0.329 | -0.011 |
| MCI | 0.499 | 0.223 | 0.139 | 0.024 | -0.121 | -0.329 | -0.218 |
| MCI | 0.381 | 0.151 | 0.074 | 0.035 | -0.202 | -0.331 | -0.284 |
| AD | -0.649 | -0.980 | -0.725 | -0.494 | -0.707 | -0.331 | -0.377 |
| MCI | -0.125 | 0.012 | -0.064 | -0.280 | -0.348 | -0.338 | -0.313 |
| MCI | -0.851 | -0.791 | -0.212 | -0.624 | -0.405 | -0.339 | -0.673 |
| MCI | -0.469 | -0.546 | -0.827 | -0.437 | -0.615 | -0.342 | -0.250 |
| CN | -0.458 | -0.302 | -0.166 | -0.439 | -0.419 | -0.344 | -0.710 |
| MCI | 0.427 | -0.107 | -0.391 | 0.297 | 0.061 | -0.360 | 0.669 |
| AD | 0.039 | -0.177 | -0.128 | 0.057 | -0.054 | -0.364 | -0.043 |
| CN | 0.065 | -0.079 | -0.045 | -0.074 | -0.613 | -0.366 | -0.476 |
| CN | -1.773 | -1.224 | -1.421 | -1.382 | -0.967 | -0.367 | -0.899 |
| MCI | -0.875 | -0.841 | -0.583 | -0.955 | -0.780 | -0.375 | -1.085 |
| CN | 0.384 | -0.215 | -0.724 | 0.101 | -0.629 | -0.378 | -0.184 |
| MCI | 0.139 | -0.215 | -0.912 | -0.574 | -0.870 | -0.379 | -1.144 |
| MCI | 0.233 | 0.441 | 0.466 | 0.105 | -0.004 | -0.380 | -0.246 |
| MCI | -0.122 | -0.139 | 0.030 | -0.222 | -0.193 | -0.382 | -0.104 |
| AD | -0.476 | -0.362 | -0.381 | -0.238 | -0.274 | -0.384 | -0.159 |
| MCI | -0.550 | -0.563 | -0.400 | -0.400 | -0.379 | -0.386 | -0.253 |
| MCI | -0.333 | -0.764 | -0.651 | -0.483 | -0.657 | -0.392 | -0.439 |
| CN | -0.425 | -0.282 | -0.615 | -0.456 | -0.196 | -0.399 | -0.802 |
| MCI | 0.607 | 0.131 | -0.110 | 0.113 | -0.396 | -0.403 | 0.464 |
| CN | -1.012 | -0.959 | -0.443 | -0.493 | -0.362 | -0.406 | -0.118 |
| MCI | -1.018 | -0.972 | -1.095 | -0.847 | -0.833 | -0.409 | -0.582 |
| CN | 0.338 | 0.048 | 0.173 | 0.144 | -0.014 | -0.409 | 0.460 |
| MCI | -0.772 | -0.470 | 0.005 | -0.499 | -0.212 | -0.411 | -0.516 |
| MCI | -0.809 | -0.657 | -0.170 | -0.227 | -0.152 | -0.417 | 0.051 |
| MCI | -0.379 | -0.109 | -0.691 | -0.309 | -0.617 | -0.422 | 0.156 |
| MCI | -0.738 | -0.444 | -0.305 | -0.387 | -0.606 | -0.426 | -0.075 |
| MCI | -0.341 | -0.425 | -0.519 | -0.364 | -0.164 | -0.427 | -1.064 |
| AD | -0.965 | -1.069 | -0.494 | -0.459 | -0.570 | -0.429 | 0.220 |
| MCI | -1.172 | -0.677 | -0.438 | -1.018 | -0.767 | -0.432 | -1.054 |
| MCI | -0.622 | -0.559 | -0.233 | -0.695 | -0.460 | -0.438 | -0.985 |
| AD | 0.428 | 0.068 | -0.655 | 0.304 | -0.209 | -0.444 | 0.472 |
| MCI | -0.256 | -0.183 | -0.970 | -0.355 | -0.294 | -0.445 | -0.545 |
| CN | -0.455 | -0.376 | -0.844 | -0.572 | -0.451 | -0.447 | -0.923 |
| AD | -1.592 | -1.684 | -2.094 | -1.812 | -1.495 | -0.450 | -1.024 |
| MCI | -0.018 | -0.091 | -0.302 | -0.450 | -0.501 | -0.456 | -0.611 |
| AD | -0.743 | -0.743 | -0.422 | -0.430 | -0.344 | -0.462 | -0.373 |
| MCI | 0.172 | -0.185 | -0.205 | -0.125 | -0.224 | -0.462 | -0.417 |
| CN | -0.360 | -0.378 | -0.803 | -0.565 | -0.510 | -0.467 | -0.962 |
| MCI | -0.407 | -0.539 | -0.574 | -0.512 | -0.615 | -0.470 | -0.433 |
| AD | -2.024 | -1.753 | -1.434 | -1.441 | -1.099 | -0.473 | -0.973 |
| AD | 0.314 | -0.217 | -0.800 | -0.359 | -0.607 | -0.476 | -0.323 |
| AD | -0.359 | -0.115 | -0.173 | -0.395 | -0.552 | -0.476 | -0.157 |
| MCI | -0.667 | -0.771 | -0.431 | -0.295 | -0.381 | -0.478 | 0.249 |
| MCI | -0.392 | -0.475 | -0.210 | -0.342 | -0.292 | -0.479 | -0.236 |
| AD | -1.531 | -1.549 | -1.013 | -1.167 | -0.956 | -0.485 | -1.108 |
| MCI | -0.570 | -0.248 | -0.404 | -0.511 | -0.756 | -0.490 | -0.369 |
| MCI | 0.312 | -0.022 | -0.233 | -0.039 | -0.179 | -0.490 | -0.215 |
| MCI | 0.080 | -0.259 | -1.008 | -0.535 | -0.832 | -0.493 | -0.522 |
| MCI | -0.245 | -0.375 | -0.336 | -0.279 | -0.704 | -0.496 | -0.265 |
| MCI | 0.006 | -0.158 | 0.028 | -0.209 | -0.313 | -0.499 | -0.099 |
| MCI | -0.957 | -0.858 | -0.157 | -0.422 | -0.289 | -0.500 | -0.390 |
| MCI | -0.008 | 0.020 | -0.574 | -0.272 | -0.384 | -0.504 | -0.831 |
| CN | -0.091 | -0.132 | 0.056 | 0.024 | -0.032 | -0.505 | -0.192 |
| AD | 0.333 | 0.162 | 0.155 | -0.044 | -0.140 | -0.505 | -0.095 |
| CN | -0.602 | -0.487 | -0.408 | -0.602 | -0.403 | -0.510 | -0.925 |
| CN | -0.745 | -0.422 | -0.071 | -0.509 | -0.439 | -0.522 | -0.707 |
| MCI | -0.531 | -0.640 | -0.459 | -0.655 | -0.615 | -0.523 | -1.033 |
| MCI | -0.604 | -0.553 | -0.292 | -0.696 | -0.612 | -0.523 | -0.655 |
| MCI | -0.302 | -0.287 | -0.937 | -0.558 | -0.413 | -0.524 | -1.145 |
| AD | -0.250 | -0.086 | -0.016 | -0.422 | -0.388 | -0.526 | -0.510 |
| MCI | -0.054 | -0.075 | 0.064 | -0.147 | -0.253 | -0.527 | -0.395 |
| AD | -0.284 | -0.435 | -0.294 | -0.421 | -0.305 | -0.527 | -0.514 |
| MCI | 1.020 | 0.635 | 0.030 | 0.532 | -0.178 | -0.533 | 0.100 |
| AD | -0.963 | -0.610 | -0.115 | -0.747 | -0.524 | -0.534 | -0.647 |
| MCI | -0.061 | -0.274 | -0.301 | -0.097 | -0.302 | -0.538 | -0.044 |
| AD | -0.776 | -0.633 | -0.130 | -0.465 | -0.360 | -0.541 | -0.346 |
| CN | -0.389 | -0.415 | -0.498 | -0.538 | -0.272 | -0.542 | -0.903 |
| CN | -0.285 | -0.417 | -0.595 | -0.588 | -0.422 | -0.544 | -0.797 |
| MCI | -0.361 | -0.281 | -0.357 | -0.376 | -0.326 | -0.549 | -0.559 |
| AD | -0.308 | -0.061 | 0.050 | 0.016 | 0.003 | -0.558 | -0.239 |
| MCI | -0.263 | -0.274 | 0.377 | -0.041 | 0.048 | -0.562 | 0.193 |
| CN | -0.737 | -0.788 | -1.020 | -1.252 | -1.090 | -0.562 | -1.151 |
| AD | -0.389 | -0.198 | -0.257 | -0.249 | -0.361 | -0.565 | -0.184 |
| MCI | 0.264 | 0.202 | -0.150 | -0.197 | -0.609 | -0.567 | -0.376 |
| AD | 0.307 | 0.281 | 0.267 | -0.029 | -0.154 | -0.569 | -0.471 |
| MCI | -0.223 | -0.343 | -1.001 | -0.528 | -0.456 | -0.569 | -0.965 |
| MCI | -0.365 | -0.394 | -0.438 | -0.433 | -0.521 | -0.569 | -0.250 |
| AD | -0.916 | -0.650 | -0.237 | -0.585 | -0.504 | -0.573 | -0.523 |
| MCI | -0.773 | -0.629 | -0.363 | -0.780 | -0.826 | -0.574 | -0.250 |
| MCI | -0.751 | -0.769 | -0.208 | -0.154 | -0.252 | -0.576 | -0.005 |
| CN | -1.050 | -1.021 | -1.212 | -1.119 | -1.019 | -0.578 | -0.875 |
| MCI | -0.847 | -0.895 | -0.599 | -0.944 | -0.672 | -0.583 | -1.776 |
| MCI | -0.135 | -0.332 | 0.301 | -0.062 | -0.066 | -0.585 | -0.256 |
| MCI | -1.886 | -1.302 | -0.474 | -0.649 | -0.488 | -0.590 | -0.400 |
| MCI | -1.024 | -0.842 | -0.436 | -1.029 | -0.628 | -0.591 | -1.252 |
| CN | -0.594 | -0.525 | -0.394 | -0.633 | -0.599 | -0.592 | -0.467 |
| AD | -0.888 | -0.707 | -0.143 | -0.244 | -0.112 | -0.597 | -0.113 |
| MCI | 0.963 | 0.348 | 0.035 | 0.819 | -0.014 | -0.598 | 0.682 |
| MCI | -1.130 | -0.890 | -0.538 | -1.187 | -0.897 | -0.599 | -0.846 |
| MCI | -0.912 | -1.009 | -1.263 | -1.248 | -1.137 | -0.606 | -0.788 |
| MCI | -0.716 | -0.345 | -0.077 | -0.555 | -0.382 | -0.626 | -0.756 |
| CN | -1.345 | -1.334 | -0.721 | -0.655 | -0.769 | -0.630 | -0.703 |
| CN | 0.162 | 0.485 | 0.159 | -0.041 | -0.304 | -0.632 | -0.473 |
| MCI | 0.868 | 0.228 | -0.053 | 0.415 | -0.032 | -0.635 | 0.218 |
| AD | -1.824 | -1.607 | -0.780 | -0.789 | -0.588 | -0.636 | -0.747 |
| CN | -0.141 | -0.388 | -0.521 | -0.402 | -0.614 | -0.644 | -0.647 |
| MCI | -0.410 | -0.297 | 0.028 | -0.460 | -0.375 | -0.660 | -0.498 |
| MCI | -0.099 | -0.116 | -0.433 | -0.435 | -0.728 | -0.661 | -0.649 |
| MCI | -0.511 | -0.501 | -1.216 | -0.673 | -0.899 | -0.664 | -0.584 |
| AD | -0.098 | -0.071 | 0.071 | -0.244 | -0.282 | -0.670 | -0.586 |
| MCI | 0.350 | 0.105 | -0.157 | 0.064 | -0.367 | -0.681 | 0.192 |
| MCI | -0.674 | -0.796 | -0.054 | -0.387 | -0.377 | -0.688 | -0.275 |
| MCI | -0.474 | -0.382 | -0.295 | -0.455 | -0.690 | -0.689 | -0.408 |
| AD | -0.390 | -0.886 | -1.740 | -1.162 | -1.639 | -0.697 | -0.194 |
| MCI | -1.355 | -0.913 | -0.758 | -1.028 | -0.738 | -0.700 | -0.342 |
| CN | -0.220 | -0.431 | -0.532 | -0.418 | -0.586 | -0.702 | -0.783 |
| MCI | -0.249 | -0.283 | -0.844 | -0.279 | -0.635 | -0.709 | -0.584 |
| MCI | -0.873 | -0.867 | 0.168 | -0.308 | -0.343 | -0.713 | -0.233 |
| AD | -1.437 | -1.569 | -1.642 | -2.034 | -1.570 | -0.721 | -0.893 |
| CN | -0.519 | -0.337 | -0.205 | -0.350 | -0.401 | -0.730 | -0.317 |
| AD | -1.092 | -0.968 | 0.121 | -0.531 | -0.407 | -0.746 | -0.592 |
| AD | 0.159 | 0.019 | -0.238 | -0.270 | -0.493 | -0.747 | -0.656 |
| AD | -0.046 | -0.018 | -0.464 | -0.138 | -0.222 | -0.748 | -0.386 |
| CN | -0.346 | -0.266 | -1.048 | -0.421 | -0.826 | -0.754 | -0.722 |
| CN | -0.530 | -0.322 | -0.266 | -0.552 | -0.662 | -0.760 | -1.010 |
| CN | 0.734 | 0.181 | -0.433 | 0.162 | -0.401 | -0.768 | -0.613 |
| CN | 0.082 | -0.199 | -0.047 | -0.282 | -0.389 | -0.769 | -0.473 |
| MCI | -0.293 | -0.630 | -0.524 | -0.488 | -0.509 | -0.771 | -0.964 |
| MCI | -0.679 | -0.862 | -0.543 | -0.662 | -0.696 | -0.774 | -0.467 |
| MCI | -0.416 | -0.355 | -0.235 | -0.618 | -0.737 | -0.775 | -0.816 |
| AD | -0.434 | -0.265 | -0.170 | -0.481 | -0.622 | -0.783 | -0.787 |
| AD | -0.196 | -0.371 | -0.435 | -0.056 | -0.323 | -0.784 | -0.253 |
| MCI | -1.403 | -1.199 | -1.159 | -0.919 | -0.903 | -0.789 | -0.744 |
| MCI | -0.712 | -1.037 | -0.451 | -0.235 | -0.377 | -0.791 | -0.038 |
| AD | -0.179 | -0.631 | -0.506 | -0.036 | -0.288 | -0.791 | -0.339 |
| MCI | -1.150 | -1.017 | 0.119 | -0.390 | -0.381 | -0.794 | -0.435 |
| CN | 0.122 | -0.135 | -0.138 | -0.185 | -0.398 | -0.796 | -0.355 |
| AD | -0.208 | -0.136 | 0.074 | -0.319 | -0.409 | -0.797 | -0.486 |
| AD | -0.160 | -0.326 | -1.028 | -0.659 | -1.073 | -0.797 | -0.677 |
| MCI | -1.127 | -1.158 | -0.548 | -0.723 | -0.710 | -0.813 | -0.271 |
| MCI | -0.275 | -0.127 | -1.125 | -0.014 | -0.784 | -0.814 | -0.422 |
| AD | -0.795 | -0.702 | -0.696 | -1.097 | -0.971 | -0.816 | -1.064 |
| CN | -0.090 | 0.031 | 0.225 | -0.020 | -0.131 | -0.827 | 0.060 |
| MCI | 0.328 | -0.135 | -0.609 | -0.188 | -0.649 | -0.842 | -0.454 |
| MCI | -0.120 | -0.371 | -0.138 | -0.621 | -0.446 | -0.845 | -1.516 |
| MCI | -1.408 | -1.859 | -1.382 | -1.033 | -1.093 | -0.851 | -1.182 |
| CN | 0.562 | -0.049 | -0.813 | -0.107 | -0.917 | -0.852 | -0.287 |
| AD | -0.998 | -0.841 | -0.210 | -0.850 | -0.637 | -0.860 | -0.493 |
| MCI | -1.036 | -1.411 | -2.030 | -1.634 | -1.882 | -0.861 | -0.674 |
| MCI | -0.860 | -1.067 | -1.486 | -1.385 | -1.331 | -0.866 | -1.227 |
| CN | -0.906 | -1.238 | -0.957 | -0.923 | -0.805 | -0.873 | -1.356 |
| CN | -0.538 | -0.942 | -1.002 | -0.654 | -0.944 | -0.874 | -0.747 |
| MCI | -0.851 | -0.867 | -0.691 | -0.867 | -0.630 | -0.874 | -1.392 |
| MCI | -1.777 | -1.817 | -0.947 | -1.159 | -1.080 | -0.875 | -1.168 |
| AD | -0.636 | -0.552 | -0.158 | -0.692 | -0.573 | -0.883 | -0.826 |
| MCI | -0.595 | -0.707 | -0.583 | -0.688 | -0.585 | -0.885 | -0.777 |
| MCI | -0.811 | -1.150 | -0.880 | -0.758 | -0.920 | -0.888 | -0.597 |
| CN | -0.843 | 0.033 | 0.342 | -0.892 | -0.631 | -0.893 | -0.731 |
| MCI | -0.135 | -0.212 | -0.109 | -0.321 | -0.518 | -0.894 | -0.669 |
| MCI | -1.224 | -1.169 | -0.885 | -0.864 | -1.104 | -0.899 | -1.062 |
| MCI | -1.568 | -1.211 | -0.777 | -0.805 | -0.733 | -0.900 | -0.872 |
| MCI | -0.983 | -1.010 | -0.652 | -1.240 | -0.847 | -0.906 | -1.265 |
| MCI | 0.292 | -0.345 | -0.547 | -0.154 | -0.492 | -0.911 | -0.283 |
| CN | 0.405 | 0.427 | 0.357 | 0.441 | 0.060 | -0.919 | 0.143 |
| MCI | -0.704 | -0.653 | -0.445 | -0.852 | -0.956 | -0.922 | -0.913 |
| MCI | -0.403 | -0.251 | -0.077 | -0.390 | -0.445 | -0.928 | -0.508 |
| AD | -0.276 | -0.293 | -0.134 | -0.561 | -0.663 | -0.945 | -0.617 |
| MCI | -0.595 | -0.376 | -0.444 | -0.370 | -0.465 | -0.948 | -0.208 |
| MCI | -1.230 | -1.492 | -0.963 | -1.140 | -0.995 | -0.950 | -1.001 |
| MCI | -0.492 | -0.623 | -0.982 | -0.916 | -1.110 | -0.956 | -0.575 |
| CN | -0.737 | -1.010 | -0.580 | -0.275 | -0.335 | -0.960 | -0.302 |
| MCI | -0.706 | -0.803 | -1.853 | -1.613 | -1.728 | -0.961 | -1.584 |
| MCI | -1.203 | -1.281 | -1.000 | -1.286 | -1.046 | -0.974 | -1.710 |
| CN | -0.893 | -1.287 | -0.933 | -0.762 | -0.836 | -0.975 | -0.950 |
| MCI | -1.051 | -0.918 | -1.101 | -1.042 | -1.191 | -0.978 | -0.866 |
| CN | -0.455 | -0.743 | -0.406 | -0.587 | -0.726 | -0.991 | -0.265 |
| MCI | -0.531 | -0.030 | -0.028 | -0.415 | -0.386 | -0.998 | -0.328 |
| MCI | -0.030 | -0.247 | -0.712 | -0.679 | -0.774 | -1.000 | -0.823 |
| MCI | -0.800 | -0.684 | -0.239 | -0.685 | -0.601 | -1.005 | -1.010 |
| MCI | -0.354 | -0.623 | -1.063 | -0.716 | -1.136 | -1.006 | -0.243 |
| CN | -1.411 | -1.578 | -0.927 | -1.341 | -1.461 | -1.011 | -1.433 |
| AD | -0.283 | -0.523 | -0.206 | -0.414 | -0.728 | -1.015 | -0.875 |
| MCI | -1.514 | -1.435 | -0.819 | -0.934 | -0.999 | -1.018 | -0.718 |
| MCI | -0.859 | -0.734 | -0.544 | -0.978 | -1.028 | -1.025 | -0.422 |
| MCI | -0.454 | -0.422 | -0.166 | -0.555 | -0.629 | -1.037 | -0.851 |
| MCI | -0.453 | -0.954 | -0.531 | -0.552 | -0.998 | -1.039 | -0.139 |
| MCI | -1.479 | -1.530 | -0.714 | -0.696 | -0.773 | -1.042 | -0.092 |
| AD | -0.278 | -0.674 | -1.145 | -0.631 | -1.090 | -1.048 | -0.944 |
| AD | -1.030 | -0.910 | -1.053 | -1.048 | -1.084 | -1.054 | -1.189 |
| AD | -1.784 | -1.860 | -1.312 | -1.609 | -1.345 | -1.056 | -1.742 |
| CN | -0.850 | -1.135 | -0.871 | -0.869 | -0.987 | -1.060 | -0.808 |
| AD | -1.574 | -1.275 | -0.760 | -1.504 | -1.117 | -1.075 | -1.091 |
| AD | -2.144 | -2.185 | -0.777 | -1.415 | -0.981 | -1.081 | -1.429 |
| CN | 0.758 | -0.011 | -1.052 | -0.165 | -1.233 | -1.089 | -0.358 |
| MCI | -0.254 | -0.355 | -0.319 | -0.207 | -0.602 | -1.091 | -0.642 |
| MCI | -1.575 | -1.414 | -0.851 | -1.032 | -1.108 | -1.112 | -1.455 |
| CN | -1.015 | -0.794 | -0.332 | -0.789 | -0.639 | -1.115 | -0.707 |
| MCI | -1.057 | -0.520 | -0.232 | -1.217 | -0.953 | -1.126 | -1.523 |
| AD | -1.154 | -0.912 | -0.278 | -1.038 | -0.878 | -1.139 | -0.655 |
| AD | -0.738 | -0.734 | -0.371 | -0.645 | -0.771 | -1.143 | -0.754 |
| MCI | -0.041 | -0.090 | -0.117 | -0.346 | -0.679 | -1.168 | -0.834 |
| MCI | -0.987 | -1.371 | -0.869 | -0.840 | -1.059 | -1.186 | -1.148 |
| AD | -0.937 | -0.719 | -0.366 | -1.000 | -0.716 | -1.188 | -1.301 |
| MCI | -0.250 | -0.941 | -1.093 | -0.601 | -1.260 | -1.195 | -0.829 |
| MCI | -1.014 | -1.201 | -1.028 | -0.753 | -0.870 | -1.196 | -0.778 |
| AD | -0.757 | -0.557 | -0.132 | -0.686 | -0.639 | -1.200 | -0.899 |
| CN | -1.216 | -1.311 | -0.744 | -0.819 | -1.003 | -1.204 | -1.374 |
| CN | -1.149 | -1.157 | -1.679 | -1.484 | -1.829 | -1.216 | -0.994 |
| MCI | -0.695 | -0.594 | -0.227 | -0.727 | -0.871 | -1.227 | -1.163 |
| CN | -0.907 | -1.137 | -0.967 | -0.961 | -1.152 | -1.285 | -1.083 |
| CN | -0.590 | -0.670 | -0.855 | -0.753 | -0.777 | -1.330 | -1.101 |
| MCI | -0.739 | -0.924 | -1.123 | -0.983 | -1.096 | -1.332 | -0.866 |
| MCI | -0.934 | -1.019 | -0.921 | -1.246 | -0.946 | -1.345 | -2.094 |
| AD | -2.408 | -1.881 | -1.038 | -1.782 | -1.489 | -1.355 | -1.051 |
| AD | -1.376 | -1.473 | -0.986 | -1.309 | -1.263 | -1.364 | -1.219 |
| AD | -0.919 | -0.605 | -0.632 | -0.696 | -0.896 | -1.387 | -0.943 |
| AD | -1.238 | -1.257 | -1.838 | -1.943 | -1.848 | -1.387 | -1.899 |
| MCI | -0.620 | -0.998 | -0.855 | -0.952 | -1.197 | -1.416 | -1.128 |
| MCI | -0.288 | -0.601 | -0.685 | -0.802 | -1.186 | -1.459 | -1.151 |
| MCI | -0.598 | -0.860 | -0.968 | -0.739 | -1.084 | -1.469 | -1.300 |
| MCI | -1.340 | -0.941 | -0.236 | -0.882 | -0.781 | -1.471 | -1.094 |
| CN | -0.527 | -1.022 | -0.523 | -0.766 | -1.201 | -1.472 | -1.033 |
| AD | -1.134 | -1.312 | -1.691 | -1.651 | -1.883 | -1.482 | -1.325 |
| AD | -0.809 | -1.300 | -1.231 | -1.051 | -1.535 | -1.518 | -1.314 |
| MCI | -1.820 | -1.905 | -2.357 | -2.288 | -2.706 | -1.523 | -1.388 |
| CN | -1.346 | -1.425 | -0.951 | -1.076 | -1.201 | -1.526 | -1.133 |
| CN | -0.342 | -0.803 | -0.594 | -0.456 | -0.901 | -1.531 | -0.914 |
| CN | -0.404 | -0.681 | -1.794 | -0.917 | -1.716 | -1.568 | -1.090 |
| AD | -0.287 | -0.796 | -0.828 | -0.939 | -1.596 | -1.580 | -0.892 |
| CN | -0.690 | -0.892 | -0.428 | -0.824 | -0.905 | -1.595 | -0.933 |
| AD | -1.579 | -1.760 | -0.970 | -1.049 | -1.114 | -1.600 | -1.484 |
| CN | -1.288 | -1.316 | -0.690 | -1.270 | -1.218 | -1.607 | -1.383 |
| MCI | -0.450 | -1.070 | -1.044 | -0.926 | -1.281 | -1.644 | -1.279 |
| CN | -0.915 | -1.432 | -0.891 | -0.736 | -1.075 | -1.662 | -1.050 |
| AD | -0.415 | -0.833 | -0.814 | -0.685 | -1.215 | -1.664 | -1.083 |
| AD | -0.310 | -1.107 | -0.821 | -0.543 | -0.880 | -1.665 | -0.468 |
| MCI | -0.504 | -1.011 | -0.989 | -1.011 | -1.533 | -1.670 | -1.258 |
| MCI | -1.074 | -1.528 | -1.438 | -1.180 | -1.694 | -1.674 | -0.952 |
| CN | -1.422 | -1.415 | -2.199 | -2.172 | -2.720 | -1.676 | -1.611 |
| MCI | -1.296 | -1.568 | -0.859 | -0.581 | -0.766 | -1.798 | -0.592 |
| MCI | -1.003 | -1.773 | -1.441 | -1.207 | -1.637 | -1.816 | -1.667 |
| AD | -0.763 | -0.851 | -0.989 | -0.835 | -1.277 | -1.848 | -1.127 |
| MCI | -0.430 | -1.159 | -1.089 | -0.980 | -1.904 | -1.944 | -1.457 |
| AD | -0.584 | -0.726 | -1.039 | -0.891 | -0.962 | -2.081 | -1.428 |
| MCI | -2.173 | -2.556 | -2.848 | -2.765 | -2.778 | -2.175 | -2.787 |
| CN | -1.559 | -1.887 | -1.162 | -1.177 | -1.409 | -2.307 | -1.474 |
| CN | -1.547 | -2.078 | -1.036 | -1.160 | -1.590 | -2.752 | -1.799 |
| AD | -0.741 | -1.402 | -1.249 | -0.767 | -1.391 | -2.835 | -1.551 |
| MCI | -3.099 | -3.004 | -2.720 | -2.724 | -2.711 | -2.837 | -2.570 |
| MCI | -1.942 | -2.122 | -1.934 | -2.067 | -2.526 | -3.082 | -2.743 |
